# Biochemical, structural, and single-molecule characterization of LIG1 active site mutants demonstrate role of F635 and F872 residues for faithful ligation

**DOI:** 10.1101/2024.11.07.622578

**Authors:** Mustafa Kalaycioğlu, Mitchell Gulkis, Kar Men Lee, Surajit Chatterjee, Kanal Balu, Qun Tang, Jacob E. Ratcliffe, Erick Castro, Danah Almahdor, Melike Çağlayan

## Abstract

Human DNA ligase 1 (LIG1) finalizes DNA repair pathways by an ultimate ligation step and discriminates against nicks containing unusual ends, yet the contribution of the conserved active site residues for faithful end joining remains unknown. Here, using biochemistry, X-ray crystallography, and single-molecule approaches, we comprehensively characterized LIG1 mutants carrying Ala(A) and Leu(L) substitutions at the active site residues Phe(F)635 and Phe(F)872. Our results showed an abolished ligation of nick DNA substrates with all 12 non-canonical mismatches, while the mutagenic nick sealing of oxidatively damaged ends by wild-type enzyme is significantly reduced by F635A/L and F872A/L substitutions. Furthermore, sugar discrimination against a single ribonucleotide at 3’- or 5’-end of nick DNA is distinctly affected depending on architecture of 3’-terminus:template base pairing. Finally, our LIG1 structures demonstrated the importance of DNA end alignment governed by the distance to nick site through F635 and F872 residues, and single-molecule measurements showed similar nick DNA binding modes for LIG1 wild-type and active site mutants in real-time. Overall, our study provides a mechanistic insight into the mechanism by which conserved F635 and F872 residues contribute to ligation efficiency of nick repair intermediates that mimic DNA polymerase-mediated mismatch, damaged, or ribonucleotide insertion products and how LIG1 ensures faithful end joining at the final step of DNA repair to maintain genome integrity.

## Introduction

Genomic DNA is susceptible to damage from numerous sources, including endogenous and environmental factors (1). One potentially toxic form of DNA damage is a single-strand break, otherwise known as a nick (2). DNA nicks are intermediates that can be generated naturally throughout almost all aspects of DNA metabolism, such as replication, repair, and recombination, but they can also be generated by exogenous factors such as ionizing radiation and environmental factors (3). If left unrepaired and allowed to persist, these strand-breaks could block replication fork progression at nick sites and potentially lead to lethal double-strand breaks (4). Therefore, a thorough dissection of the mechanism by which nicks are faithfully repaired is crucial to understand how overall genome stability is maintained.

DNA ligases catalyze the formation of a phosphodiester bond at single-strand breaks by the in-line nucleophilic attack of the 3’-hydroxyl (OH) on the 5’-phosphate (PO_4_) groups (5,6). Previously solved crystal structures of ATP-dependent DNA ligases from other sources such as *Chlorella virus* DNA ligase, T4 DNA ligase, *Mycobacterium tuberculosis* LigD, and *Pyrococcus furiosus* extensively contributed to understand the mechanism of nick ligation (7–22). The catalytic activity is largely attributed to residues in the conserved adenylation domain (AdD) and oligonucleotide binding fold (OB-fold) domains, which are referred to as the catalytic core (7–11). Eukaryotic DNA ligases utilize a large N-terminal DNA-binding domain (DBD) to aid in binding their nick substrate and the DBD interacts with the C-terminal OB-fold domain, helping to bridge the two domains and encircle the nick DNA substrate (12–16). Through the catalytic core, ATP-dependent DNA ligases catalyze three-step consecutive ligation reaction: the formation of the DNA ligase-adenylate intermediate (DNA ligase-AMP) in step 1, subsequent transfer of AMP moiety to the 5’-PO_4_ end of the nick (DNA-AMP) in step 2, and a final phosphodiester bond formation coupled to AMP release in step 3 (10–20). DNA ligase 1 (LIG1) is the main replicative ligase, responsible for joining Okazaki fragments during each round of replication and finalizes DNA repair pathways by last nick sealing step after DNA polymerase-mediated nucleotide incorporation (23–25). The first X-ray structure of LIG1 demonstrated the conserved mechanism of human DNA ligases, by which the ligase encircles the nick with all three domains making contacts with the DNA and uses active site residues residing in the AdD and OBD domains to specifically bind and catalyze nick sealing (23). Furthermore, it was shown that the phenylalanine (F) residues, F635 and F872, residing in the AdD and OBD domains, respectively, push against the 3’- and 5’-ends of the nick, respectively (Supplementary Scheme 1A-B). These residues were also shown to insert into the minor groove and enforce underwinding of several base pairs upstream of the nick (23). Additionally, it has been reported in the *Chlorella* virus ligase that these two conserved active site residues are essential for nick sealing both *in vitro* and *in vivo* (12). Indeed, the amino acid sequence alignment of human, virus, bacteria, and yeast DNA ligases show that F635 is conserved back to *Saccharomyces cerevisiae* and F872 is conserved even further back, to *Escherichia coli* (Supplementary Scheme 1C-D).

DNA polymerases and DNA ligases finalize almost all excision repair pathways at downstream steps involving a nucleotide insertion into a gap during DNA synthesis, which creates a nick for subsequent ligation (26). The insertion of a correct nucleotide by DNA polymerase creates a nick substrate with a canonical 3’-OH and 5’-PO_4_ ends to be sealed efficiently by DNA ligase at the final step. Our previous studies demonstrated that mismatch or damaged nucleotide (8-oxodGTP) incorporation by DNA polymerase confounds the following ligation step, leading to an interruption in the coordinated repair pathway and formation of ligation failure products (27–31).

Genomic ribonucleotides are a common type of DNA damage and can lead genomic instability by causing local transition from B- to A-form DNA, altering structure of the DNA duplex, increased propensity to single strand breaks and replicative stress (32). Ribonucleotides (rNTPs) are commonly incorporated during replication by DNA polymerase ε and δ and can be removed by RNase H2-mediated ribonucleotide excision repair (RER) or Top1-mediated ribonucleotide removal in the absence of RER (33). We recently demonstrated that LIG1 lacks a discrimination against a single ribonucleotide at 3’-end of nick DNA which mimic rNTP insertion products of repair and replication DNA polymerases (34). On the other hand, we uncover the mechanism by which LIG1 excludes nick sealing in the presence of a single ribonucleotide at 5’-end of nick DNA that could be formed as intermediates during RER (35). In the present study, we further explored the ligase strategies for faithful nick sealing and investigated the roles of LIG1 active site residues Phe(F)635 and Phe(F)872 on the efficiency of ligation and nick DNA binding at biochemical, structural, and single-molecule levels. For this purpose, we created alanine (A) and leucine (L) substitutions and generated four single mutants F635A, F635L, F872A, and F872L to characterize the impact of these mutations on the ligation efficiency of nick DNA substrates containing canonical, mismatched, damaged, and ribonucleotide-containing ends *in vitro*. We comprehensively investigated all 12 possible non-canonical mismatches and all 4 nick substrates containing 8-oxoG at the 3’-end that mimic base substitution errors and damaged nucleotide insertion products of DNA polymerases, respectively. Furthermore, we questioned the impact of mutations at LIG1 F635 and F872 active side residues on sugar discrimination against nick DNA substrates containing a single nucleotide at either 3’- or 5’-end.

Our findings demonstrated completely abolished nick sealing of all 12 non-canonical mismatches by LIG1 F635A, F635L, F872A, and F872L mutants. The substrate specificity of all 4 mutants for 3’-damaged end demonstrated reduced ligation efficiency depending on the Hoogsteen *versus* Watson-Crick base pairing feature of 8-oxoG. We also showed less efficient nick sealing in the presence of 3’-ribonucleotide while only F872A showed increased ligation for nicks containing 5’-ribonucleotide when compared to wild-type enzyme. Furthermore, X-ray structures of LIG1 active site mutants demonstrated the importance of F635 and F872 residues for proper alignment of 3’- and 5’-ends, respectively, and F to A substitutions resulted in drastic differences in the distances relative to nucleotides around nick site. Finally, single-molecule characterization of nick DNA binding by LIG1 F635A and F872A mutants in real-time showed similar modes as wild-type enzyme. Overall, our study provides a mechanistic insight into how LIG1 active site residues F635 and F872 contribute to maintain ligase fidelity while engaging with non-canonical repair intermediates and how they play role for surveilling DNA ends containing a mismatched or damaged base and “wrong” sugar at the last step of DNA repair and replication.

## Results

### Impact of LIG1 F635 and F872 active site mutations on the ligation efficiency of nick DNA substrates containing mismatches

We first comprehensively investigated the ligation efficiency of LIG1 active site mutants carrying Phe(F) to Ala (A) or Leu (L) substitutions at F635 and F872 (F635A, F635L, F872A, F872L) for the nick DNA substrates with all possible 12 non-canonical mismatches (Supplementary Scheme 2A).

As we previously reported (28), our results showed that LIG1 wild-type can indiscriminately ligate nick DNA substrates containing 3’-mismatches (Figure 1 and Supplementary Figure 2). For example, higher ligation efficiency was observed for 3’-mismatches: 3’-G:T, 3’-C:T, and 3’-T:C. In contrast, nick DNA substrates containing 3’-G:A and 3’-A:G mismatches showed the lowest efficiency with less than 10 % of ligation products. In the presence of the F635A mutation, the ligation efficiency drastically decreased with less than 10 % of ligation products for all 12 non-canonical mismatches (Figure 2 and Supplementary Figure 3). Similarly, our results showed that LIG1 F635L mutant is not capable of nick sealing, except the 3’-T:C and 3’-C:T mismatches, which show slightly higher amount of ligation products than those of LIG1 F635A mutant (Figure 3 and Supplementary Figure 4). Additionally, we obtained a decrease in the ligation products for all 12 non-canonical mismatches by both F872A (Figure 4 and Supplementary Figure 5) and F872L (Figure 5 and Supplementary Figure 6) mutants. Overall, our results demonstrated the crucial role of LIG1 active site residues, F635 and F872, for the efficient ligation of mismatched termini. The nick sealing efficiency by all four active site mutants for nick DNA substrates containing canonical ends (3’-A:T, 3’-T:A, 3’-G:C, 3’-C:G) was similar to LIG1 wild-type with slight differences at earlier time points (Supplementary Figures 7-11).

**Figure 1.**
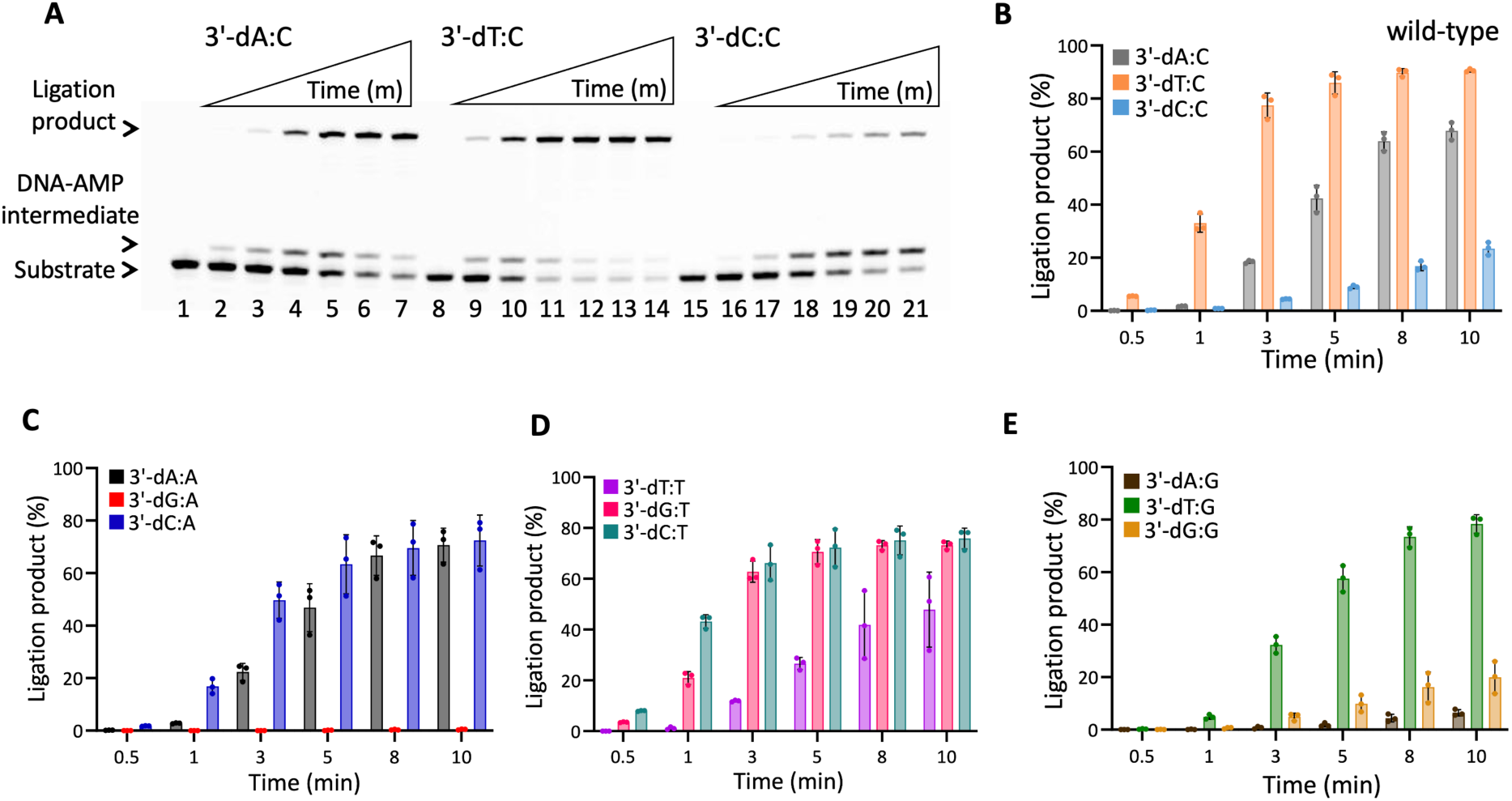
DNA substrate specificity of LIG1 wild-type for all 12 mismatches. (**A**) Lanes 1, 8, and 15 are the negative enzyme controls for nick substrates with 3’-dA:C, 3’-dT:C, and 3’-dC:C, respectively. Lanes 2-7, 9-14, and 16-21 are the ligation reaction products and correspond to time points of 0.5, 1, 3, 5, 8, and 10 min. (**B-E**) Graphs show time-dependent changes in the amount of ligation products. The data represent the average from three independent experiments ± SD.

**Figure 2.**
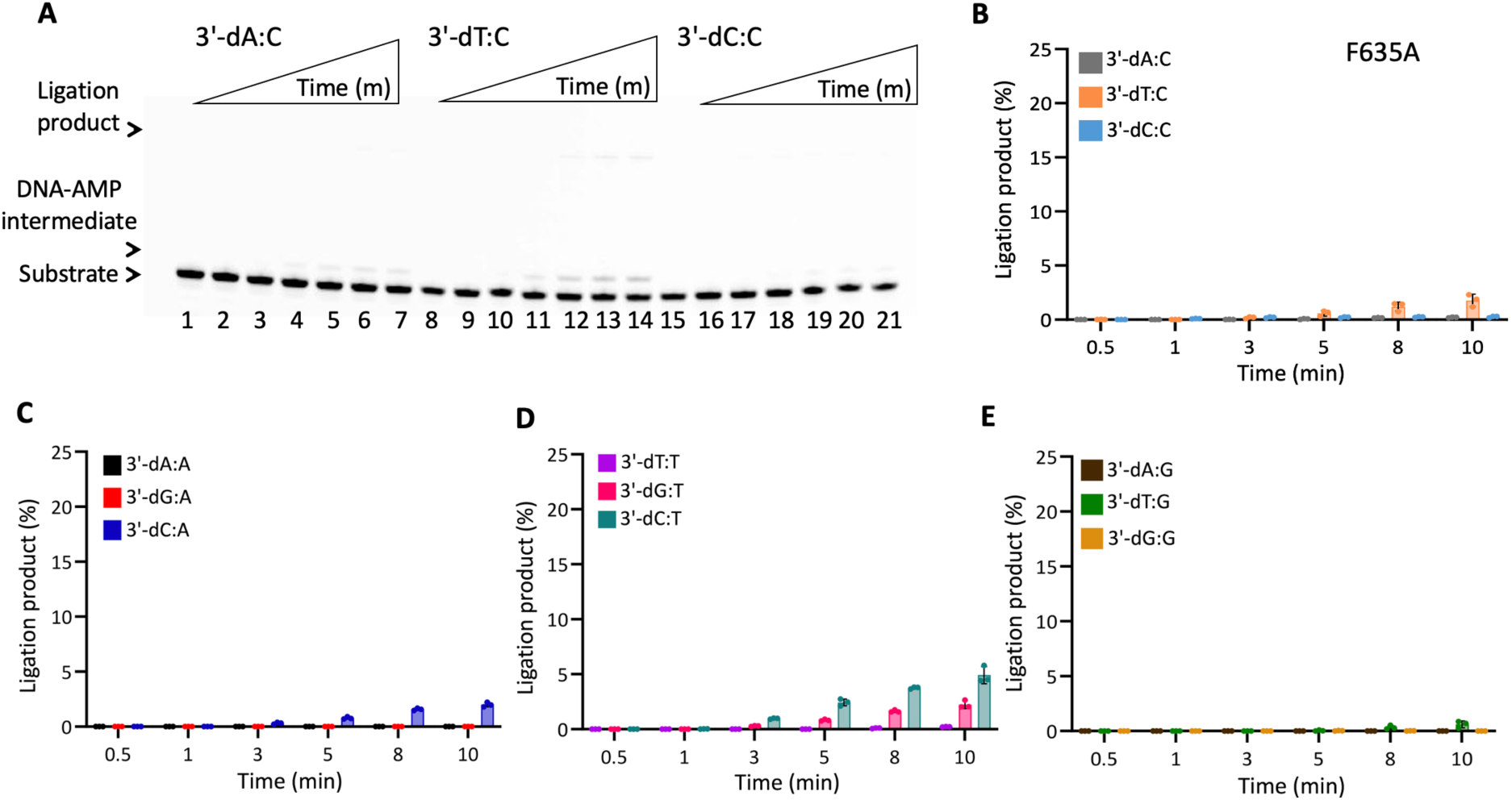
DNA substrate specificity of LIG1 F635A for all 12 mismatches. (**A**) Lanes 1, 8, and 15 are the negative enzyme controls for nick substrates with 3’-dA:C, 3’-dT:C, and 3’-dC:C, respectively. Lanes 2-7, 9-14, and 16-21 are the ligation reaction products and correspond to time points of 0.5, 1, 3, 5, 8, and 10 min. (**B-E**) Graphs show time-dependent changes in the amount of ligation products. The data represent the average from three independent experiments ± SD.

**Figure 3.**
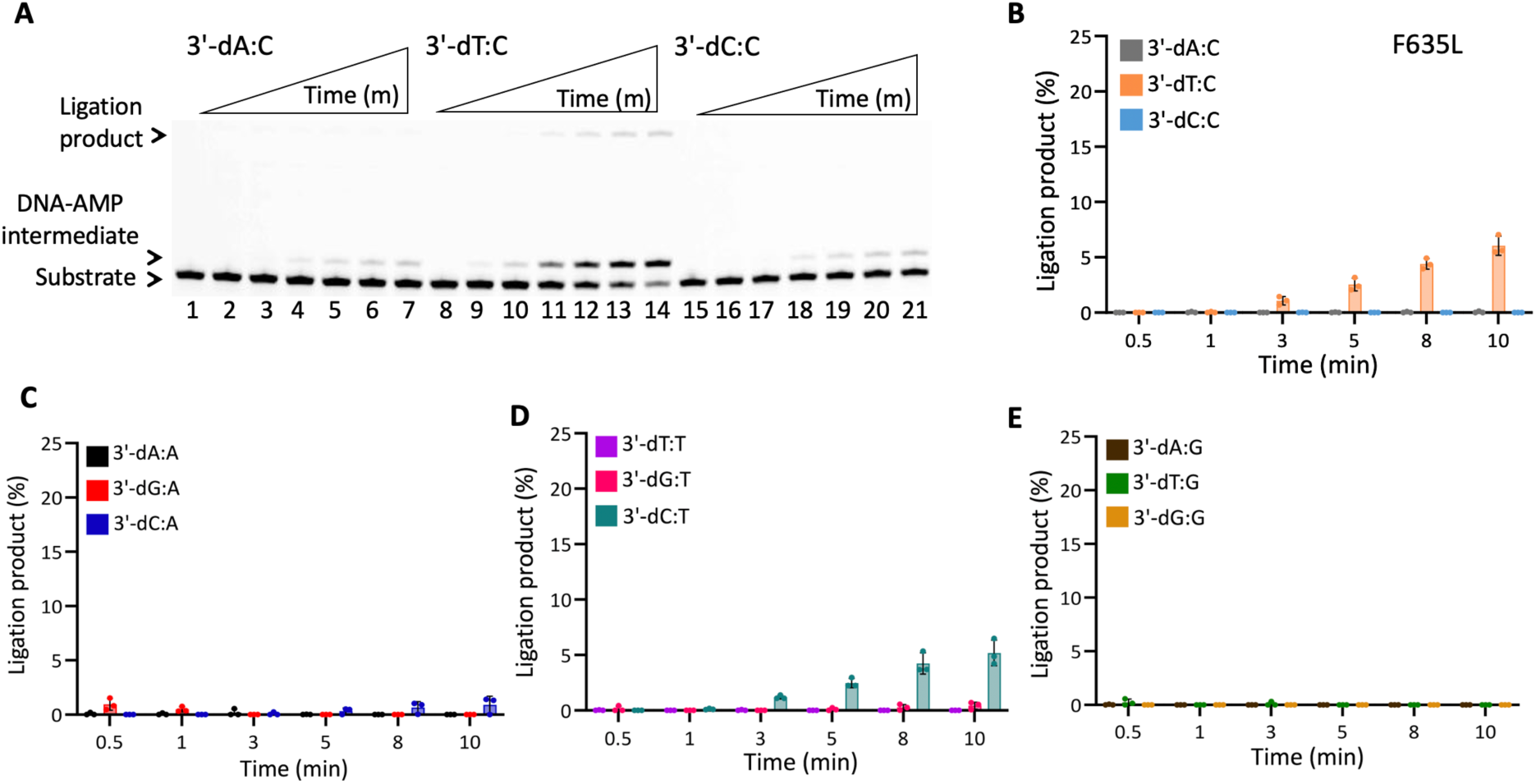
DNA substrate specificity of LIG1 F635L for all 12 mismatches. (**A**) Lanes 1, 8, and 15 are the negative enzyme controls for nick substrates with 3’-dA:C, 3’-dT:C, and 3’-dC:C, respectively. Lanes 2-7, 9-14, and 16-21 are the ligation reaction products and correspond to time points of 0.5, 1, 3, 5, 8, and 10 min. (**B-E**) Graphs show time-dependent changes in the amount of ligation products. The data represent the average from three independent experiments ± SD.

**Figure 4.**
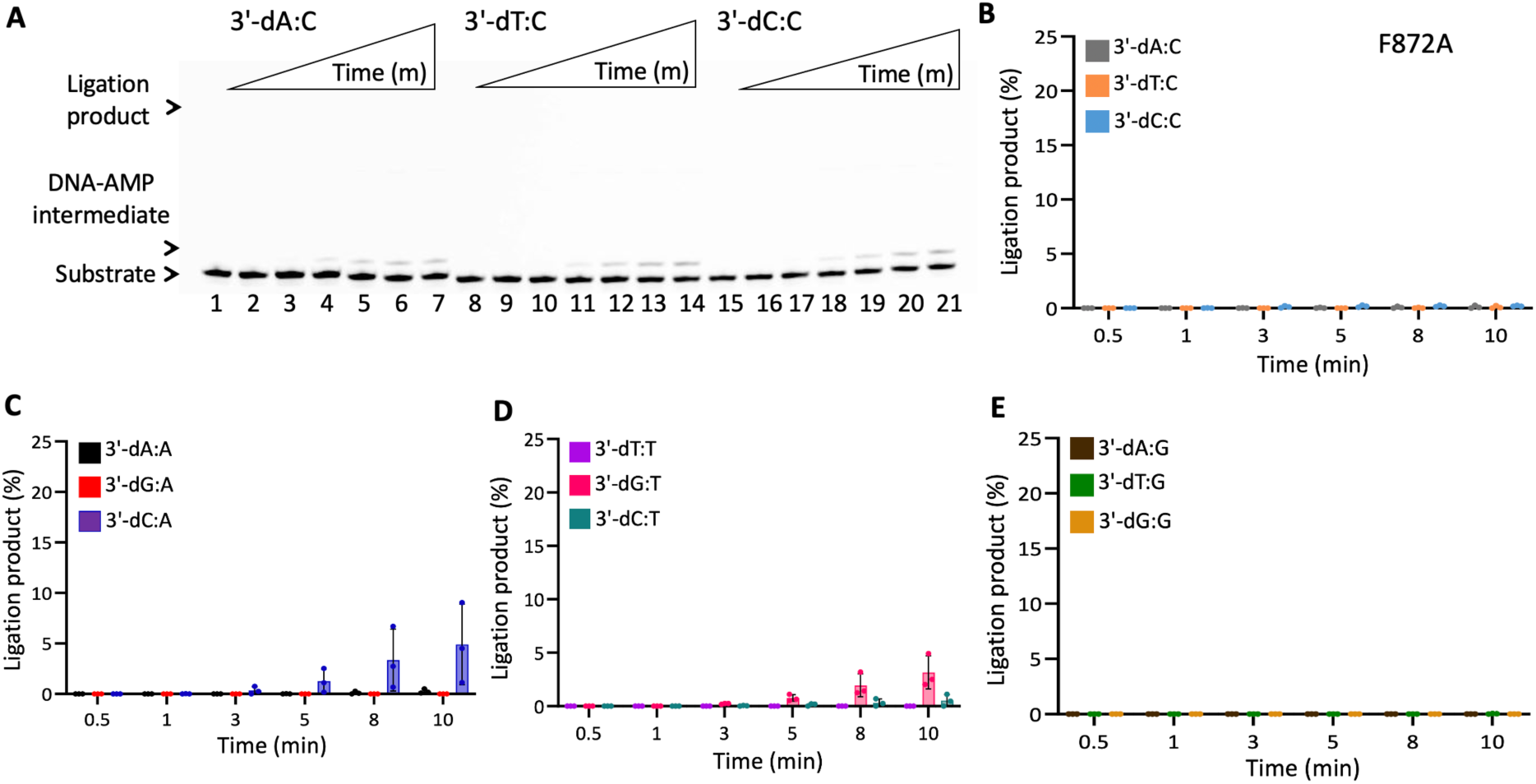
DNA substrate specificity of LIG1 F872A for all 12 mismatches. (**A**) Lanes 1, 8, and 15 are the negative enzyme controls for nick substrates with 3’-dA:C, 3’-dT:C, and 3’-dC:C, respectively. Lanes 2-7, 9-14, and 16-21 are the ligation reaction products and correspond to time points of 0.5, 1, 3, 5, 8, and 10 min. (**B-E**) Graphs show time-dependent changes in the amount of ligation products. The data represent the average from three independent experiments ± SD.

**Figure 5.**
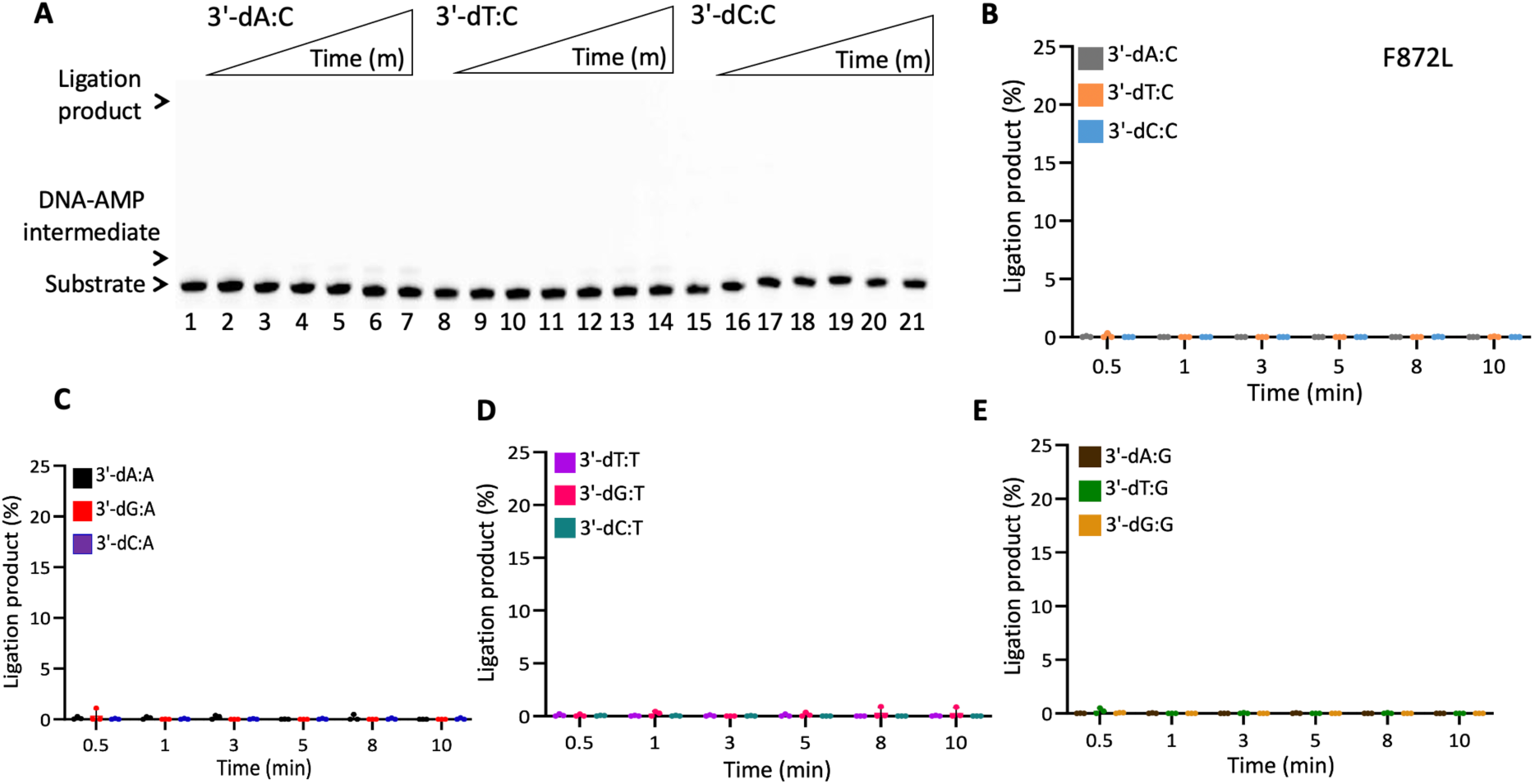
DNA substrate specificity of LIG1 F872L for all 12 mismatches. (**A**) Lanes 1, 8, and 15 are the negative enzyme controls for nick substrates with 3’-dA:C, 3’-dT:C, and 3’-dC:C, respectively. Lanes 2-7, 9-14, and 16-21 are the ligation reaction products and correspond to time points of 0.5, 1, 3, 5, 8, and 10 min. (**B-E**) Graphs show time-dependent changes in the amount of ligation products. The data represent the average from three independent experiments ± SD.

### Nick sealing efficiency of damaged ends by LIG1 active site mutants

In addition to all 12 mismatches, we characterized the ligation profile of all four LIG1 active site mutants for nick DNA substrates containing 3’-8oxodG opposite template base A, T, G, or C (Supplementary Scheme 2B). In line with our previous reports (27–30), for LIG1 wild-type, we observed the mutagenic ligation of 3’-8oxodG:A and the formation of DNA-AMP intermediate is accompanied by nick sealing of 3’-8oxodG:C (Figure 6A-B), while we obtained no ligation product in the presence of nick DNA substrates containing 3’-8oxodG:G and 3’-8oxodG:T with a time-dependent accumulation of DNA-AMP intermediates (Figure 6C-D). For F635A mutant, we observed reduced mutagenic ligation of 3’-8oxodG:A along with the accumulation of more DNA-AMP intermediates (Figure 7A, lanes 2-7) with a time-dependent increase in the amount of ligation products (Figure 7B). However, our results showed a diminished nick sealing for all other substrates containing 3’-8oxoG:C (Figure 7A, lanes 9-14), 3’-8oxodG:G and 3’-8oxodG:T (Figure 7C-D). For F635L, we obtained similar results with F635A for nick substrates containing 3’-8oxodG opposite A, T, G, or C (Figure 8). LIG1 F872A and F872L mutants were not able to seal any of nick substrates with damaged ends (Figures 9-10) and we obtained minor accumulation of DNA-AMP intermediates by F872A only for 3’-8oxodG:A and 3’-8oxodG:C (Figures 9A-B). Overall results demonstrated that F635 and F872 mutations decrease the mutagenic ligation of 3’-8oxodG:A when compared to wild-type enzyme, and we observed a complete ablation in nick sealing efficiency with F872A and F872L mutants.

**Figure 6.**
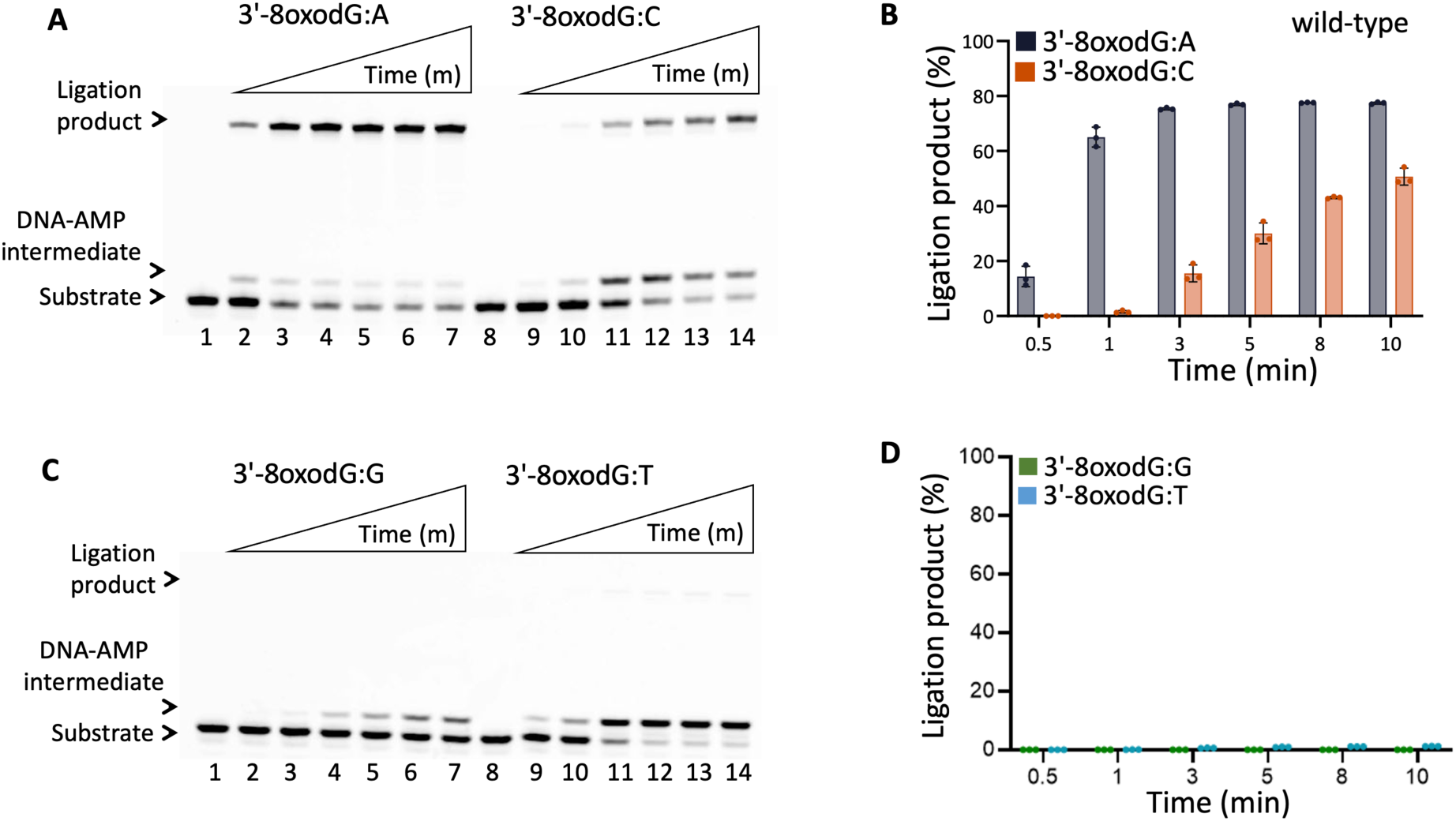
Ligation efficiency of nick DNA with damaged ends by LIG1 wild-type. (**A,C**) Lanes 1 and 8 are the negative enzyme controls for nick substrates with 3’-8oxodG:A and 3’-8oxodG:C (A) and 3’-8oxodG:G and 3’-8oxodG:T (C). Lanes 2-7 and 9-14 are the ligation reaction products and correspond to time points of 0.5, 1, 3, 5, 8, and 10 min. (**B,D**) Graphs show time-dependent changes in the amount of ligation products. The data represent the average from three independent experiments ± SD.

**Figure 7.**
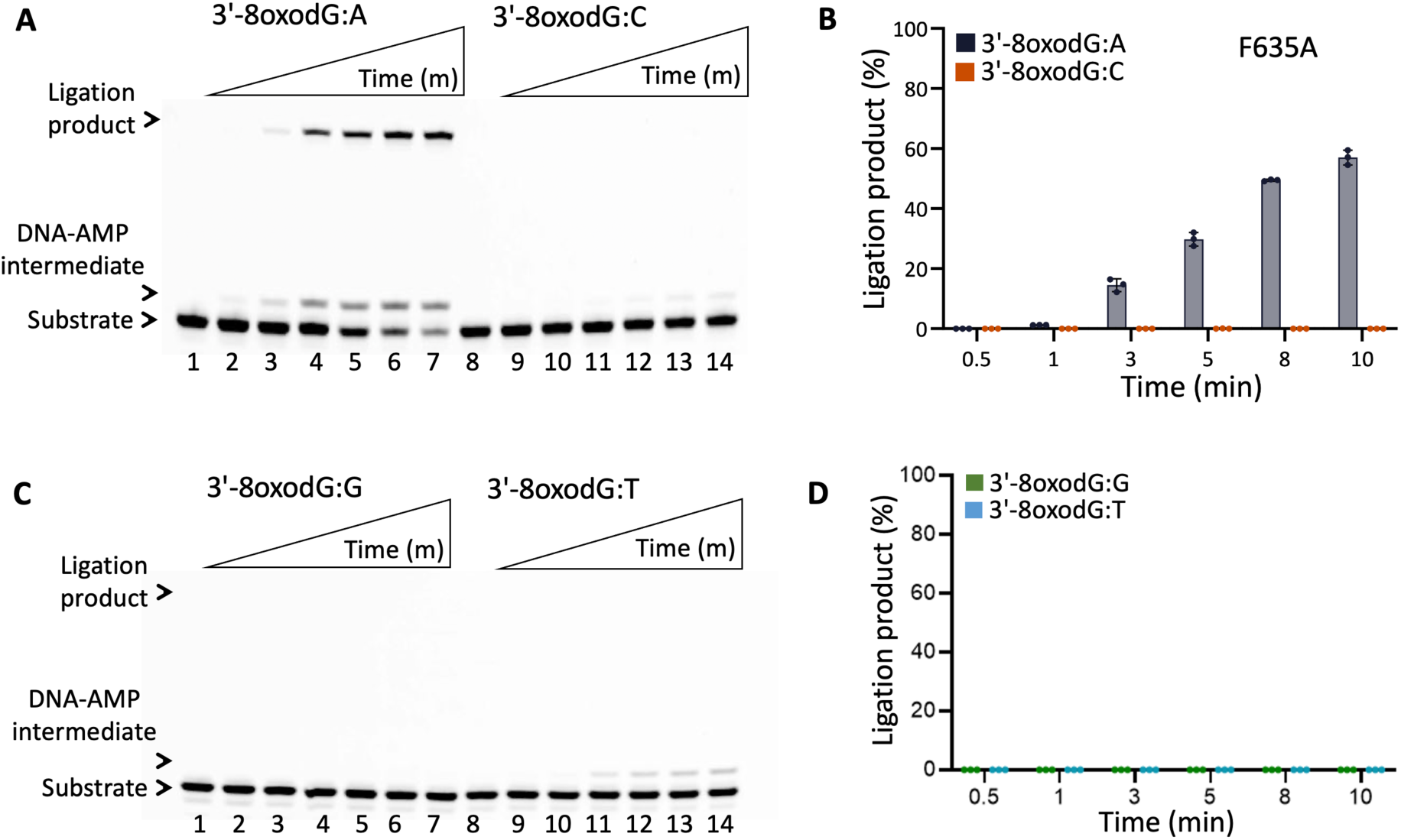
Ligation efficiency of nick DNA with damaged ends by LIG1 F635A. (**A,C**) Lanes 1 and 8 are the negative enzyme controls for nick substrates with 3’-8oxodG:A and 3’-8oxodG:C (A) and 3’-8oxodG:G and 3’-8oxodG:T (C). Lanes 2-7 and 9-14 are the ligation reaction products and correspond to time points of 0.5, 1, 3, 5, 8, and 10 min. (**B,D**) Graphs show time-dependent changes in the amount of ligation products. The data represent the average from three independent experiments ± SD.

**Figure 8.**
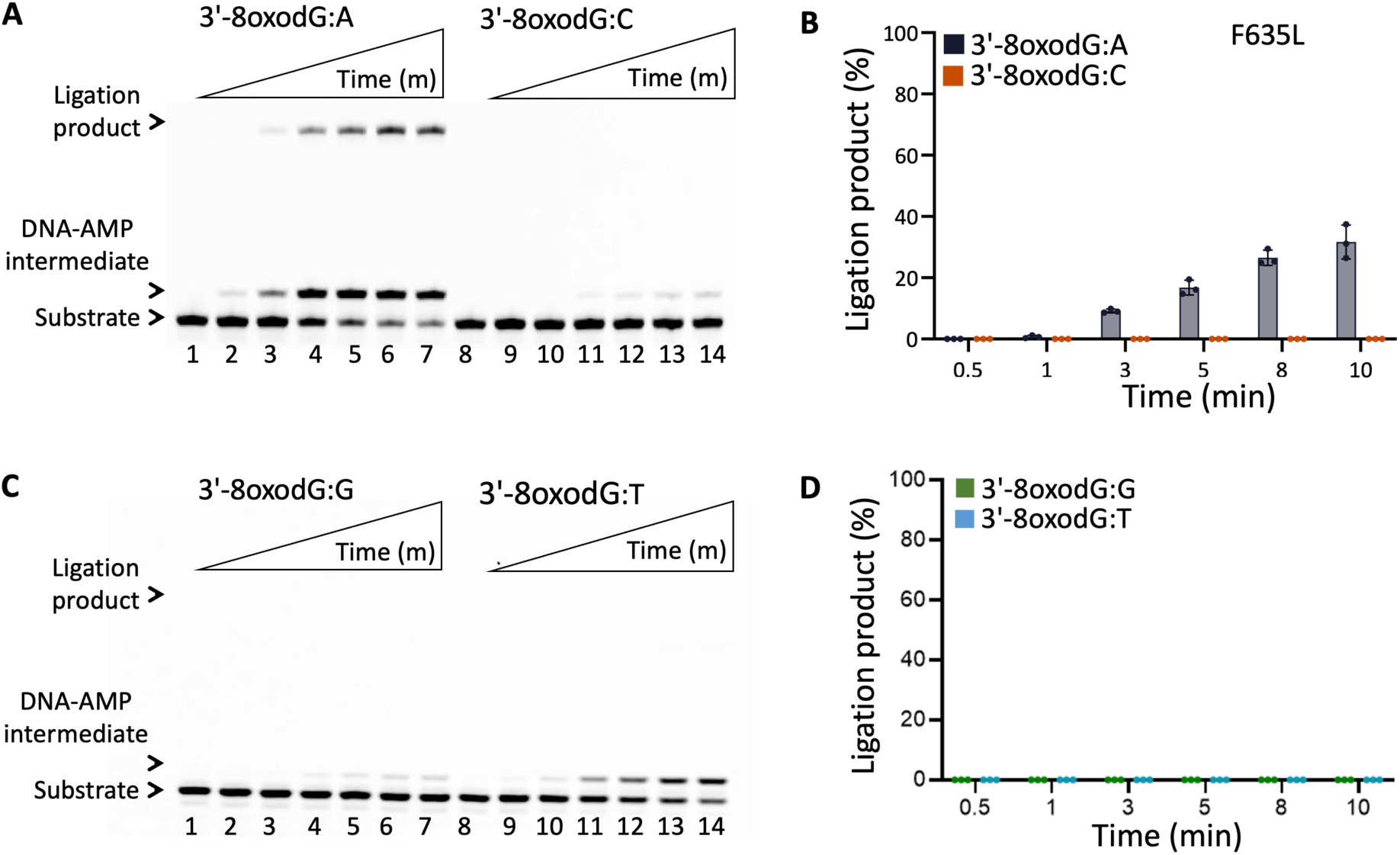
Ligation efficiency of nick DNA with damaged ends by LIG1 F635L. (**A,C**) Lanes 1 and 8 are the negative enzyme controls for nick substrates with 3’-8oxodG:A and 3’-8oxodG:C (A) and 3’-8oxodG:G and 3’-8oxodG:T (C). Lanes 2-7 and 9-14 are the ligation reaction products and correspond to time points of 0.5, 1, 3, 5, 8, and 10 min. (**B,D**) Graphs show time-dependent changes in the amount of ligation products. The data represent the average from three independent experiments ± SD.

**Figure 9.**
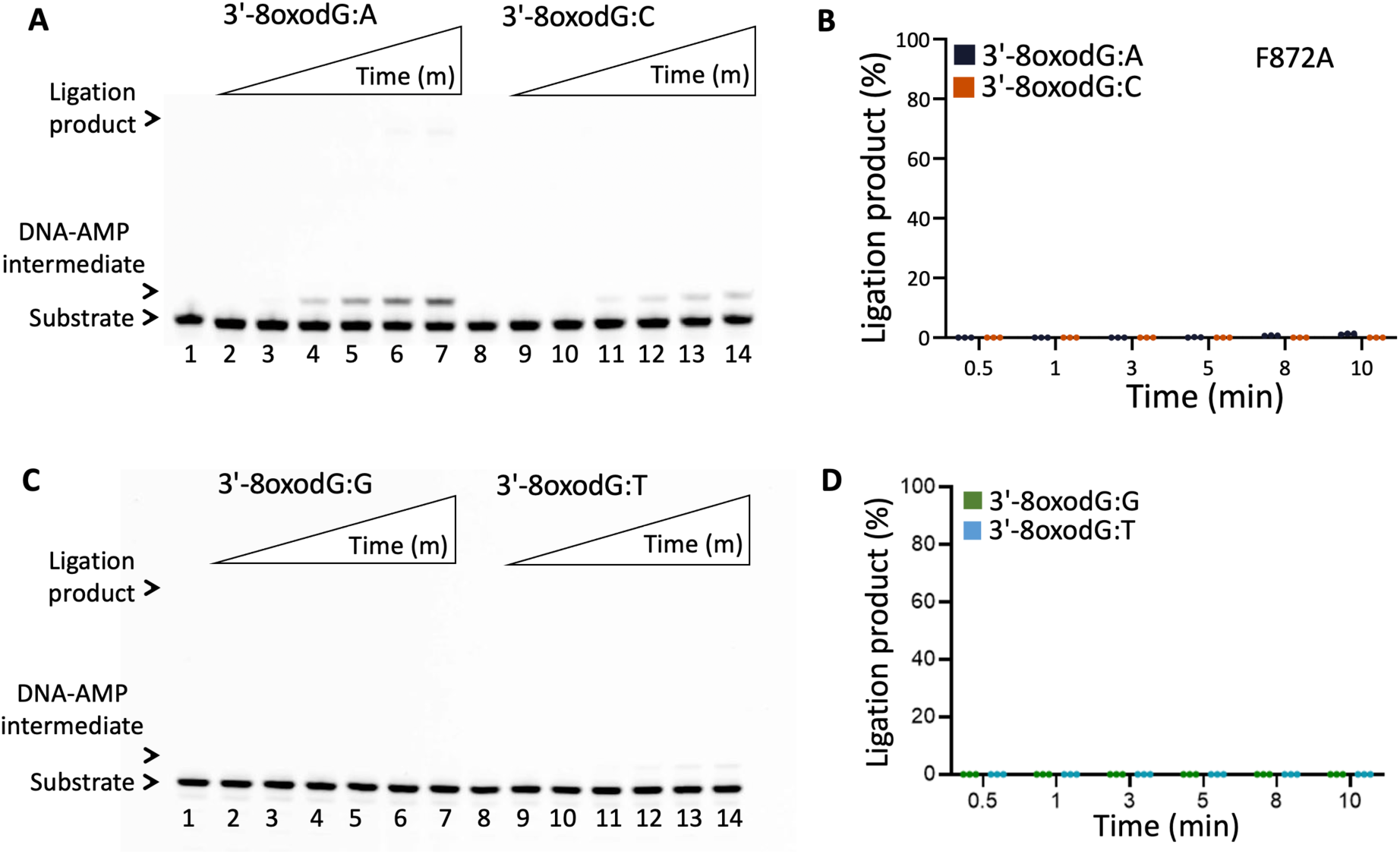
Ligation efficiency of nick DNA with damaged ends by LIG1 F872A. (**A,C**) Lanes 1 and 8 are the negative enzyme controls for nick substrates with 3’-8oxodG:A and 3’-8oxodG:C (A) and 3’-8oxodG:G and 3’-8oxodG:T (C). Lanes 2-7 and 9-14 are the ligation reaction products and correspond to time points of 0.5, 1, 3, 5, 8, and 10 min. (**B,D**) Graphs show time-dependent changes in the amount of ligation products. The data represent the average from three independent experiments ± SD.

### Role of LIG1 F635 and F872 for sugar discrimination against nick DNA containing a 3’-ribonucleotide

In addition to all 12 possible non-canonical mismatches and nick substrates with damaged ends, we comprehensively investigated the ligation efficiency of four LIG1 active site mutants in the presence of nick DNA substrates containing a single ribonucleotide at 3’-end, which mimic the products of ribonucleotide incorporation by repair or replication DNA polymerases. For this purpose, we used nick substrates carrying 3’-rA, 3’-rG, or 3’-rC opposite A, T, G, or C on a template position of DNA (Supplementary Scheme 2C). As we reported previously (34), our results with LIG1 wild-type showed very efficient ligation regardless of the base pairing architecture of the nick substrate, demonstrating a lack of sugar discrimination against 3’-ribonucleotide (Figure 11 and Supplementary Figure 12). We only showed relatively less efficient ligation in the presence of 3’-rA:G, 3’-rG:C, and 3’-rC:C compared to the analogous 3’-deoxyribonucleotide nick DNA substrates. For LIG1 variants carrying mutations at F635, our results demonstrated different nick sealing efficiency than that of LIG1 wild-type except canonical nicks containing 3’-rA:T, 3’-rG:C, and 3’-rC:G (Figures 12-13 and Supplementary Figures 13-14). For example, in the presence of nicks containing 3’-rA:G, 3’-rG:A, and 3’-rC:C, we obtained diminished ligation especially by LIG1 F635A (Figure 12 and Supplementary Figure 13). We also obtained less efficient nick sealing for 3’-rA:G, 3’-rG:A, and 3’-rC:C by F635L, and the end joining of all other 3’-ribonucleotides was relatively higher than F635A and similar to the wild-type enzyme (Figure 13 and Supplementary Figure 14). For LIG1 F872A and F872L mutants, our results showed a decrease in overall ligation efficiency when compared to LIG1 wild-type being able to only ligate nicks containing canonical 3’-ribonucleotides (Figures 14-15 and Supplementary Figures 15-16). Notably, F872L showed the most drastic decrease in ligation efficiency. In addition to the nick substrates (3’-rA:G, 3’-rG:A, and 3’-rC:C) that cannot be ligated efficiently by F635A and F635L mutants, we obtained less than 10 % of ligation products in the presence of 3’-rG:G, 3’-rA:A, 3’-rC:A, and 3’-rC:T by LIG1 F872L mutant (Figure 15 and Supplementary Figure 16). Overall, these mutants showed greater reliance on the architecture of proper 3’-ribonuceotide:template base pairing to seal nicks containing a “wrong sugar” at 3’-end of nick DNA.

**Figure 10.**
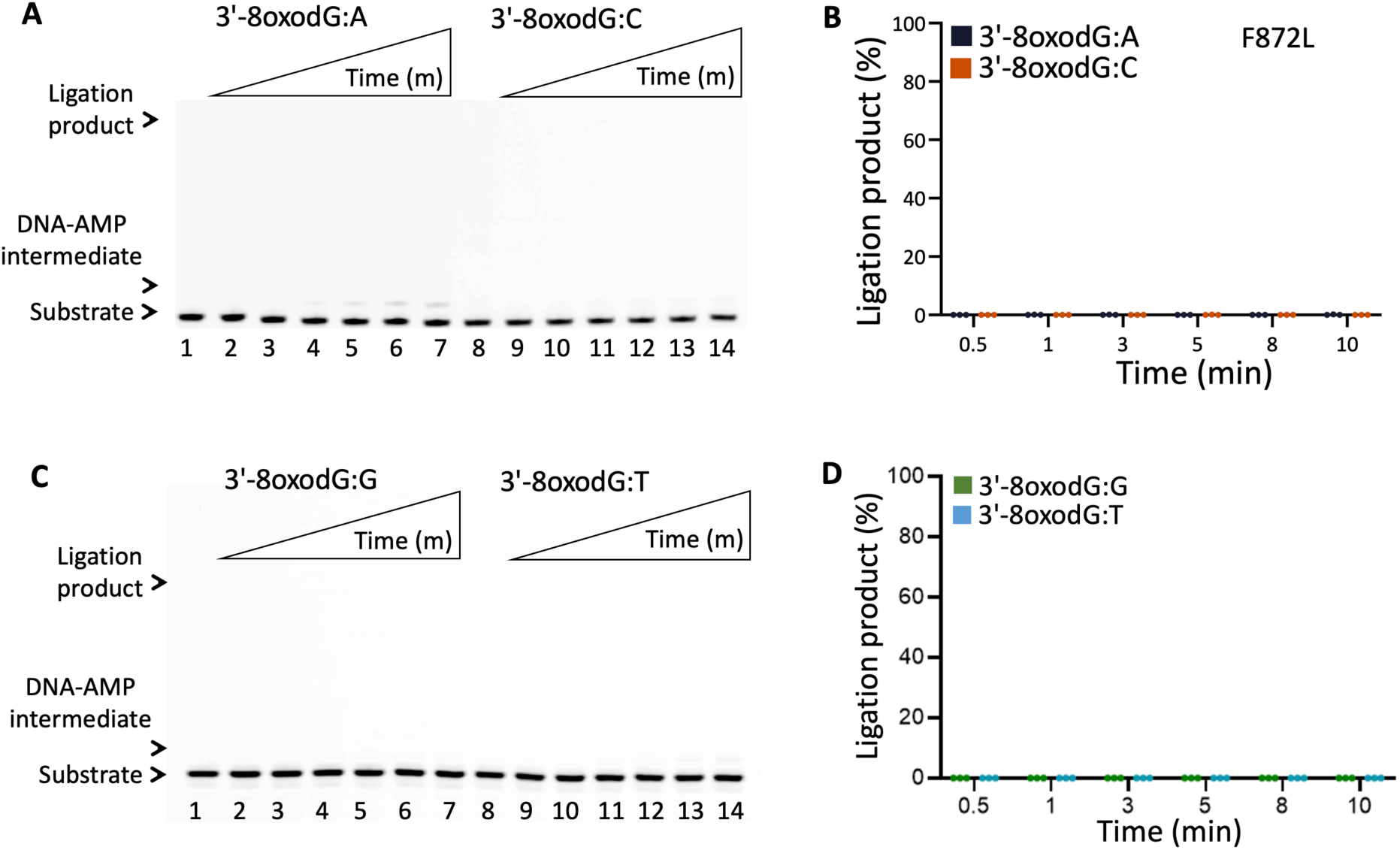
Ligation efficiency of nick DNA with damaged ends by LIG1 F872L. (**A,C**) Lanes 1 and 8 are the negative enzyme controls for nick substrates with 3’-8oxodG:A and 3’-8oxodG:C (A) and 3’-8oxodG:G and 3’-8oxodG:T (C). Lanes 2-7 and 9-14 are the ligation reaction products and correspond to time points of 0.5, 1, 3, 5, 8, and 10 min. (**B,D**) Graphs show time-dependent changes in the amount of ligation products. The data represent the average from three independent experiments ± SD.

**Figure 11.**
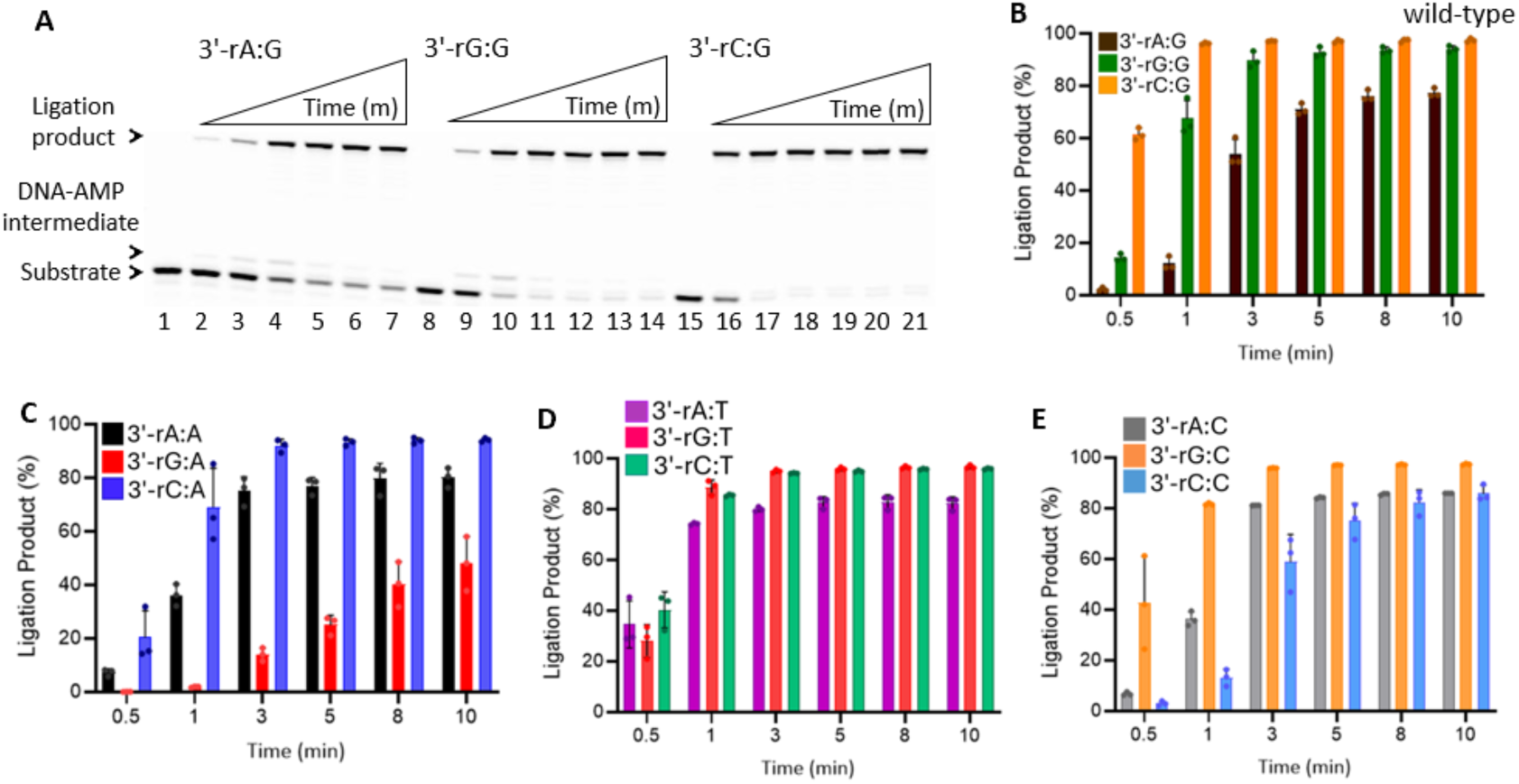
Sugar discrimination of LIG1 wild-type against nick DNA containing 3’-ribonucleotide. (**A**) Lanes 1, 8, and 15 are the negative enzyme controls for nick substrates with 3’-rA:G, 3’-rG:G, and 3’-rC:G, respectively. Lanes 2-7, 9-14, and 16-21 are the ligation reaction products for 3’-rA:G, 3’-rG:G, and 3’-rC:G, respectively, and correspond to time points of 0.5, 1, 3, 5, 8, and 10 min. (**B-E**) Graphs show time-dependent changes in the amount of ligation products. The data represent the average from three independent experiments ± SD.

**Figure 12.**
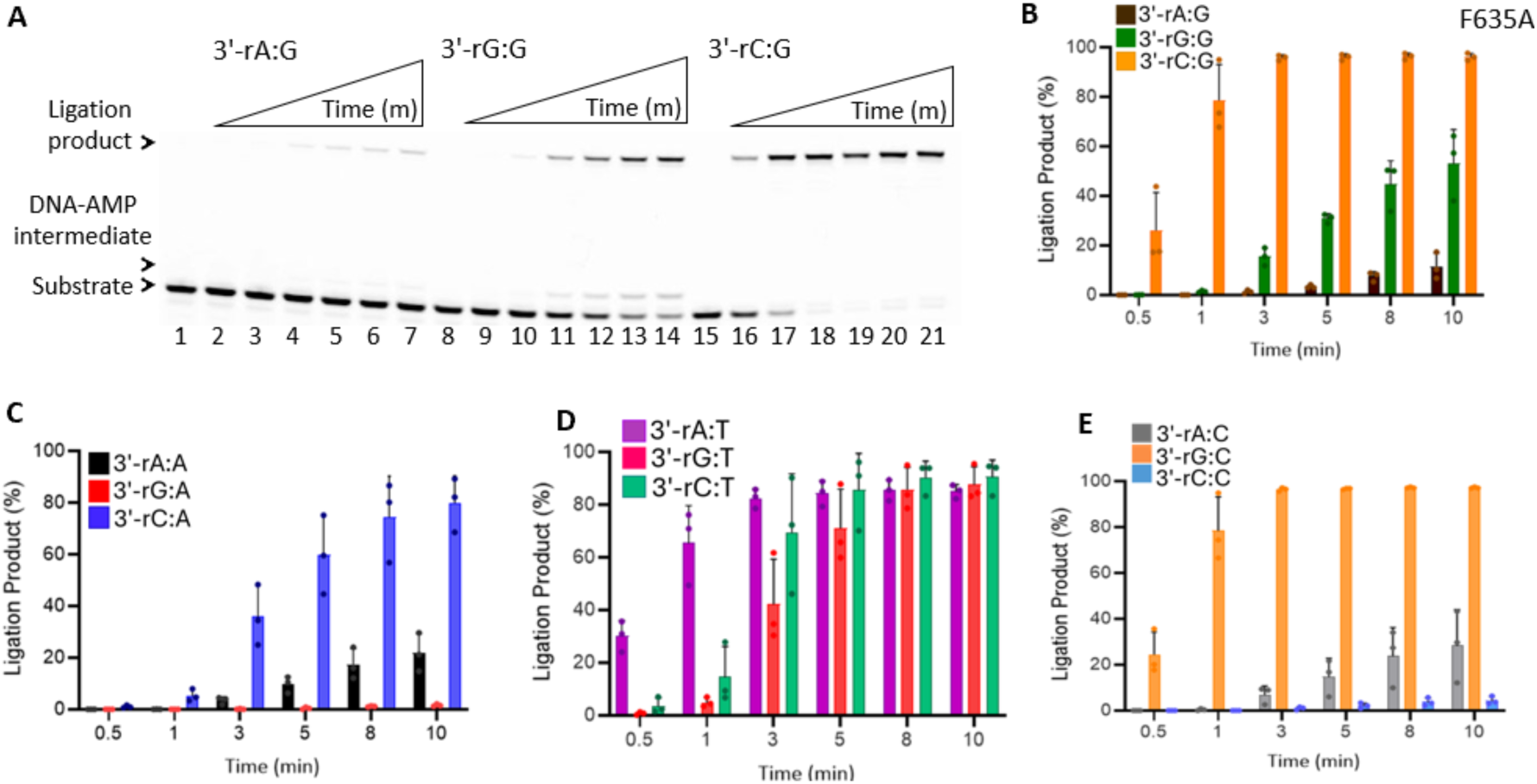
Sugar discrimination of LIG1 F635A against nick DNA containing 3’-ribonucleotide. (**A**) Lanes 1, 8, and 15 are the negative enzyme controls for nick substrates with 3’-rA:G, 3’-rG:G, and 3’-rC:G, respectively. Lanes 2-7, 9-14, and 16-21 are the ligation reaction products for 3’-rA:G, 3’-rG:G, and 3’-rC:G, respectively, and correspond to time points of 0.5, 1, 3, 5, 8, and 10 min. (**B-E**) Graphs show time-dependent changes in the amount of ligation products. The data represent the average from three independent experiments ± SD.

**Figure 13.**
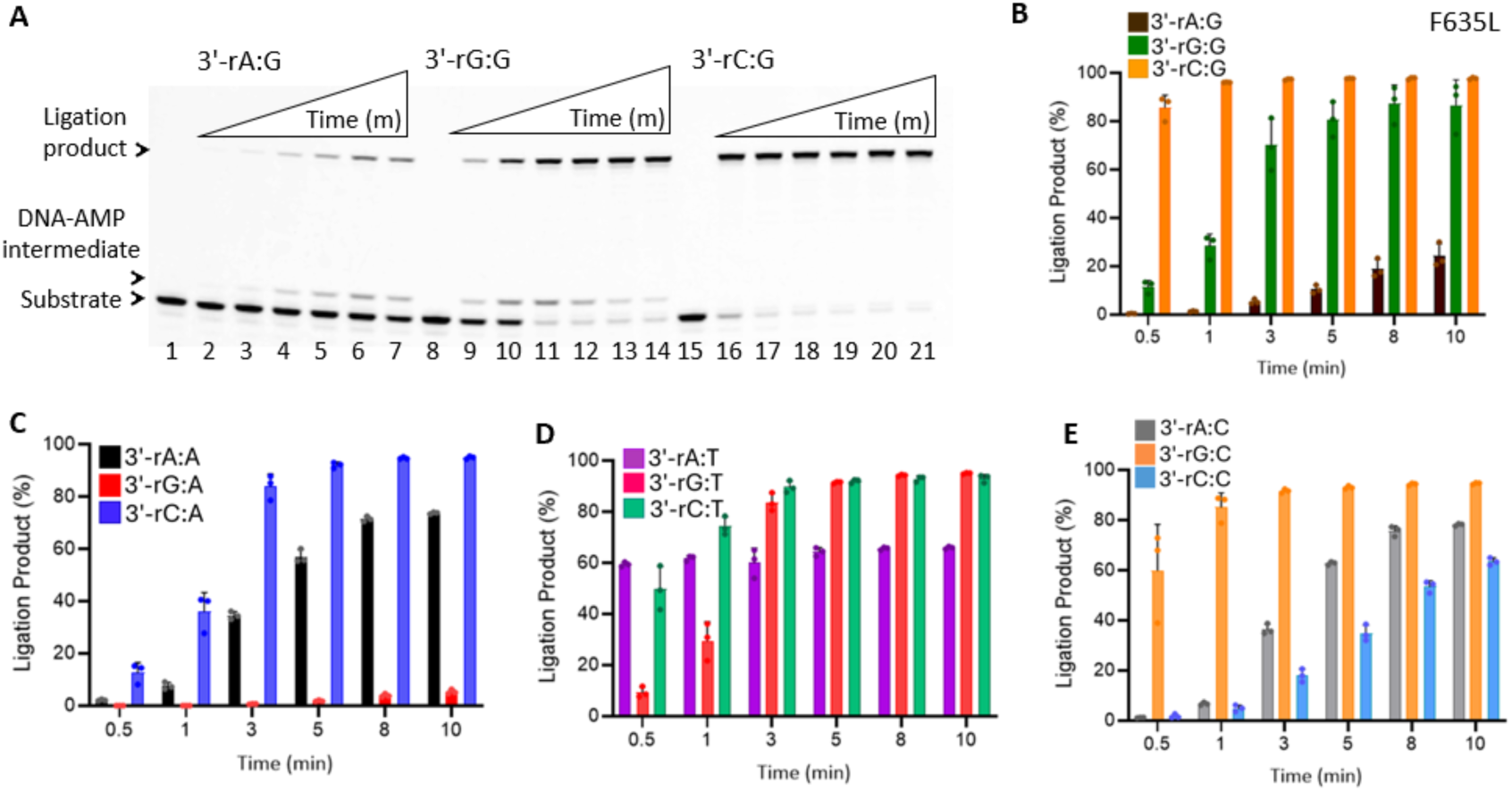
Sugar discrimination of LIG1 F635L against nick DNA containing 3’-ribonucleotide. (**A**) Lanes 1, 8, and 15 are the negative enzyme controls for nick substrates with 3’-rA:G, 3’-rG:G, and 3’-rC:G, respectively. Lanes 2-7, 9-14, and 16-21 are the ligation reaction products for 3’-rA:G, 3’-rG:G, and 3’-rC:G, respectively, and correspond to time points of 0.5, 1, 3, 5, 8, and 10 min. (**B-E**) Graphs show time-dependent changes in the amount of ligation products. The data represent the average from three independent experiments ± SD.

**Figure 14.**
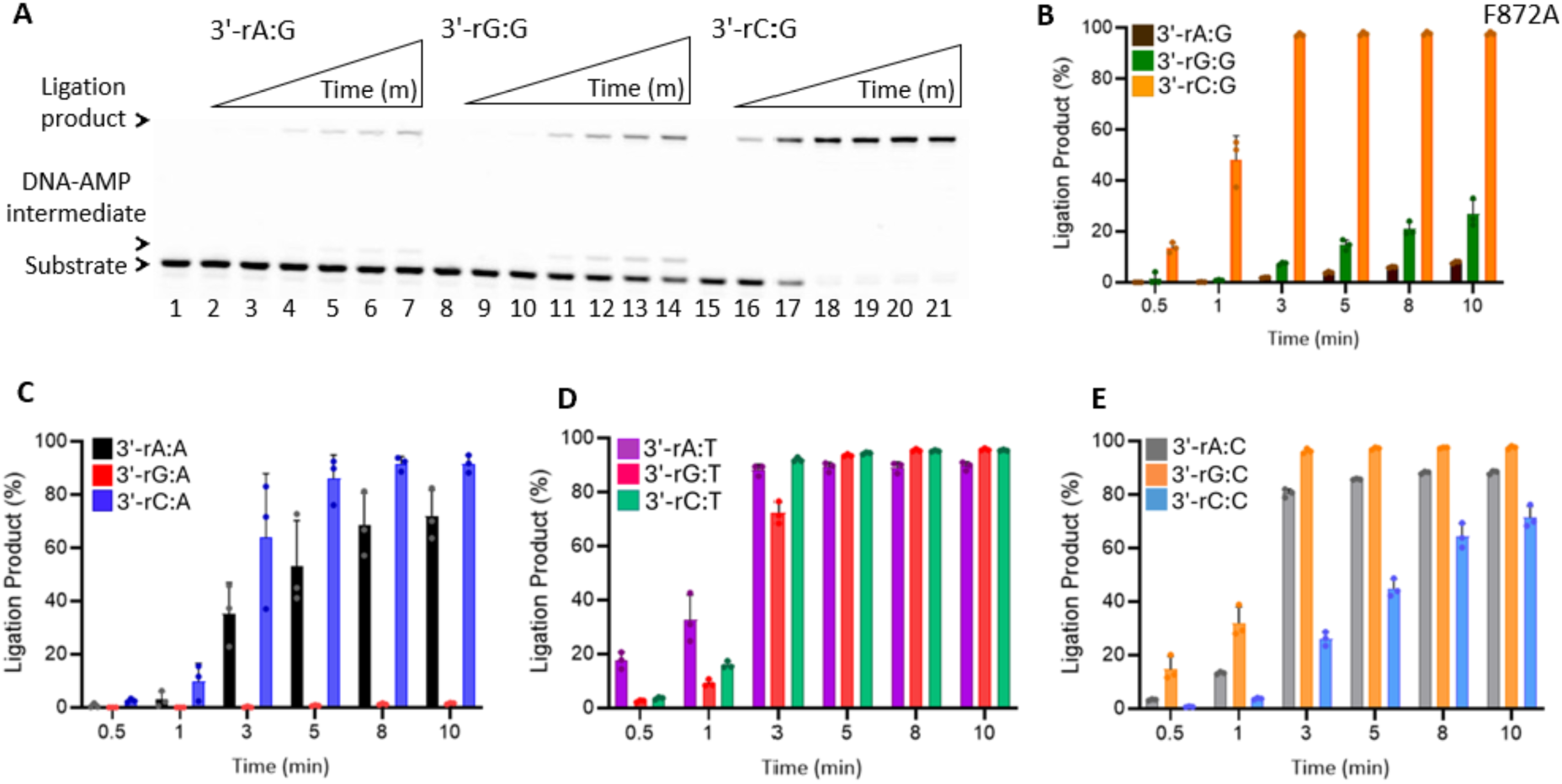
Sugar discrimination of LIG1 F872A against nick DNA containing 3’-ribonucleotide. (**A**) Lanes 1, 8, and 15 are the negative enzyme controls for nick substrates with 3’-rA:G, 3’-rG:G, and 3’-rC:G, respectively. Lanes 2-7, 9-14, and 16-21 are the ligation reaction products for 3’-rA:G, 3’-rG:G, and 3’-rC:G, respectively, and correspond to time points of 0.5, 1, 3, 5, 8, and 10 min. (**B-E**) Graphs show time-dependent changes in the amount of ligation products. The data represent the average from three independent experiments ± SD.

**Figure 15.**
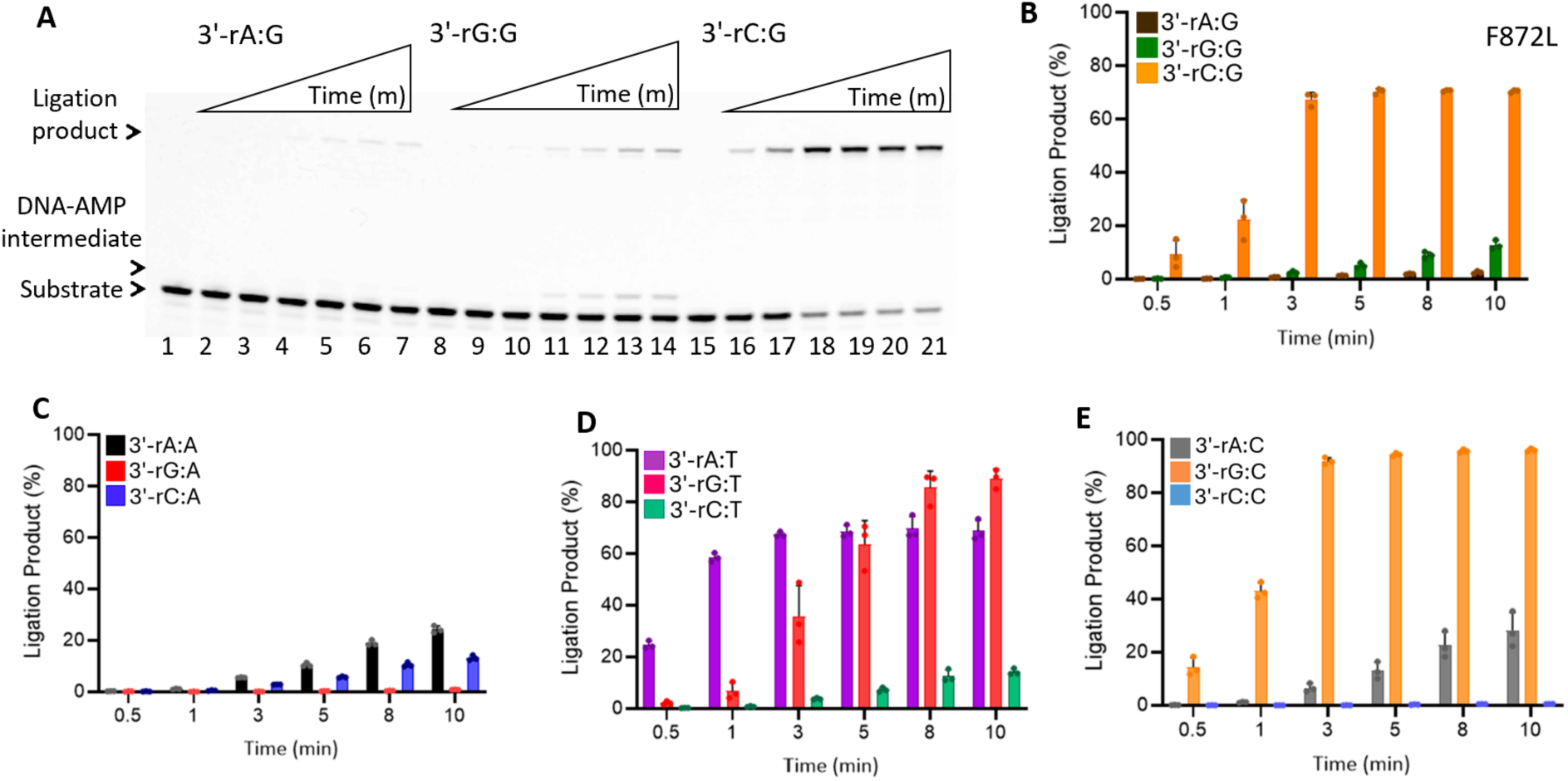
Sugar discrimination of LIG1 F872L against nick DNA containing 3’-ribonucleotide. (**A**) Lanes 1, 8, and 15 are the negative enzyme controls for nick substrates with 3’-rA:G, 3’-rG:G, and 3’-rC:G, respectively. Lanes 2-7, 9-14, and 16-21 are the ligation reaction products for 3’-rA:G, 3’-rG:G, and 3’-rC:G, respectively, and correspond to time points of 0.5, 1, 3, 5, 8, and 10 min. (**B-E**) Graphs show time-dependent changes in the amount of ligation products. The data represent the average from three independent experiments ± SD.

### Role of LIG1 F635 and F872 for sugar discrimination against nick DNA containing a 5’-ribonucleotide

To elucidate the roles of F635 and F872 residues for sugar discrimination at both ends of nick DNA, we also investigated the ligation efficiency of four LIG1 active site mutants in the presence of nick DNA substrates containing a single ribonucleotide at 5’-end which mimic the intermediates of RER that is initiated by RNase H2 and results in error-free removal of mis-incorporated ribonucleotides. To test this, we used nick substrates carrying 5’-rA, 5’-rG, or 5’-rC opposite A, T, G, or C (Supplementary Scheme 2D). First, it’s important to note that there was almost no ligation product by either LIG1 wild-type or active site mutants in the presence nick substrates containing 5’-ribonucleotide at the time points (0.5-10 min) of ligation reaction where we tested the nick sealing of 3’-ribonucleotides (Figures 11-15). Therefore, the ligation profile for nicks containing 5’-ribonucleotides was evaluated for longer time points (0.5-60 min). Our results for LIG1 wild-type indicates an ability to discriminate sugar at the 5’-end of the nick except canonical substrates 5’-rA:T, 5’-rG:C, and 5’-rC:G (Figure 16 and Supplementary Figure 17). For LIG1 variants carrying mutation at F635, there was complete ablation of nick sealing for all substrates, including the ones with canonical base pairing (Figures 17-18 and Supplementary Figures 18-19). However, LIG1 F872A mutant showed an increased ligation efficiency for the nick substrates containing 5’-ribonucleotide when compared to wild-type enzyme (Figure 19 and Supplementary Figure 20). Specifically, nick DNA with 5’-rA:T, 5’-rG:T, 5’-rG:C, 5’-rC:G, and 5’-rC:A all showed an increased amount of ligation product. In the presence of F872L, there was a decrease in ligation efficiency for nicks with 5’-ribonucleotides when compared to wild-type enzyme (Figure 20 and Supplementary Figure 21). Overall, these results show that LIG1 wild-type and mutants cannot efficiently seal nicks containing a “wrong” sugar at the 5’-end of the nick.

**Figure 16.**
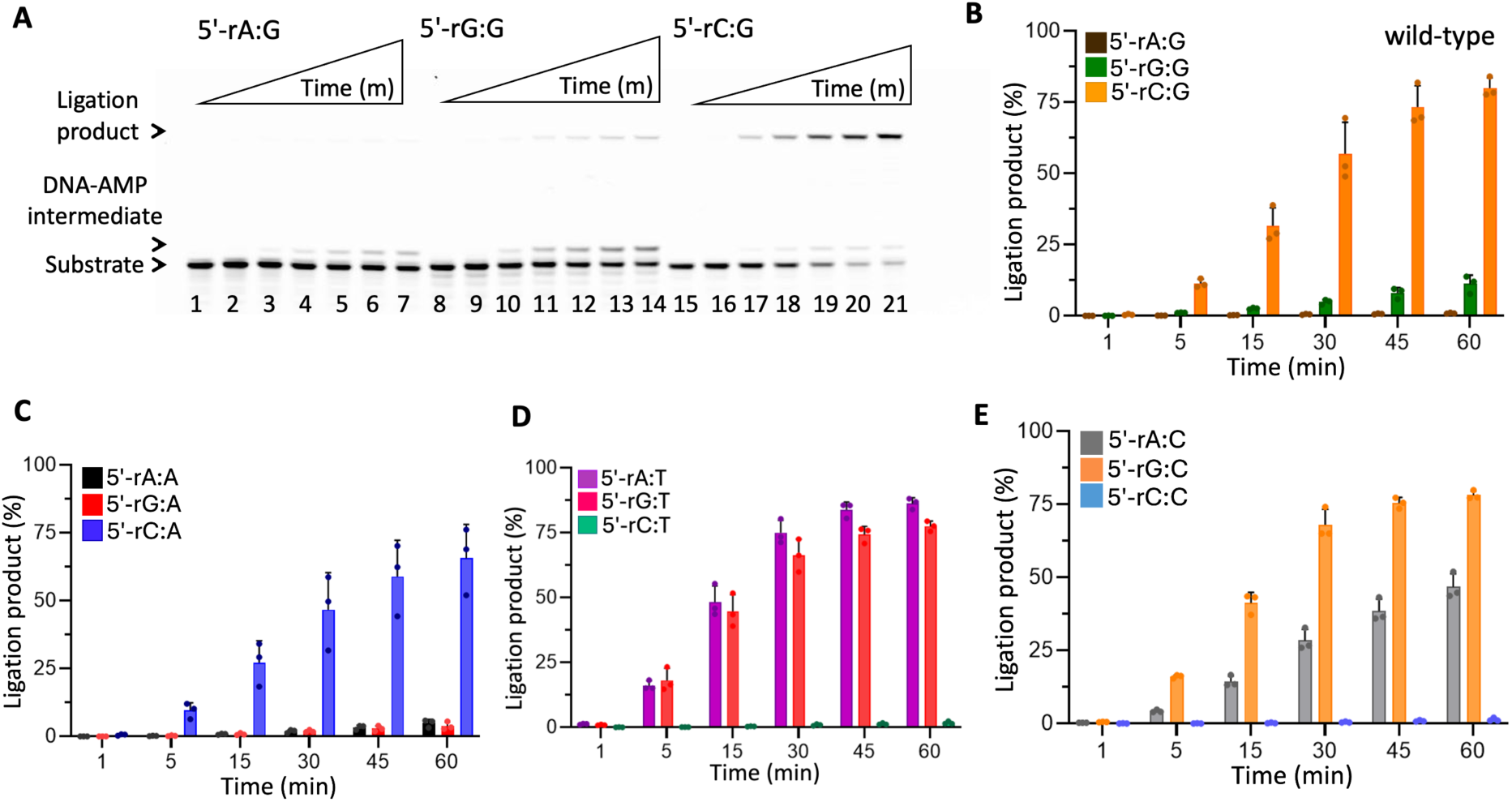
Sugar discrimination of LIG1 wild-type against nick DNA containing 5’-ribonucleotide. (**A**) Lanes 1, 8, and 15 are the negative enzyme controls for nick substrates with 5’-rA:G, 5’-rG:G, and 5’-rC:G, respectively. Lanes 2-7, 9-14, and 16-21 are the ligation reaction products for 5’-rA:G, 5’-rG:G, and 5’-rC:G, respectively, and correspond to time points of 1, 5, 15, 30, 45, and 60 min. (**B-E**) Graphs show time-dependent changes in the amount of ligation products. The data represent the average from three independent experiments ± SD.

**Figure 17.**
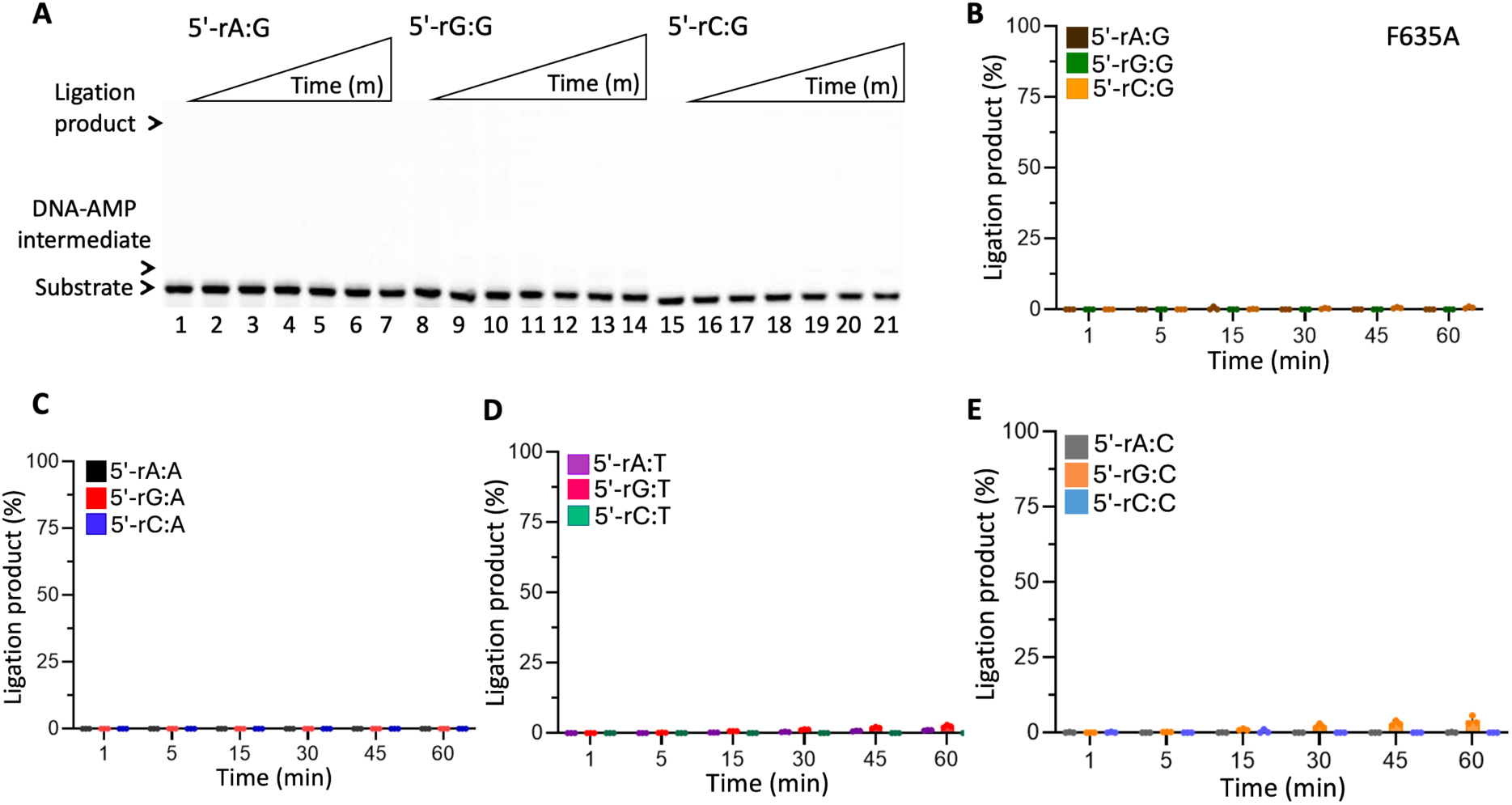
Sugar discrimination of LIG1 F635A against nick DNA containing 5’-ribonucleotide. (**A**) Lanes 1, 8, and 15 are the negative enzyme controls for nick substrates with 5’-rA:G, 5’-rG:G, and 5’-rC:G, respectively. Lanes 2-7, 9-14, and 16-21 are the ligation reaction products for 5’-rA:G, 5’-rG:G, and 5’-rC:G, respectively, and correspond to time points of 1, 5, 15, 30, 45, and 60 min. (**B-E**) Graphs show time-dependent changes in the amount of ligation products. The data represent the average from three independent experiments ± SD.

**Figure 18.**
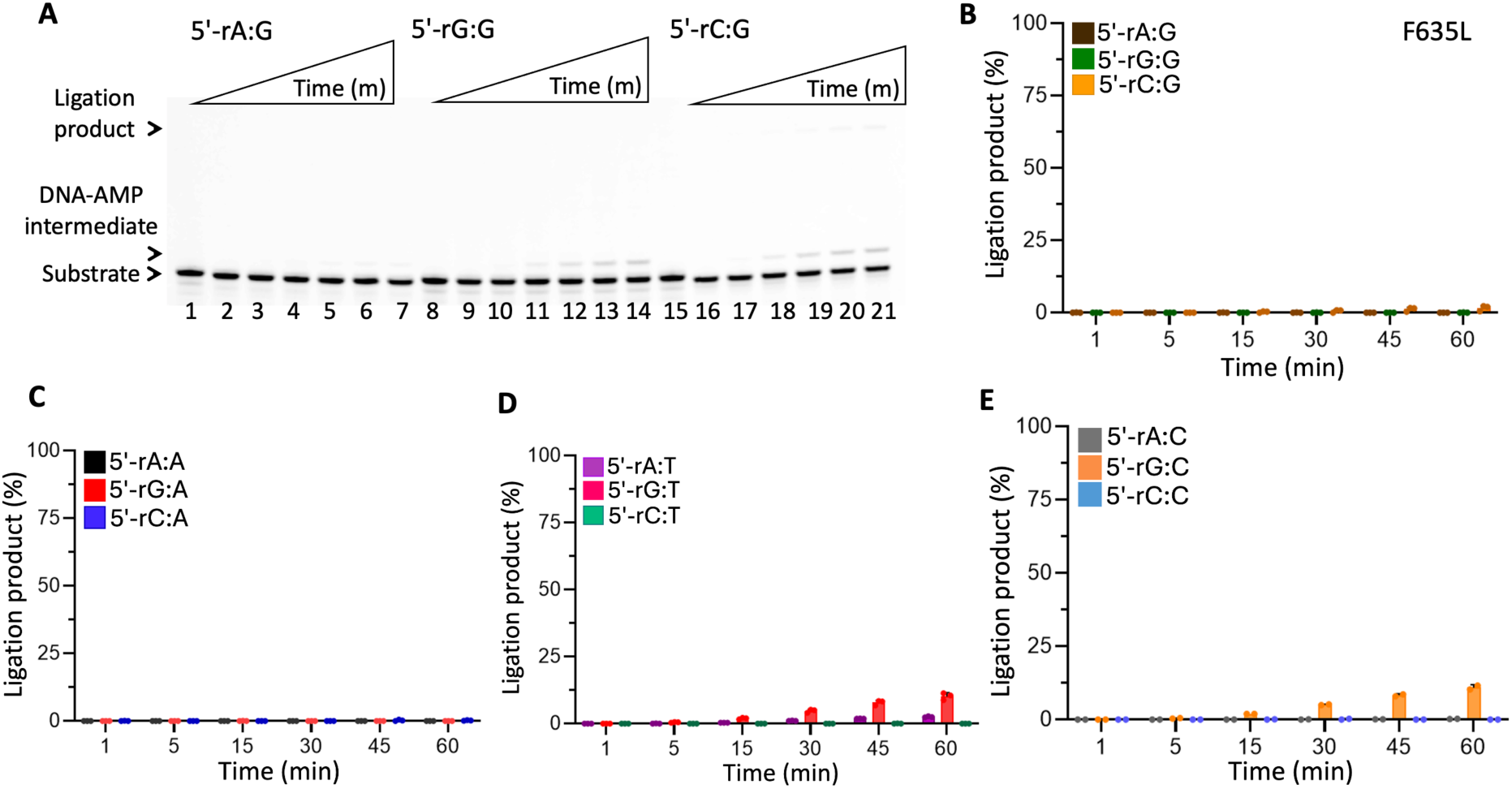
Sugar discrimination of LIG1 F635L against nick DNA containing 5’-ribonucleotide. (**A**) Lanes 1, 8, and 15 are the negative enzyme controls for nick substrates with 5’-rA:G, 5’-rG:G, and 5’-rC:G, respectively. Lanes 2-7, 9-14, and 16-21 are the ligation reaction products for 5’-rA:G, 5’-rG:G, and 5’-rC:G, respectively, and correspond to time points of 1, 5, 15, 30, 45, and 60 min. (**B-E**) Graphs show time-dependent changes in the amount of ligation products. The data represent the average from three independent experiments ± SD.

**Figure 19.**
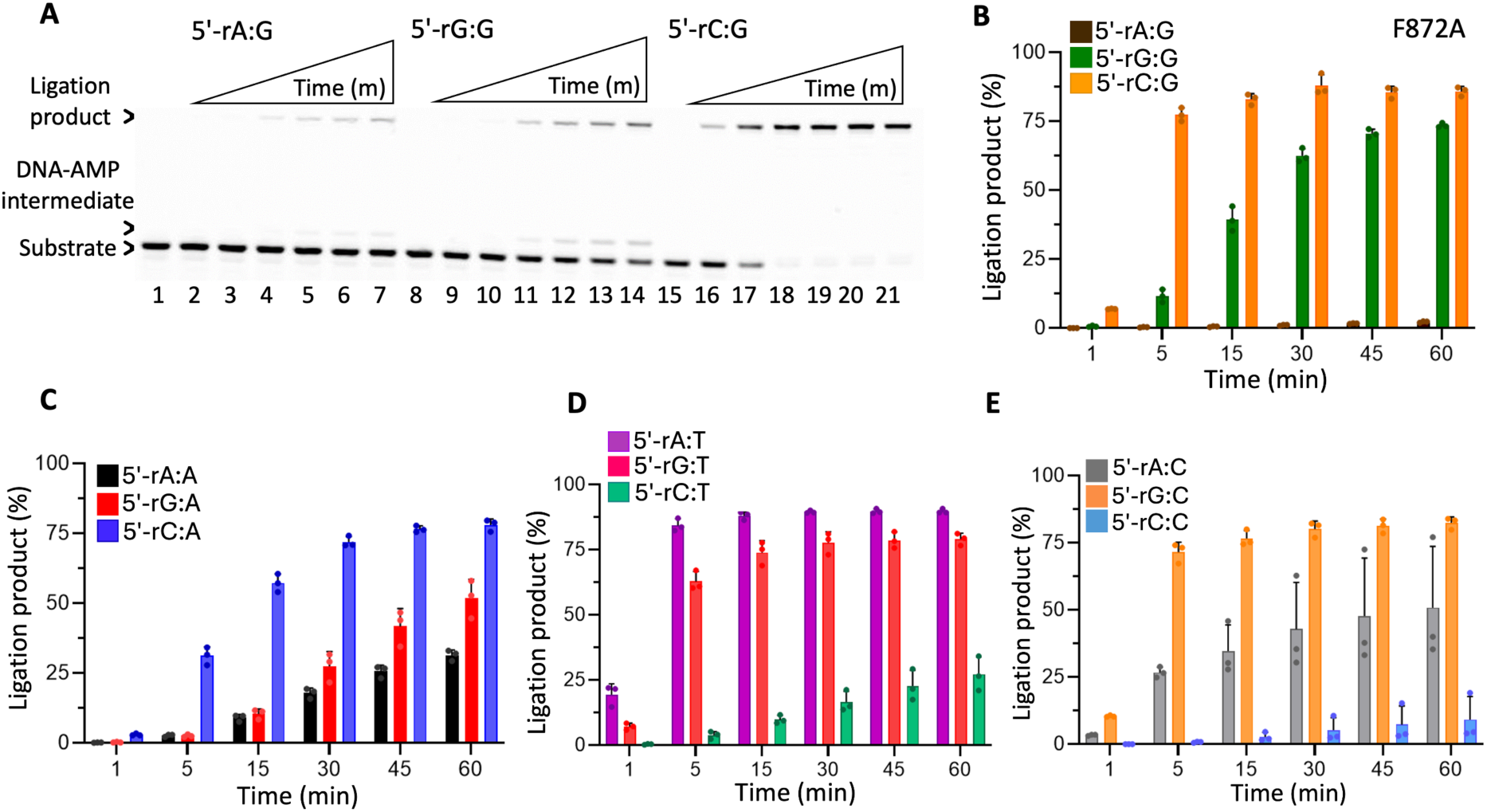
Sugar discrimination of LIG1 F872A against nick DNA containing 5’-ribonucleotide. (**A**) Lanes 1, 8, and 15 are the negative enzyme controls for nick substrates with 5’-rA:G, 5’-rG:G, and 5’-rC:G, respectively. Lanes 2-7, 9-14, and 16-21 are the ligation reaction products for 5’-rA:G, 5’-rG:G, and 5’-rC:G, respectively, and correspond to time points of 1, 5, 15, 30, 45, and 60 min. (**B-E**) Graphs show time-dependent changes in the amount of ligation products. The data represent the average from three independent experiments ± SD.

**Figure 20.**
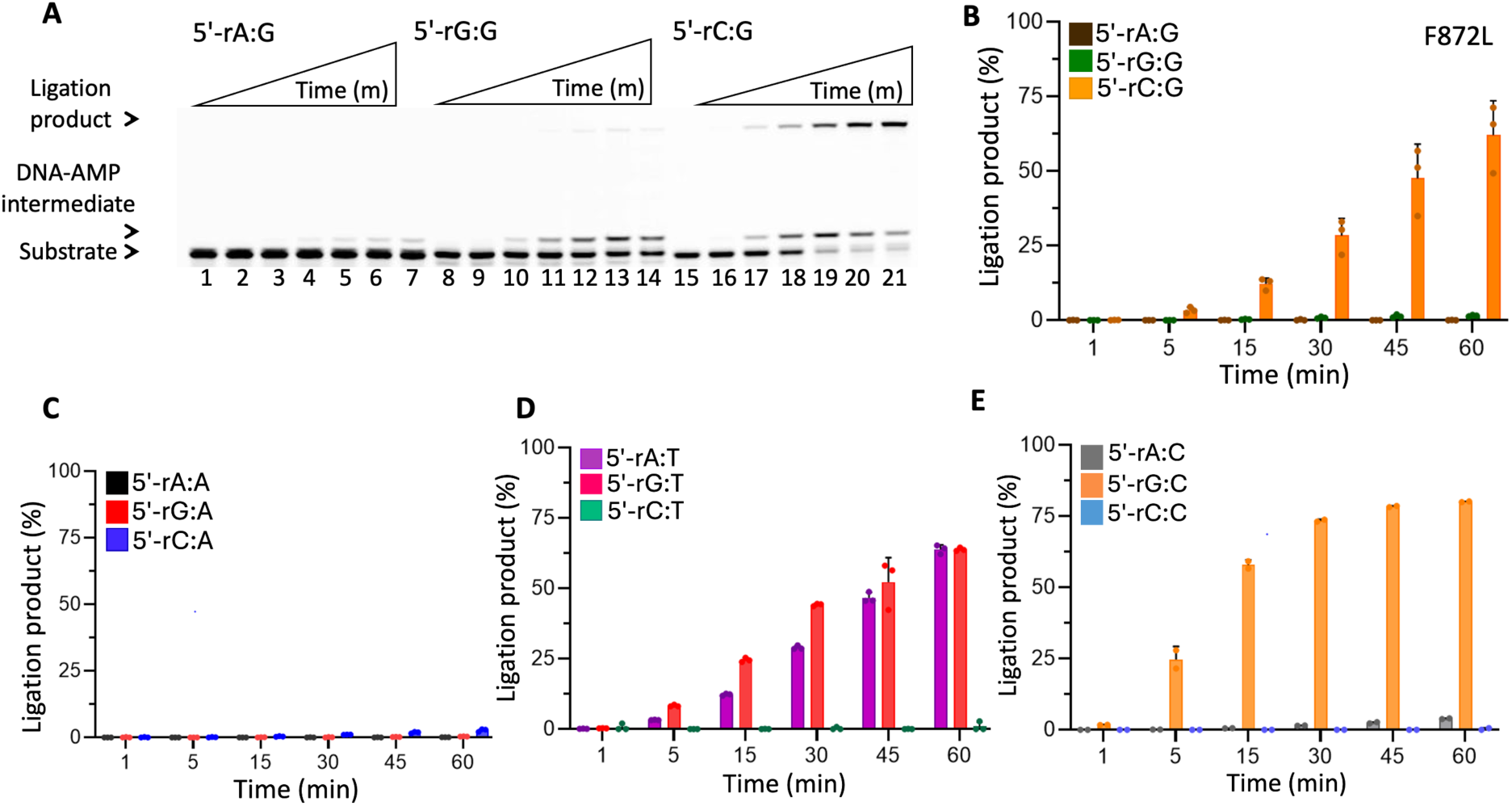
Sugar discrimination of LIG1 F872L against nick DNA containing 5’-ribonucleotide. (**A**) Lanes 1, 8, and 15 are the negative enzyme controls for nick substrates with 5’-rA:G, 5’-rG:G, and 5’-rC:G, respectively. Lanes 2-7, 9-14, and 16-21 are the ligation reaction products for 5’-rA:G, 5’-rG:G, and 5’-rC:G, respectively, and correspond to time points of 1, 5, 15, 30, 45, and 60 min. (**B-E**) Graphs show time-dependent changes in the amount of ligation products. The data represent the average from three independent experiments ± SD.

### Overall comparisons of ligation efficiency between LIG1 F635 and F872 mutants

To easily compare the impact of the active site mutations on ligation efficiency of 70 nick DNA substrates tested in this study, we generated dot plots using a single time point (3 or 15 min) from the bar graphs showing the time-dependent changes in the amount of ligation products (Figures 21-24). From these comparisons, we conclude that substituting the phenylalanine with either alanine or leucine at LIG1 residues F635 and F872 drastically impacts ligation of all 12 possible mismatches while not dramatically impacting the ligation of canonical substrates (Figure 21). For nicks containing 3’-8oxodG, mutagenic ligation is reduced and completely diminished by F635 and F872 mutations, respectively (Figure 22). We observed clear differences in the ligation efficiencies between 3’-deoxyribonucleotide and 3’-ribonucleotide containing substrates. LIG1 F635A, F635L, and F872A mutants all show decreased ligation profile, while F872L demonstrated a drastic surveillance for base pairing architecture at 3’-end (Figure 23). Comparing these dot plots, we suggest that LIG1 has evolved to not discriminate against 3’-ribonucleotides and that F635 and F872 do not participate in sugar discrimination. The dot plots averaging ligation product at 15 min time for nicks with 5’-ribonucleotides show that all four LIG1 mutants have different nick sealing efficiency than that of wild-type LIG1 (Figure 24). F635A/L mutants show complete inability to seal any nicks, F872A exhibits an increased ligation efficiency, while reduced ligation was observed by F872L mutant. These results demonstrate that LIG1 can discriminate sugars at 5’-end of the nick, and active site residues F635 and F872 do not participate in the discrimination mechanism.

**Figure 21.**
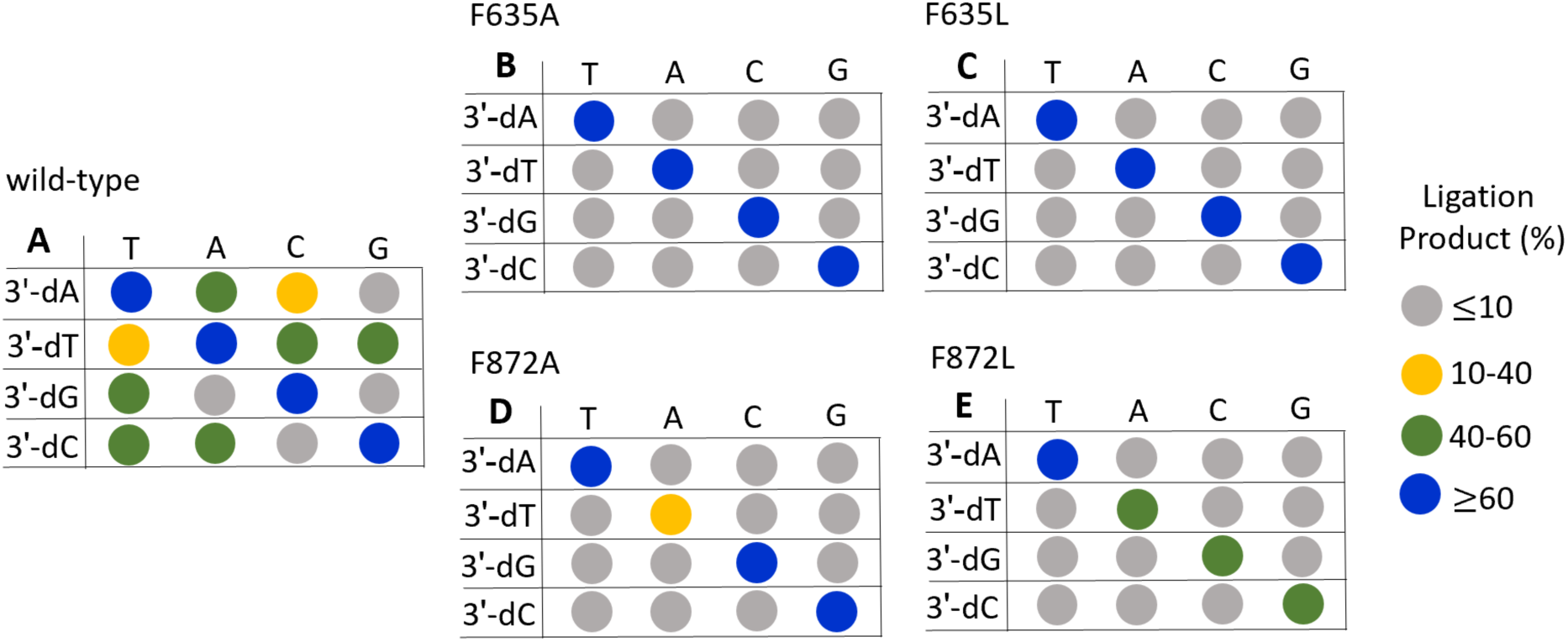
Summary of the ligation efficiencies of LIG1 wild-type, F635 and F872 mutants for nick DNA substrates containing all 12 non-canonical mismatches. **(A-E)** The amount of ligation products (%) for 3’-mismatches are shown for 3 min time point of ligation reaction for LIG1 wild-type (A) and active site mutants F635A (B), F635L (C), F872A (D), and F872L (E) as shown in Figures 1-5.

**Figure 22.**
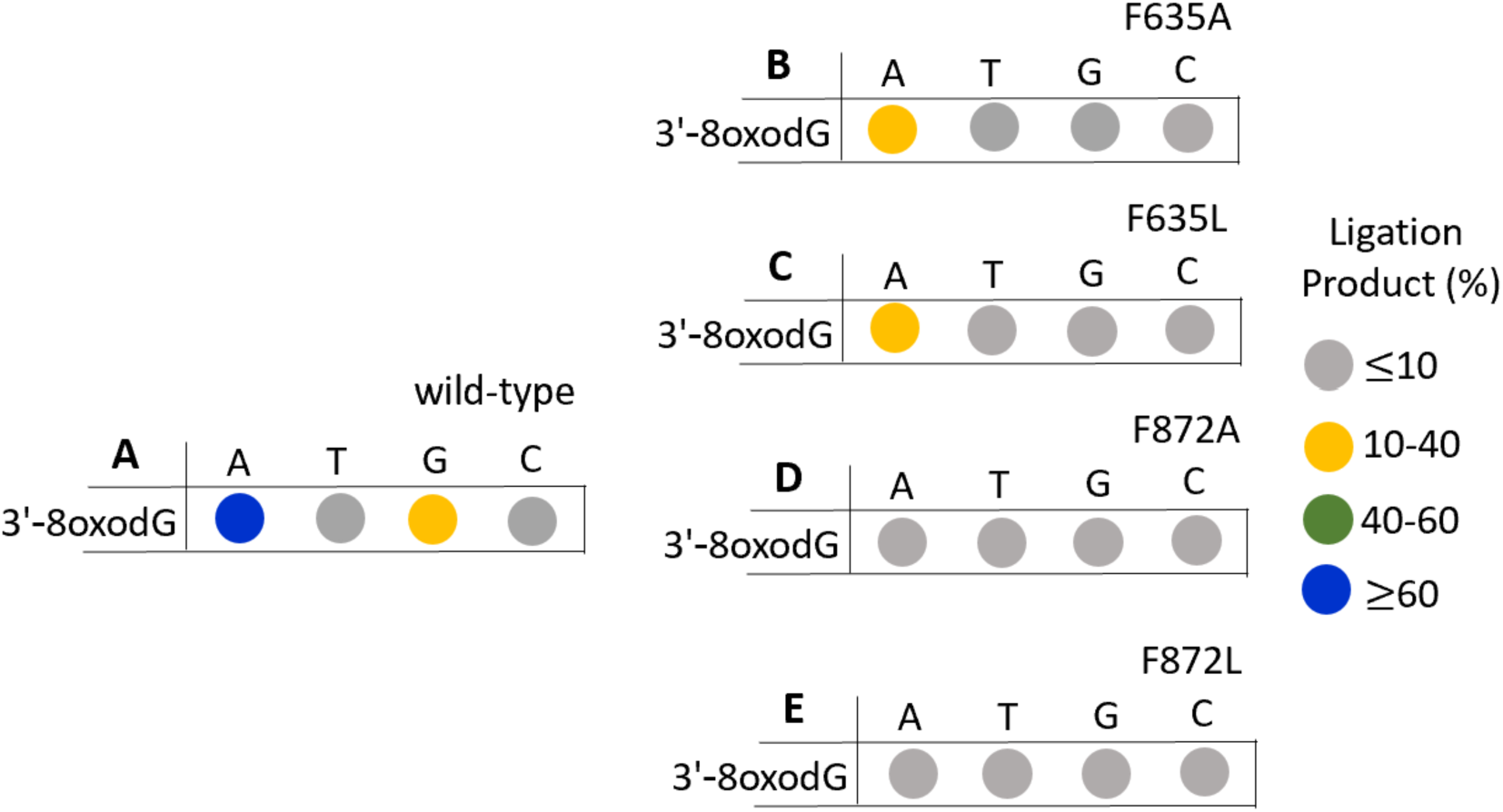
Summary of the ligation efficiencies of LIG1 F635 and F872 mutants for nick DNA substrates containing damaged ends. **(A-E)** The amount of ligation products (%) for nick DNA substrates containing 3’-8oxodG are shown for 3 min time-\ point of ligation reaction for LIG1 wild-type (A) and active site mutants F635A (B), F635L (C), F872A (D), and F872L (E) as shown in Figures 6-10.

**Figure 23.**
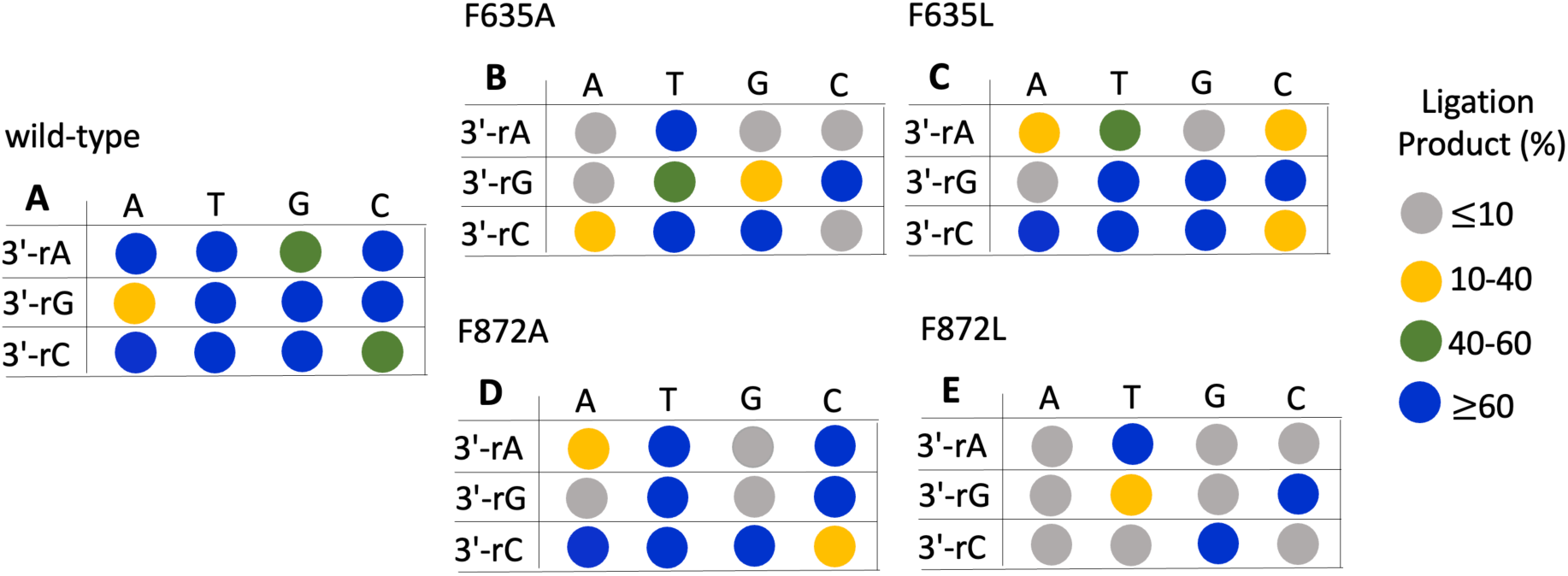
Summary of the ligation efficiencies of LIG1 F635 and F872 mutants for nick DNA substrates containing a single ribonucleotide at the 3’-end. **(A-E)** The amount of ligation products (%) for nick DNA substrates containing 3’-ribonucleotide are shown for 3 min time point of ligation reaction for LIG1 wild-type (A) and active site mutants F635A (B), F635L (C), F872A (D), and F872L (E) as shown in Figures 11-15.

**Figure 24.**
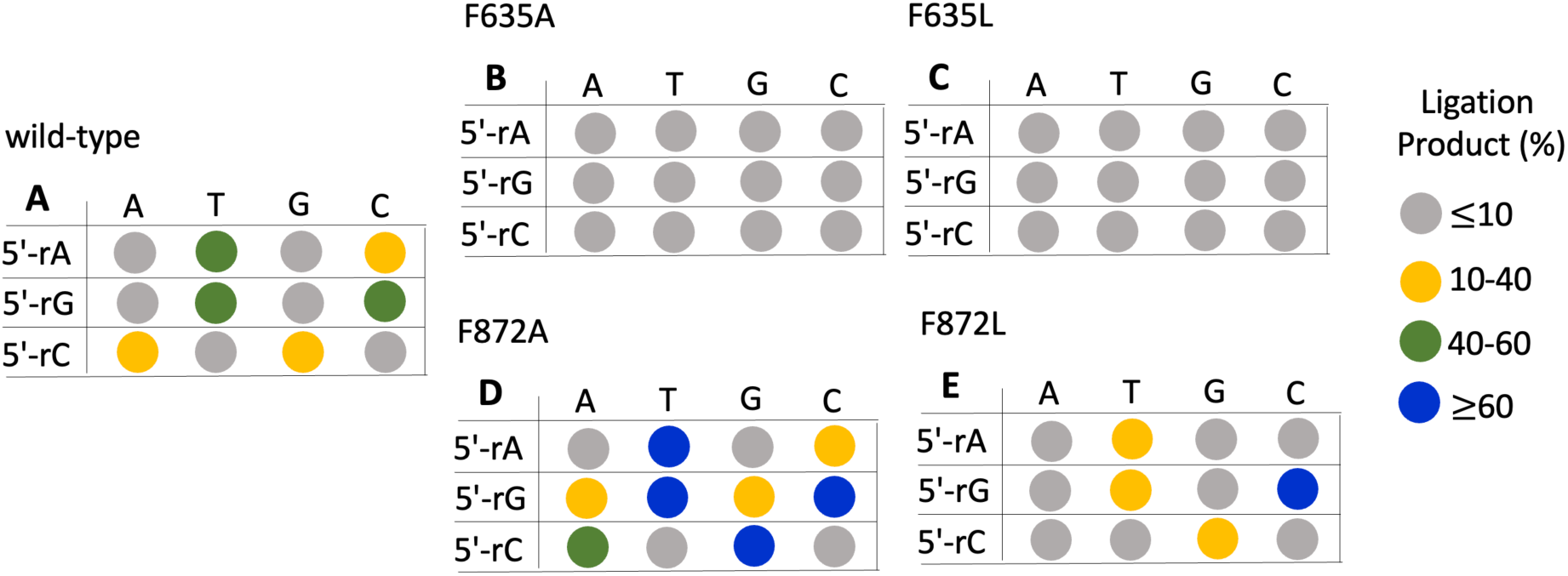
Summary of the ligation efficiencies of LIG1 F635 and F872 mutants for nick DNA substrates containing a single ribonucleotide at the 5’-end. **(A-E)** The amount of ligation products for nick DNA substrates containing 3’-ribonucleotide are shown for 15 min time point of ligation reaction for LIG1 wild-type (A) and active site mutants F635A (B), F635L (C), F872A (D), and F872L (E) as shown in Figures 16-20.

### Structures of LIG1 F635A and F872A reveal the importance of DNA end alignment

In addition to comprehensive investigation of ligation efficiency by LIG1 active site mutants for the nick DNA substrates containing all 12 non-canonical mismatches, ribonucleotides, and damaged ends (Figures 1-24), in the present study, we determined structures of LIG1 active site mutants, F635A and F872A, to understand how these mutations could affect the ligase/nick interactions at 3’- and 5’-ends (Figure 25A-B).

**Figure 25.**
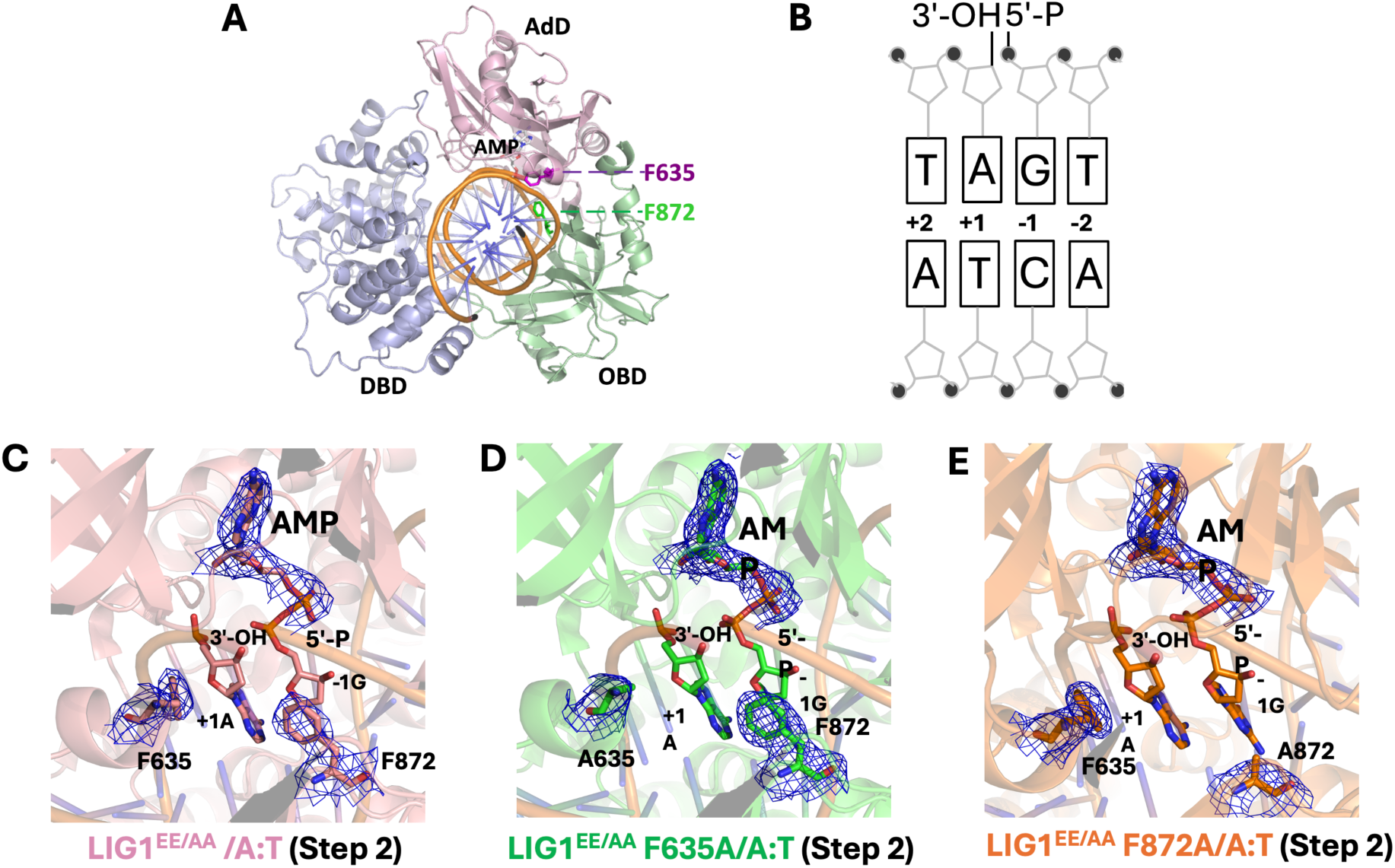
Structures of LIG1 F635A and F872A bound to the nick DNA duplex containing A:T at 3’-strand. **(A)** Structure of LIG1 shows that the catalytic core consisting of Adenylation (AdD) and Oligonucleotide-binding (OBD) domains as well as DNA-binding domain (DBD) of the protein encircle the nick DNA. **(B)** Scheme shows the sequence of nick DNA substrate used in LIG1 crytsalization **(C-E)** Structures of LIG1 EE/AA (C), F635A (D), and F872A (E) in complex with nick DNA containing a cognate A:T show step 2 of the ligation reaction where AMP is bound to 5’-PO_4_ end of nick. The 2Fo - Fc density map of the AMP, F635/A635, and F872/A872 are contoured at 1.5σ (blue). LIG1 is shown in cartoon mode and DNA, AMP, active site residues are shown in stick mode. LIG1^EE/AA^ structure was previously reported (PDB:7SUM)

In the structure of LIG1/A:T, we observed that the adenylate (AMP) moiety is transferred to the 5’-end of the nick where the DNA-AMP intermediate is formed, which refers to step 2 of ligation reaction (Figure 25C) as previously reported (36). Similarly, in the structures of LIG1 F635A and F872A mutants, we observed the ligase active site engaging with A:T during step 2 of the ligation reaction (Figure 25D-E). The overlay of LIG1 structures demonstrate the architecture of interactions at nick site does not show any difference (Supplementary Figure 22). We then analyzed the positions of LIG1 active site residues relative to the nucleotides around nick site (Figure 26). In the structure of LIG1 carrying F635 and F872, the distances to nucleotides at 3’- (+1A) and 5’- (−1G) ends are measured as 3.2 Å and 3.4Å, respectively (Figure 26A). However, in the structures of LIG1 F635A and F872A, the distances between the mutated site chains (A635 and A872) are increased and moved to 6.5Å and 4.9Å relative to 3’- and 5’-ends of nick, respectively (Figure 26B-C).

**Figure 26.**
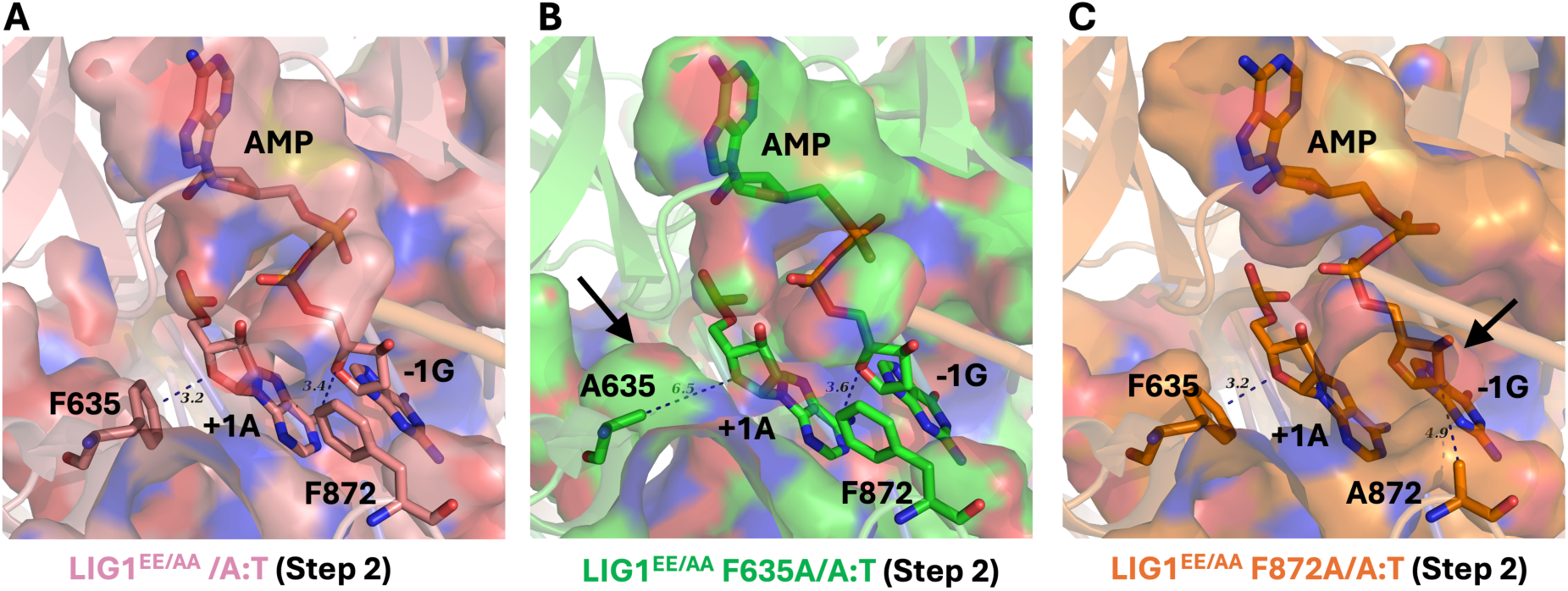
LIG1 structures reveal the importance of F635 and F872 residues for DNA end alignment. **(A-C)** LIG1 structures show differences in the distances between the active site residues (F635 or A635 and F872 or A872) relative to the nucleotides at 3’- (+1A) and 5’- (−1G) ends of nick site (arrows).

### Nick DNA binding by LIG1 F635A and F872A mutants at single-molecule level

We investigated nick sealing efficiency of LIG1 full-length wild-type and active site mutants for canonical nick and obtained ∼2- and ∼5-fold difference in the amount of ligation products by F635A and F872A mutations, respectively, when compared to the wild-type enzyme (Figure 27). As we recently reported for LIG1 wild-type (37), using TIRF microscopy and the AF488-labeled dsDNA containing a single nick site and Cy5-labeled full-length LIG1, we employed single-molecule fluorescence co-localization approach to further investigate the impact of the mutations at the LIG1 active site residues F635 and F872 on the nick DNA binding in real-time (Figure 28A).

**Figure 27.**
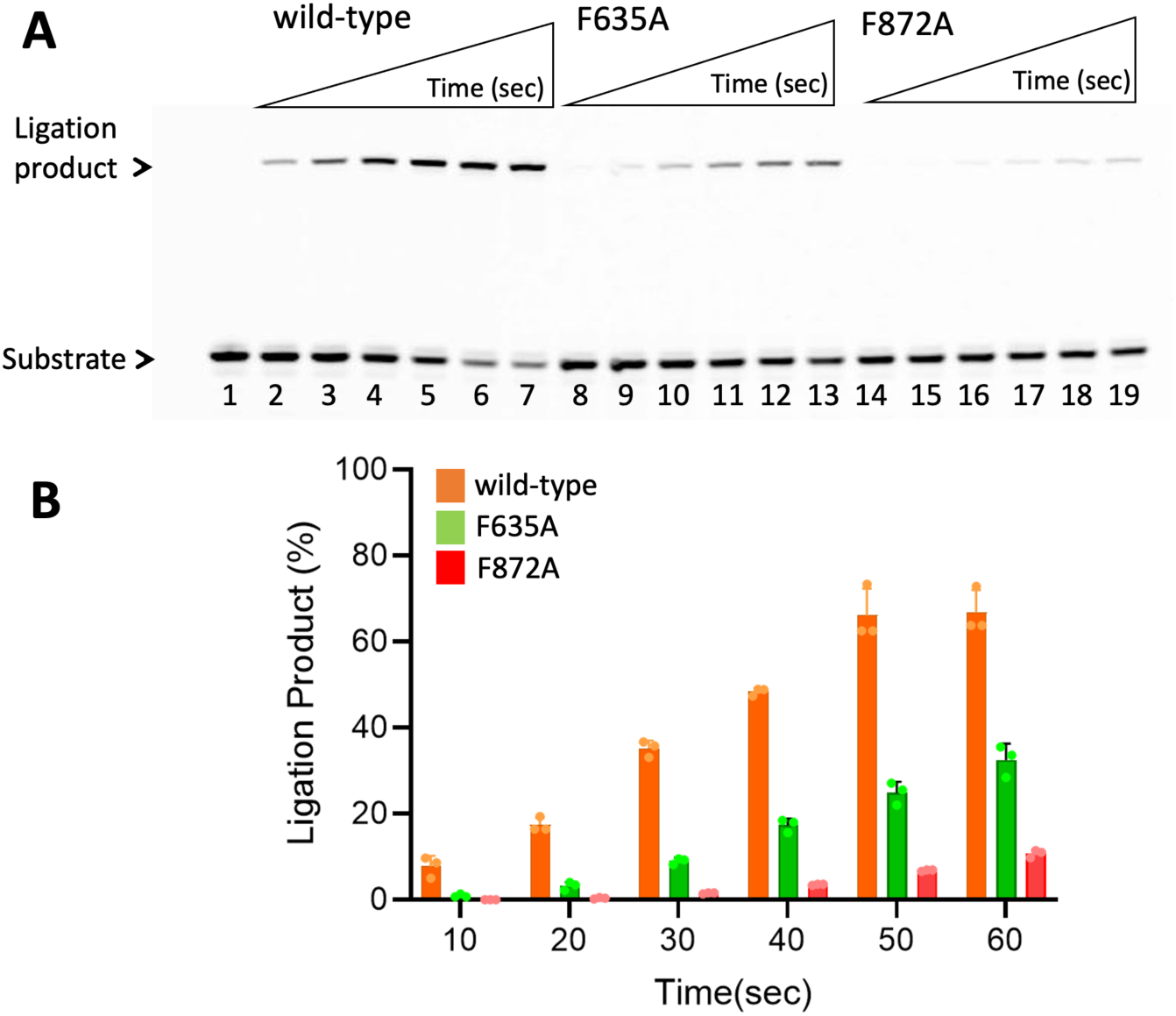
Nick sealing efficiency of LIG1 active site mutants in the presence of nick DNA with canonical end. (**A**) Lane 1 is the negative enzyme control of nick DNA substrate with 3’-dA:T. Lanes 2-7, 8-13, and 14-19 are the ligation products by LIG1 wild-type, F635A and F872A, respectively, and correspond to time points of 10, 20, 30, 40, 50, and 60 sec. (**B**) Graph shows time-dependent changes in the amount of ligation products. The data represent the average from three independent experiments ± SD.

**Figure 28.**
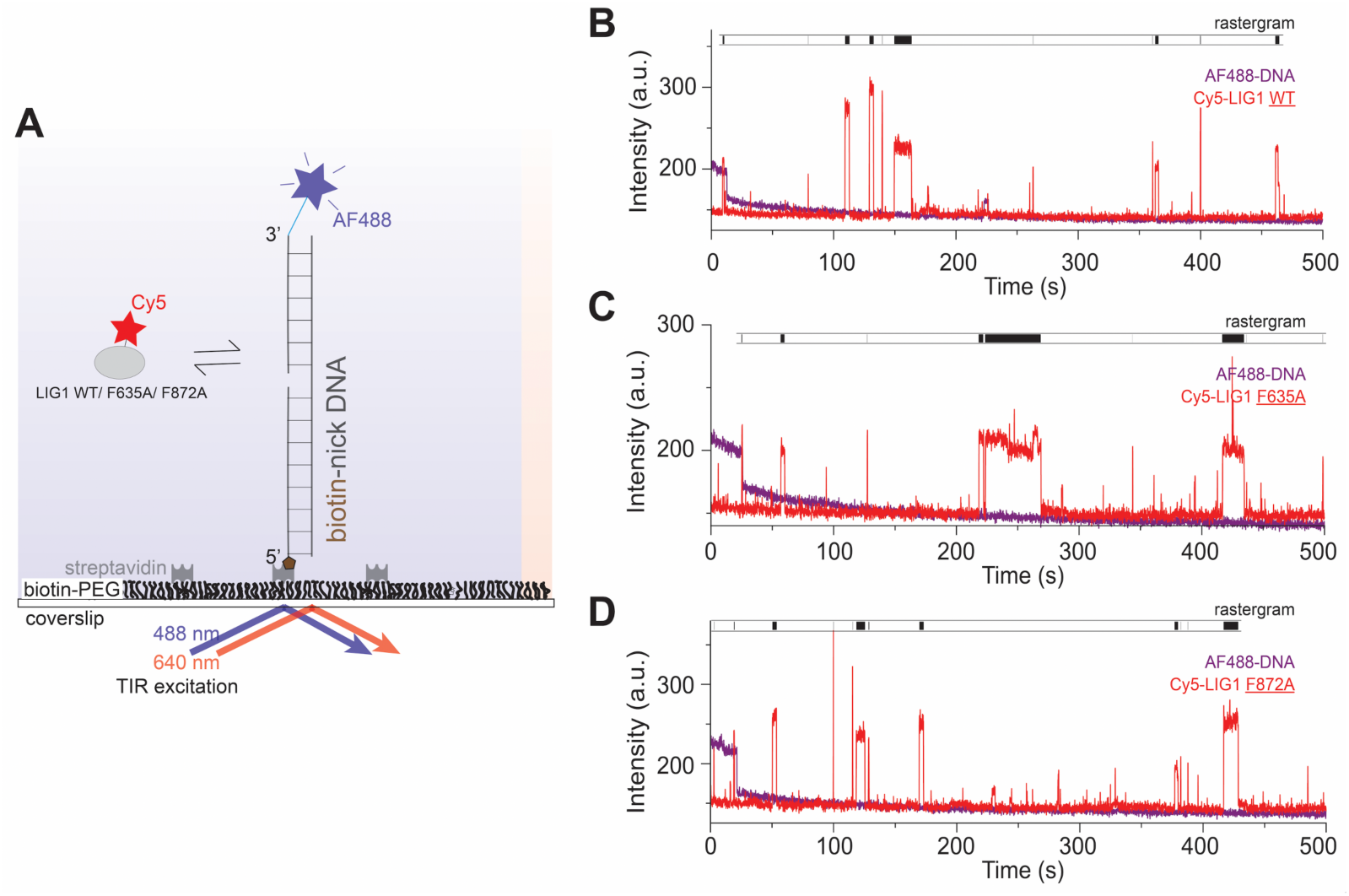
Single-molecule characterization of nick DNA binding by LIG1 active site mutants. **(A)** Scheme shows that AF488-labeled dsDNA with a single nick site immobilized on PEG-coated, biotinylated slide surface and imaged with a TIRF microscope to monitor real-time Cy5-labeled LIG1/nick DNA binding. **(B-D)** Fluorescence intensity *versus* time traces show repeated protein binding events for LIG1 wild-type and active site mutants.

LIG1^Cy5^ binding to the DNA was identified by the co-localization of AF488 and Cy5 fluorescence signals within a diffraction-limited spot. Analyses of individual fluorescence time trajectories show repeated transient Cy5 co-localizations with AF488, indicating dynamic binding of LIG1 wild-type and active site mutants on the DNA (Figure 28B-D). Rastergrams generated from several individual time traces idealized by hidden Markov model (HMM) showed dynamic binding behavior for the LIG1 proteins to the nick DNA, as indicated by both short and long-lived binding events (Figure 29A-C). From the dwell times distributions in the bound states, we next estimated the average binding lifetimes. For LIG1 wild-type, consistent with our recent study (37), we observed two populations with average binding lifetimes of 0.7 sec (60%) and 6 sec (40%) (Figure 29D). We demonstrated that the long binding events are due to formation of stable nick-bound LIG1-DNA complex. Interestingly, we observed no significant differences in the nick binding mode as depicted in the rastergrams or in binding lifetimes for the LIG1 mutants F635A and F872A, suggesting that these active site mutations do not affect the LIG1 binding to nick sites on DNA.

**Figure 29.**
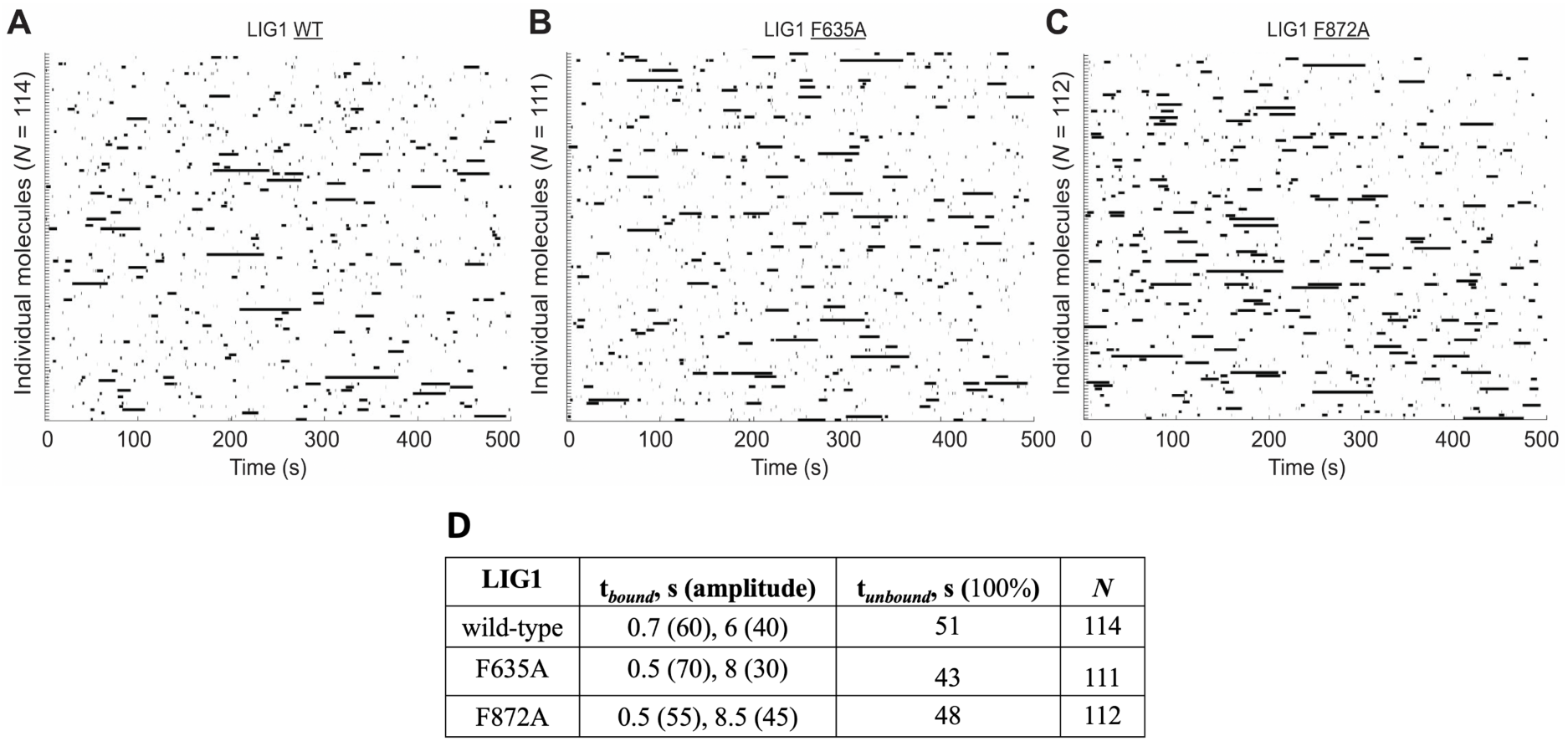
LIG1 active site mutants exhibit similar nick DNA binding characteristics with wild-type enzyme. **(A-C)** Rastergrams of randomly selected traces are shown for LIG1 proteins displaying the distinct nick DNA binding behavior. **(D)** Table shows difference in lifetimes of protein-bound states for *t*_bound_ and *t_un_*_bound_ times extracted for LIG1 wild-type, F635A and F872A mutants.

## Discussion

Nick sealing at the final step of almost all DNA repair mechanisms and during nuclear replication by DNA ligases is an essential chemical reaction for the maintenance of overall genome stability (5). Therefore, understanding the mechanism of DNA ligation in detail is critical to elucidating how cells can prevent the deleterious effects of single strand breaks, which arise from both endogenous processes and environmental factors. While functional and structural studies of human and non-human ligases have led to better understanding of the mechanism of ligation (7–22), much is still unknown about the contribution of ligase active site residues for ligation of a range of nick substrates containing canonical, mismatches, damaged, and ribonucleotide-containing ends as well as for DNA end alignment and nick binding affinity.

In the present study, we investigated the impact of the mutations at highly conserved active site phenylalanine residues, F635 or F872 of LIG1, on the efficiency of the ligation for 70 nick DNA substrates that mimic mismatches or damaged nucleotides that can be incorporated by DNA polymerases during nuclear replication and DNA repair. We found that DNA ligation by LIG1 F635A/L and F872A/L mutants is less efficient overall in the presence of all possible 12 mismatches, while nick sealing of canonical substrates is only mildly impacted. Our results demonstrated the mutagenic ligation efficiency of 3’-8oxodG:A by LIG1 wild-type is reduced by the mutations at the conserved active site residues. F635 mutants were capable of ligation while F872 mutants were not able to seal damaged ends. Furthermore, our results demonstrated an overall reduction in ligation efficiency by all four ligase active site mutants in the presence of a ribonucleotide at the 3’-end of the nick when compared to LIG1 wild-type. Specifically, F635A and F872L mutants showed the most drastic reduction, indicating an importance of 3’-ribonucleotide:template base pairing architecture. The increased flexibility of the ends allowed by these mutations meant that nucleophilic attack may not occur if there is weak or no base pairing between the 3’-terminus and template base due to the misalignment. This agrees with our observation showing a complete deficiency in ligation for all 12 non-canonical mismatches. Thus, in the presence of a 3’-ribonucleotide, F635 and F872 could cooperate to reduce the flexibility of their respective nick ends, which could eliminate the reliance on base pairing for proper alignment of nick ends. We also showed that the mutations at F635 and F872 result in a further decrease in nick sealing of 5’-ribonucleotides regardless of base pairing architecture with an exception of LIG1 F872A mutant showing an increase in ligation efficiency when compared to wild-type.

Our previously reported LIG1 structures in complex with nick DNA containing non-canonical ends demonstrated how the ligase active site engages with aberrant nicks at atomic resolution during the three-step ligation reaction (34–36). We reported that LIG1 deters A:C and favors G:T mismatches at steps 1 and 2 of the ligation reaction, respectively (36). Furthermore, in our previously solved structures of LIG1/nick DNA with 3’-ribonucleotide, we captured the ligase active site at step 3 of ligation reaction where a final phosphodiester bond is formed (34). These structures revealed that the 3’-ribonucleotide adopts 3’-endo conformation, D570 and R871 residues interact with 3’-OH of nick and 2’-OH of the ribose, respectively, and coordinate with water molecules, which overall enables the ligase to accommodate “wrong” sugar at 3’-end of nick efficiently (34). Furthermore, we previously reported in the structures of LIG1/RNA-DNA junction harboring 5’-ribonucleotide that the ligase stays adenylated during initial step 1 of ligation reaction and cannot move forward with an adenylate transfer due to a large conformational change downstream the nick resulting in a shift at Arg(R)738 residue in the Adenylation domain of the ligase, demonstrating a diminished ligation and proficient sugar discrimination against nicks with 5’-ribonucleotide (35).

In the present study, the crystal structures of LIG1 mutants F635A and F872A further provided an atomic insight into the mechanism of nick sealing, particularly, the role of conserved active site residues to DNA end alignment for proper catalysis. We suggest that the differences in distance between the active site residues relative to nucleotides at 3’- and 5’-ends lead to the formation of cavities in the absence of aromatic side chains at these positions, which enables an increased flexibility around nick site. Therefore, this structural adjustment due to F to A mutations at these critical active site residues could confound DNA end alignment which is required for a proper catalysis. All together, these results collectively support that LIG1 active site residues F635 and F872 play an important role in reducing the flexibility of the 3’- and 5’-ends of the nick, respectively, to ensure proper alignment for in-line nucleophilic attack, and can affect the nick sealing efficiency (Supplementary Scheme 1A). Further structure/function studies with those active site mutants with nicks containing mismatches or ribonucleotides are required to better understand the importance of DNA end alignment for ligase fidelity at atomic resolution. Because the ligation reaction mechanism is common in ATP-dependent ligases and both F635 and F872 residues are conserved (Supplementary Scheme 1B), their mutation in other human and non-human DNA ligases could potentially show similar effects on ligation. Finally, our single-molecule characterization of LIG1 F635A and F872A mutants demonstrated similar nick DNA binding modes with the wild-type protein, which suggests that the mutations at these active site residues do not affect initial DNA binding and further steps of ligation reaction that requires 3’- and 5’-ends to be in an alignment could be more sensitive to the absence of aromatic side chains.

Overall, our study contributes to a comprehensive understanding of how LIG1 maintains faithful nick sealing while engaging with unusual ends and how the mutations at conserved active sites, F635 and F872, impact ligation efficiency for a diverse range of the base pairing architectures that mimic DNA polymerase mediated canonical, mismatch, damaged, and ribonucleotide misincorporation products. The study could provide an insight into the conserved mechanism by which DNA ligases can efficiently repair broken strand breaks during nuclear replication and final steps of DNA repair pathways.

## Experimental Procedures

### Protein purifications

We generated Phe(F) to Ala(A) and Leu(L) amino acid substitutions at residues F635 and F872 of LIG1 that reside in the AdD and the OBD domains of the ligase, respectively, within the catalytic core of the enzyme (Supplementary Figure 1A). LIG1 C-terminal (△261) wild-type, F635A, F635L, F872A, and F872L mutant proteins with his-tag were purified as described previously (27–31). Briefly, the proteins were overexpressed in *E. coli* (DE3) cells and grown in Terrific Broth (TB) media with kanamycin (50 μgml^−1^) and chloramphenicol (34 μgml^−1^) at 37 °C. Once the OD_600_ reached 1.0, the cells were induced with 0.5 mM isopropyl β-D-thiogalactoside (IPTG) and overexpression continued overnight at 20 °C. After centrifugation, cells were lysed in lysis buffer containing 50 mM Tris-HCl (pH 7.0), 500 mM NaCl, 20 mM imidazole, 2 mM β-mercaptoethanol, 5% glycerol, and 1 mM phenylmethylsulfonyl fluoride (PMSF) by sonication at 4 °C. The lysate was pelleted at 31,000 x g for 90 min at 4 °C. The cell lysis solution was clarified and then loaded onto a HisTrap HP column that was previously equilibrated with binding buffer containing 50 mM Tris-HCl (pH 7.0), 500 mM NaCl, 20 mM imidazole, 2 mM β-mercaptoethanol, and 5% glycerol. The column was washed with binding buffer and then eluted with an increasing imidazole gradient 20-500 mM at 4 °C. The collected fractions were then subsequently loaded onto a HiTrap Heparin column that was equilibrated with binding buffer containing 20 mM Tris-HCl (pH 7.0), 50 mM NaCl, 2 mM β-mercaptoethanol, and 5% glycerol. The protein was eluted with a linear gradient of NaCl up to 1 M. The protein was further purified by Superdex 200 10/300 column in the buffer containing 50 mM Tris-HCl (pH 7.0), 200 mM NaCl, 1 mM DTT, and 5% glycerol. All proteins purified in this study were concentrated, aliquoted, and stored at −80 °C. LIG1 full-length (1-919 amino acids) wild-type, F635A and F872L proteins with his-tag were purified as described above for the LIG1 C-terminal proteins. Protein quality of LIG1 full-length and C-terminal proteins were evaluated on a 10% SDS-PAGE gel, and protein concentrations were measured using absorbance at 280 nm (Supplementary Figure 1).

### DNA ligation assays

Nick DNA substrates with a 6-carboxyfluorescein (FAM) label were used in DNA ligation assays. Nick DNA substrates containing Watson-Crick base paired ends, and all 12 non-canonical mismatches were prepared by annealing upstream oligonucleotides 3’-dA, dT, dG, or dC with template oligonucleotides A, T, G, or C (Supplementary Table 1). Nick DNA substrates containing oxidative DNA damage were prepared by annealing 3’-8oxodG with template oligonucleotides A, T, G, or C (Supplementary Table 2). Nick DNA substrates containing a single ribonucleotide at the 3’-end were prepared by annealing upstream oligonucleotides 3’-rA, 3’-rG, and 3’-rC with template oligonucleotides A, T, G, or C (Supplementary Table 3). Nick DNA substrates containing a single ribonucleotide at the 5’-end were prepared by annealing downstream oligonucleotides 5’-rA, 5’-rG, and 5’-rC with template oligonucleotides A, T, G, or C (Supplementary Table 4).

Ligation assays were performed to investigate the nick DNA substrate specificity of LIG1 wild-type and active site mutants F635A, F635L, F872A, F872L (Supplementary Scheme 2) as described (27–31). Briefly, the reaction was initiated by the addition of LIG1 (100 nM) to a mixture containing 50 mM Tris-HCl (pH 7.5), 100 mM KCl, 10 mM MgCl_2_, 1 mM ATP, 1 mM DTT, 100 µgml^-1^ BSA, 1% glycerol, and the nick DNA substrate (500 nM) in a final volume of 10 µl. The reaction mixture was incubated at 37 °C and stopped at the time points indicated in the figure legends by mixing with an equal volume of loading dye containing 95% formamide, 20 mM ethylenediaminetetraacetic acid, 0.02% bromophenol blue, and 0.02% xylene cyanol. Reaction products were separated by electrophoresis on an 18% Urea-PAGE gel, the gels were scanned with Typhoon PhosphorImager RGB, and the data was analyzed using ImageQuant software.

### Crystallization and structure determination

For the crystallization of LIG1 F635A and F872A mutants, we used the EE/AA mutant that harbors E346A and E592A mutations, resulting in the ablation of the high-fidelity site (referred to as Mg^HiFi^ site), which has been utilized in previous LIG1 structures with non-canonical substrates (34–36). We solved LIG1 structures in complex with nick DNA containing a canonical A:T (Supplementary Table 5). LIG1 (at 27 mgml^-1^)/DNA complex solution was prepared in the buffer containing 20 mM Tris-HCl (pH 7.0), 200 mM NaCl, 1 mM DTT, 1 mM EDTA, and 1 mM ATP at 1.4:1 DNA:protein molar ratio and then mixed with 1 μl reservoir solution. LIG1-nick DNA complex crystals were grown at 20 °C using the hanging drop method, harvested, and submerged in cryoprotectant solution containing reservoir solution mixed with glycerol to a final concentration of 20% glycerol before being flash cooled in liquid nitrogen for data collection. LIG1 crystals were obtained and kept at 100 °K during X-ray diffraction data collection using the beamline CHESS-7B2 and processed using HKL 2000 (HKL Research, Inc). All structures were solved by the molecular replacement method using PHASER with PDB entry 7SUM as a search model (38). Iterative rounds of model building were performed in COOT and the final models were refined with PHENIX (39,40). All structural images were drawn using PyMOL (The PyMOL Molecular Graphics System, V0.99, Schrödinger, LLC). Detailed crystallographic statistics are provided in Table 1.

**Table 1:**
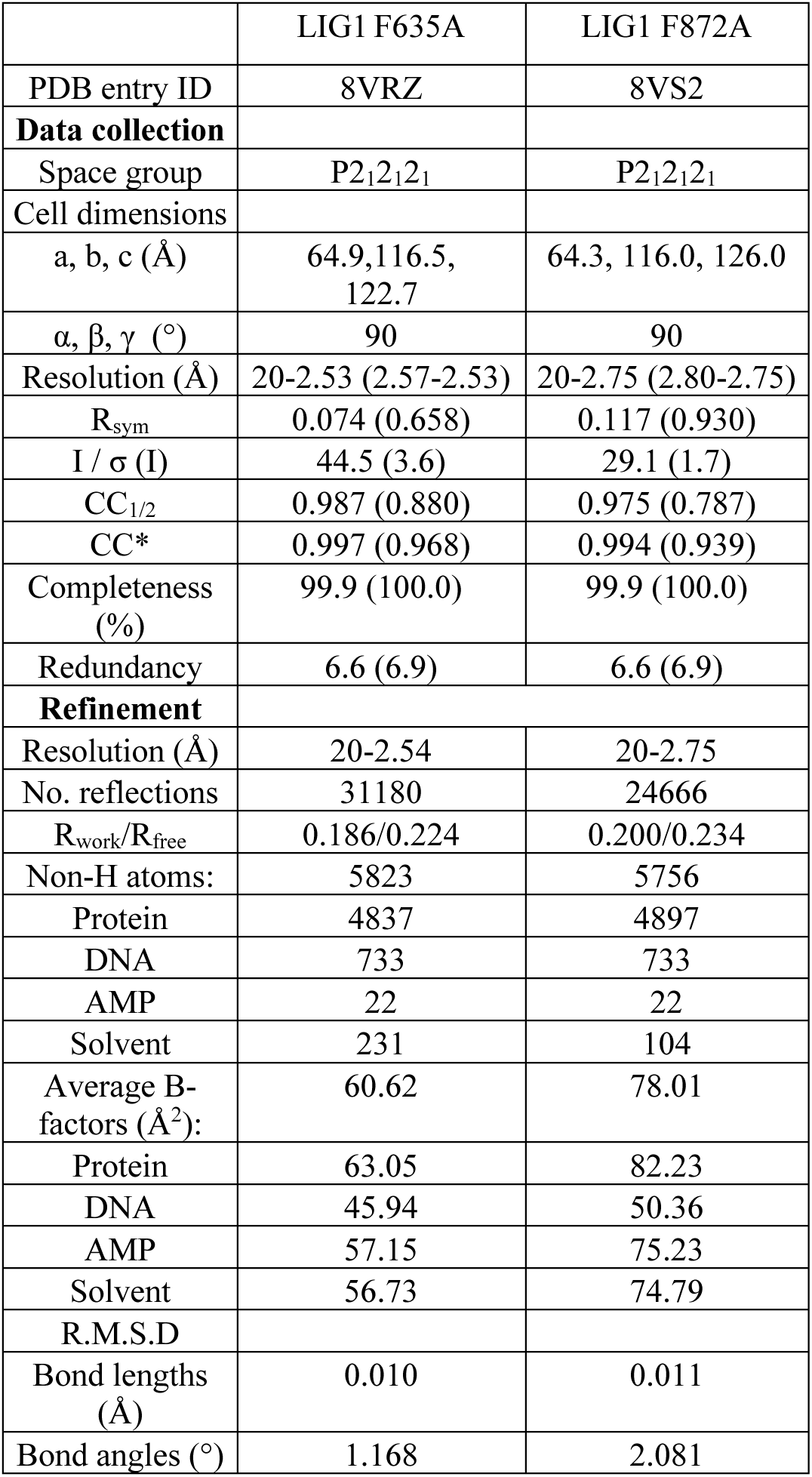
X-ray data collection and refinement statistics of LIG1 structures.

### Nick DNA binding of LIG1 active site mutants at single-molecule level

To visualize real-time LIG1-nick DNA binding, we performed single-molecule experiments using a total internal reflection fluorescence (TIRF) microscope (Nikon Eclipse Ti2-E) as reported (37). For this purpose, we fluorescently labeled LIG1 full-length wild type, F635A and F872A mutants using Amersham Cy5 monoreactive NHS ester dye (LIG1^Cy5^) pack (Cytiva). We used 3’-AF488-labeled dsDNA substrate (34-mer) containing a 5’-Biotin and a single nick site containing A:T (Supplementary Table 6). Labeling efficiencies (1.2-1.4) were determined by calculating the molar ratio of dye to protein by measuring absorbance at 280 nm and 649 nm. We finally pre-adenylated LIG1^Cy5^ proteins by incubating them in a buffer containing a final concentration of 10 mM ATP and then dialyzed against the protein storage buffer. Single-use aliquots of the fluorescently labeled and pre-adenylated LIG1 proteins were flash-frozen with liquid nitrogen and stored until use at −80 °C.

We performed single-molecule measurements as reported previously (37), Briefly, the glass coverslips were functionalized with a mixture of biotin-PEG-SVA and mPEG-SVA and microfluidic channels were set up using the passivated coverslips and washed three times with the T50 buffer containing 10 mM Tris-HCl (pH 8.0) and 50 mM NaCl. Streptavidin (0.2 mg/ml) in the T50 buffer was then flowed onto the slide, incubated with the biotin-PEG, and then washed with the T50 buffer. DNA substrate was diluted to a final concentration of 10 pM in the imaging buffer containing 1 mM HEPES (pH 7.4), 20 mM NaCl, 0.02% BSA (w/v), and 0.002% Tween 20 (v/v), flowed onto the slide, and allowed to incubate for immobilization. Excess unbound DNA substrate was then washed by the imaging buffer and the slide surface was passivated by incubating with 10 mg/ml BSA. Finally, LIG1^Cy5^ proteins (1 nM) in the imaging buffer was flowed onto the slide and allowed to equilibrate before imaging. Both AF488 and Cy5 dyes were simultaneously excited using 488 nm and 640 nm lasers, respectively. Emissions from two fluorophores were separated into two channels using a Cairn Optosplit III image splitter and simultaneously recorded at 100 ms time resolution using a Hamamatsu SCMOS camera. Locations of molecules and fluorophore intensity over time traces were extracted from the raw movie files using Nikon NIS-Elements analysis software (Nikon, version: AR 6.02.01) and selected time traces were idealized using a two-state hidden Markov model (HMM) for the unbound and bound states in QuB. Rastergrams summarizing several individual traces were generated from the individual trace HMMs using custom written MATLAB script. From the idealized traces, dwell times of the bound and unbound states were calculated using MATLAB. Cumulative frequency of the bound and unbound dwell-time distributions was plotted and fitted in Origin Lab (version 2024b) with single or double exponential functions to obtain the bound (t_bound_) and unbound (t_unbound_) states lifetimes.

## Data and material availability

Atomic coordinates and structure factors for the reported crystal structures of LIG1 have been deposited in the RCSB Protein Data Bank under accession numbers LIG1 F635A (8VRZ) and F872A (8VS2). All data are contained within the manuscript. Further information and requests of materials used in this research are available from the authors upon reasonable request and should be directed to Dr. Melike Çağlayan.

## Acknowledgement

This work is based upon research conducted at the Center for High Energy X-ray Sciences (CHEXS), which is supported by the National Science Foundation under award DMR-1829070, and the Macromolecular Diffraction at CHESS (MacCHESS) facility, which is supported by award 1-P30-GM124166-01A1 from the National Institute of General Medical Sciences, National Institutes of Health, and by New York State’s Empire State Development Corporation (NYSTAR).

## Author Contributions

M.Ç.: conceptualization; M.K.,M.G.,K.L.,D.A.,J.R.,E.C.,D.A: purified the proteins and performed the biochemical experiments. T.Q.: refined the structures; K.B.: prepared structure figures; S.C.: performed single-molecule experiments; M.K.,M.G.,K.B.,S.C.: writing-reviewing. M.Ç.: supervision, writing-reviewing, revisions, and editing. M.Ç.: funding.

