## Supplementary material for "Biochemical, structural, and single-molecule characterization of LIG1 active site mutants demonstrate role of F635 and F872 residues for faithful ligation": Supp Data

**Supplementary Information**

Supplementary Figures 1-22

Supplementary Schemes 1-2

Supplementary Tables 1-6

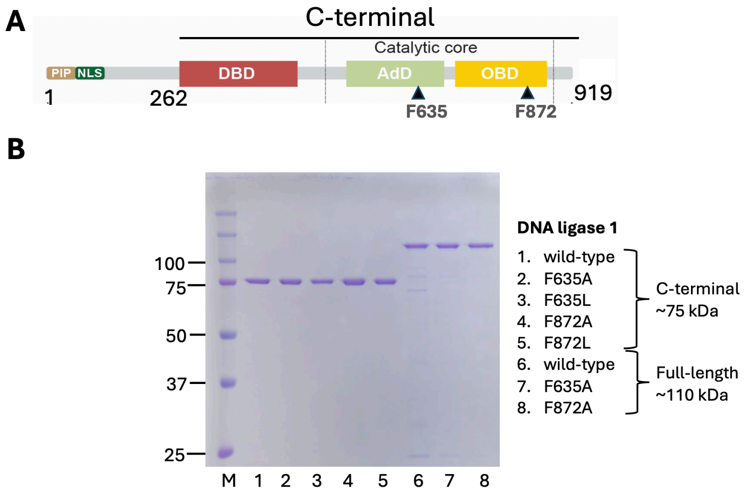


**Supplementary Figure 1. DNA ligase 1 active site mutants. (A)** The domain organization of DNA ligase 1 (LIG1) protein (1-919 amino acids) including N-terminal region (1-262 amino acids) and C-terminal domain that contains DNA-binding domain (DBD) and the catalytic core consisting of Adenylation (AdD) and Oligonucleotide-binding (OBD) domains. LIG1 active site residues F635 and F872 reside in AdD and OBD domains of the catalytic core, respectively. **(B)** Purified proteins of LIG1 C-terminal and full length proteins used in this study. M represents the protein marker.

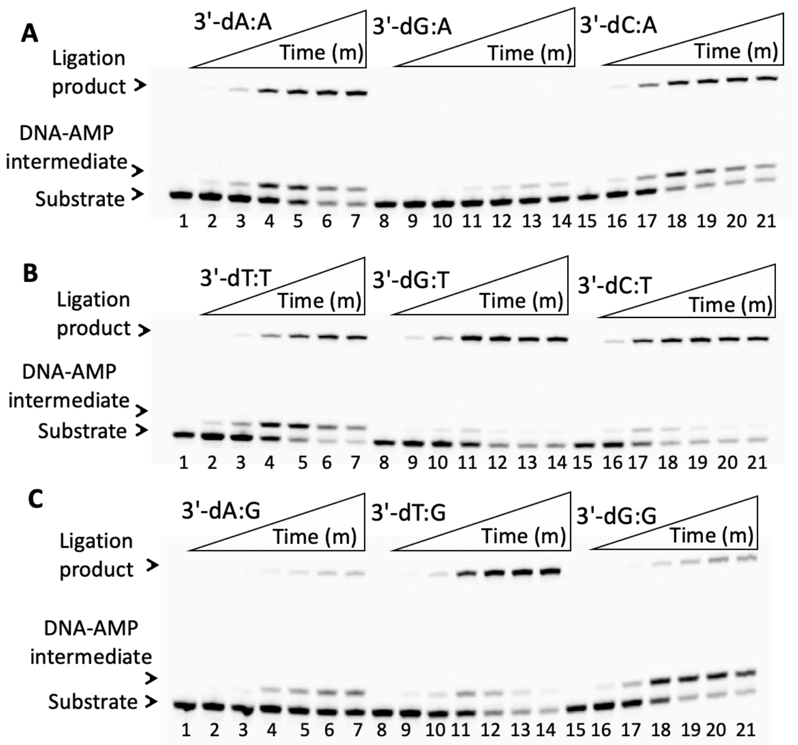


**Supplementary Figure 2. Ligation of nick DNA substrates containing template A, T, and G mismatches by LIG1 wild-type. (A-C)** Lanes 1, 8, and 15 are the negative enzyme controls of the nick DNA substrates containing template A, T, and G mismatches. Lanes 2-7, 9-14, and 16-21 are the ligation products in the presence of nick DNA substrates with 3'-mismatches by LIG1 wild-type, and correspond to time points of 0.5, 1, 3, 5, 8, and 10 min. Graphs showing the amount of ligation products for all 12 mismatches are presented in Figure 1.

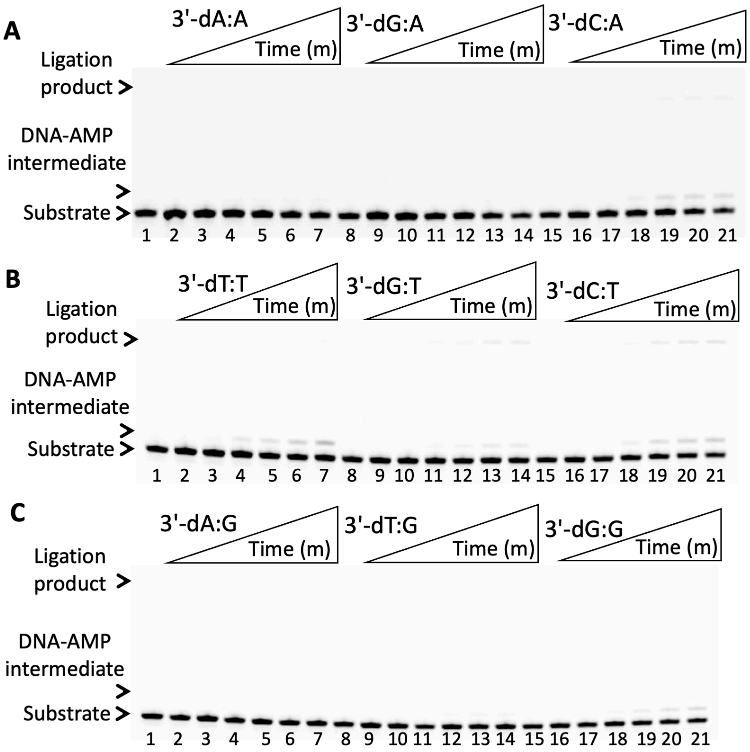


**Supplementary Figure 3. Ligation of nick DNA substrates containing template A, T, and G mismatches by LIG1 F635A. (A-C)** Lanes 1, 8, and 15 are the negative enzyme controls of the nick DNA substrates containing template A, T, and G mismatches. Lanes 2-7, 9-14, and 16-21 are the ligation products in the presence of nick DNA substrates with 3'-mismatches by LIG1 F635A, and correspond to time points of 0.5, 1, 3, 5, 8, and 10 min. Graphs showing the amount of ligation products for all 12 mismatches are presented in Figure 2.

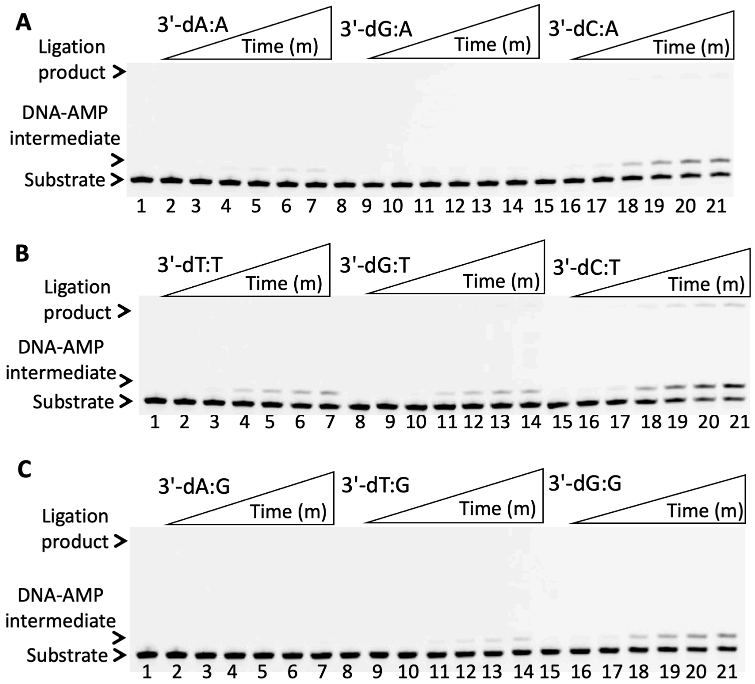


**Supplementary Figure 4. Ligation of nick DNA substrates containing template A, T, and G mismatches by LIG1 F635L. (A-C)** Lanes 1, 8, and 15 are the negative enzyme controls of the nick DNA substrates containing template A, T, and G mismatches. Lanes 2-7, 9-14, and 16-21 are the ligation products in the presence of nick DNA substrates with 3'-mismatches by LIG1 F635L, and correspond to time points of 0.5, 1, 3, 5, 8, and 10 min. Graphs showing the amount of ligation products for all 12 mismatches are presented in Figure 3.

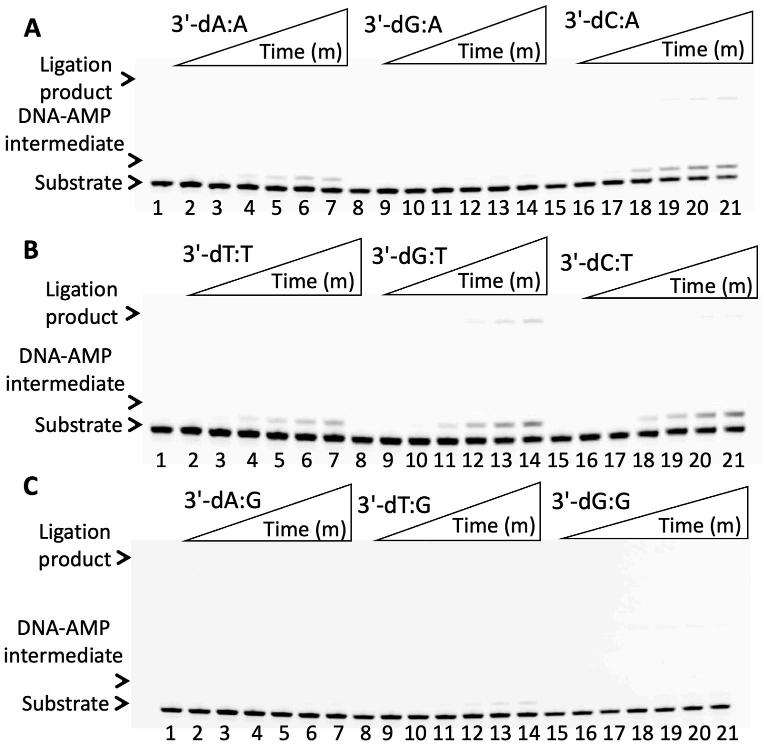


**Supplementary Figure 5. Ligation of nick DNA substrates containing template A, T, and G mismatches by LIG1 F872A. (A-C)** Lanes 1, 8, and 15 are the negative enzyme controls of the nick DNA substrates containing template A, T, and G mismatches. Lanes 2-7, 9-14, and 16-21 are the ligation products in the presence of nick DNA substrates with 3'-mismatches by LIG1 F872A, and correspond to time points of 0.5, 1, 3, 5, 8, and 10 min. Graphs showing the amount of ligation products for all 12 mismatches are presented in Figure 4.

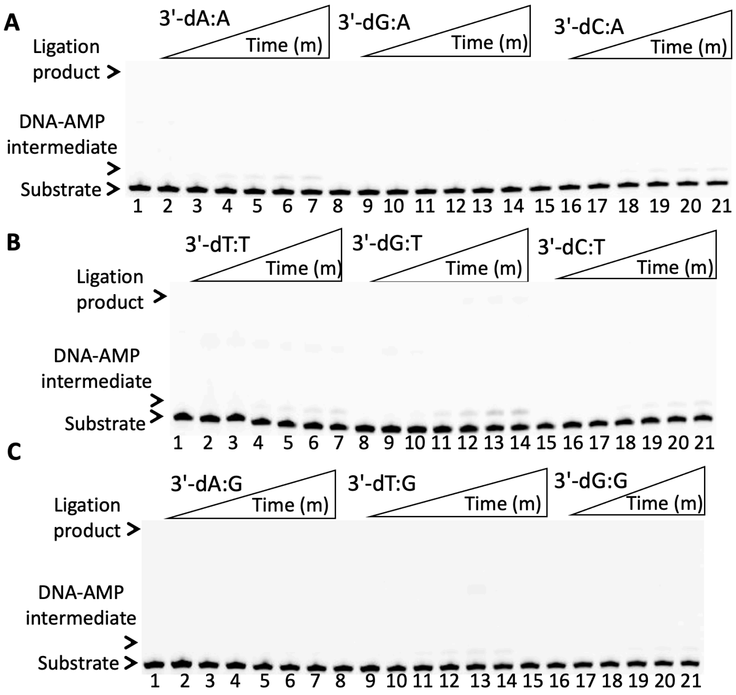


**Supplementary Figure 6. Ligation of nick DNA substrates containing template A, T, and G mismatches by LIG1 F872L. (A-C)** Lanes 1, 8, and 15 are the negative enzyme controls of the nick DNA substrates containing template A, T, and G mismatches. Lanes 2-7, 9-14, and 16-21 are the ligation products in the presence of nick DNA substrates with 3'-mismatches by LIG1 F872L, and correspond to time points of 0.5, 1, 3, 5, 8, and 10 min. Graphs showing the amount of ligation products for all 12 mismatches are presented in Figure 5.

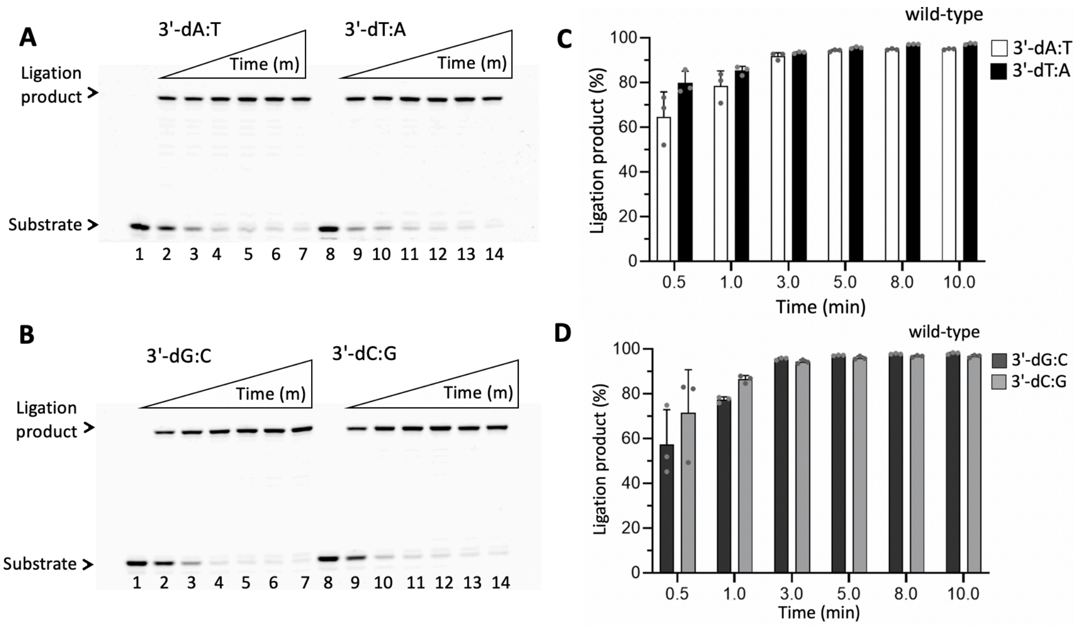


**Supplementary Figure 7. Ligation of nick DNA substrates containing canonical ends by LIG1 wild-type. (A-B)** Lanes 1 and 8 are the negative enzyme controls of the nick DNA substrates containing canonical ends. Lanes 2-7 and 9-14 are the ligation products in the presence of nick DNA substrates with 3'-dA:T, 3'-dT:A, 3'-dG:C, and 3'-dC:G by LIG1 wild-type, and correspond to time points of 0.5, 1, 3, 5, 8, and 10 min. **(C-D)** Graphs showing the amount of ligation products for all 4 canonical nick DNA substrates.

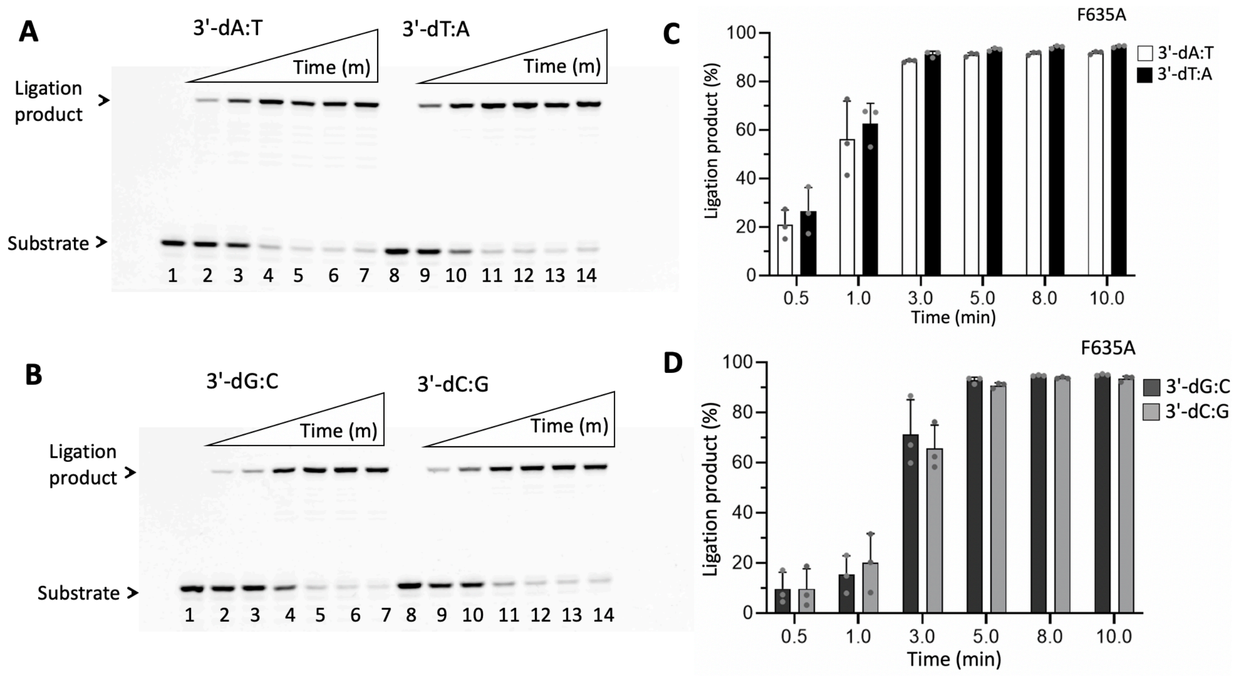


**Supplementary Figure 8. Ligation of nick DNA substrates containing canonical ends by LIG1 F635A. (A-B)** Lanes 1 and 8 are the negative enzyme controls of the nick DNA substrates containing canonical ends. Lanes 2-7 and 9-14 are the ligation products in the presence of nick DNA substrates with 3'-dA:T, 3'-dT:A, 3'-dG:C, and 3'-dC:G by LIG1 F635A, and correspond to time points of 0.5, 1, 3, 5, 8, and 10 min. **(C-D)** Graphs showing the amount of ligation products for all 4 canonical nick DNA substrates.

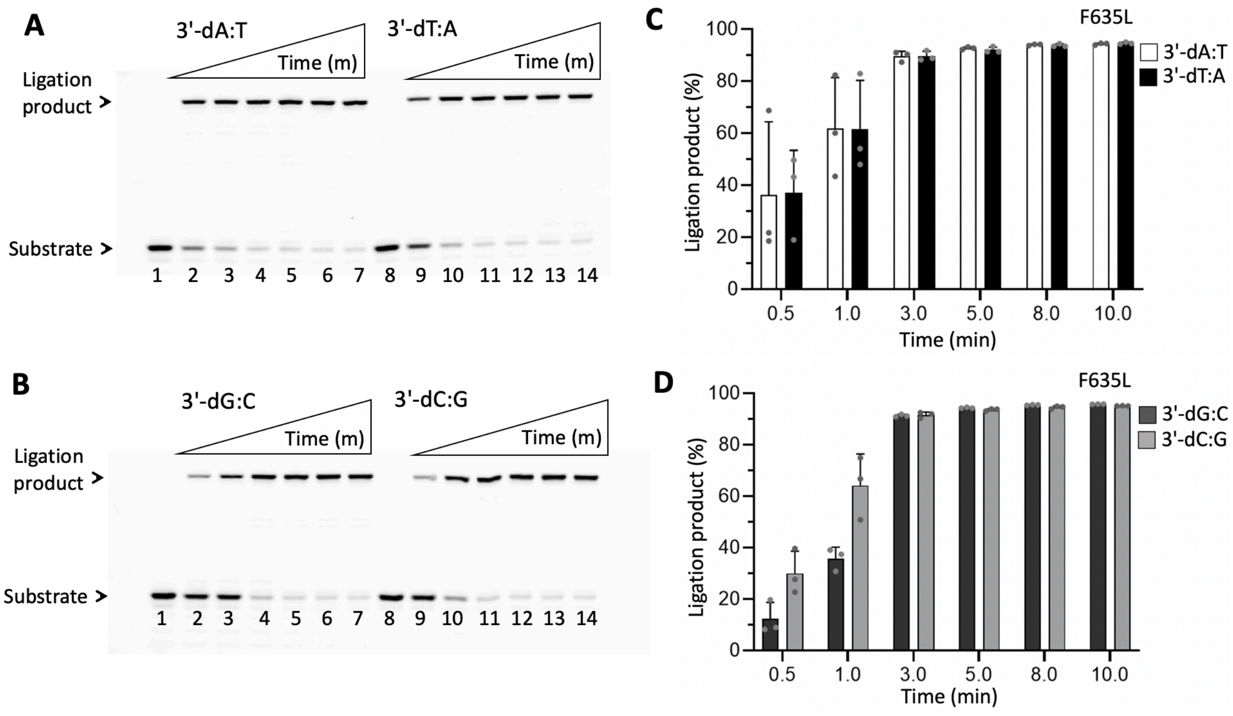


**Supplementary Figure 9. Ligation of nick DNA substrates containing canonical ends by LIG1 F635L. (A-B)** Lanes 1 and 8 are the negative enzyme controls of the nick DNA substrates containing canonical ends. Lanes 2-7 and 9-14 are the ligation products in the presence of nick DNA substrates with 3'-dA:T, 3'-dT:A, 3'-dG:C, and 3'-dC:G by LIG1 F635L, and correspond to time points of 0.5, 1, 3, 5, 8, and 10 min. **(C-D)** Graphs showing the amount of ligation products for all 4 canonical nick DNA substrates.

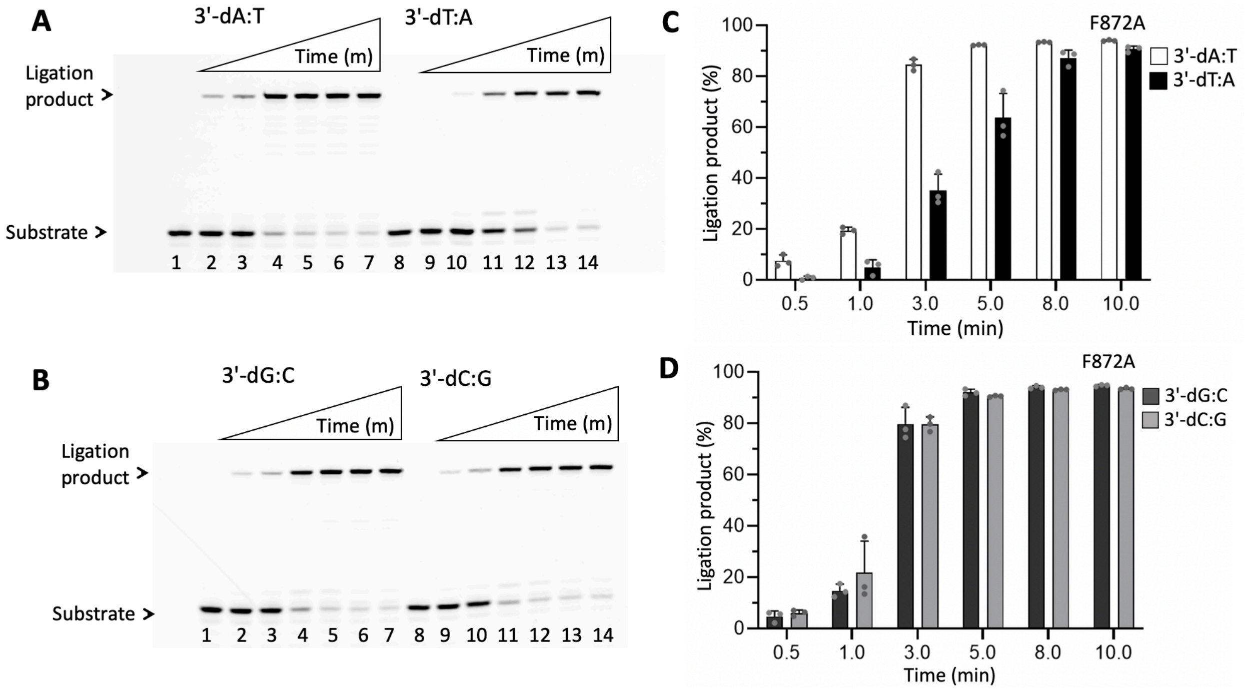


**Supplementary Figure 10. Ligation of nick DNA substrates containing canonical ends by LIG1 F872A. (A-B)** Lanes 1 and 8 are the negative enzyme controls of the nick DNA substrates containing canonical ends. Lanes 2-7 and 9-14 are the ligation products in the presence of nick DNA substrates with 3'-dA:T, 3'-dT:A, 3'-dG:C, and 3'-dC:G by LIG1 F872A, and correspond to time points of 0.5, 1, 3, 5, 8, and 10 min. **(C-D)** Graphs showing the amount of ligation products for all 4 canonical nick DNA substrates.

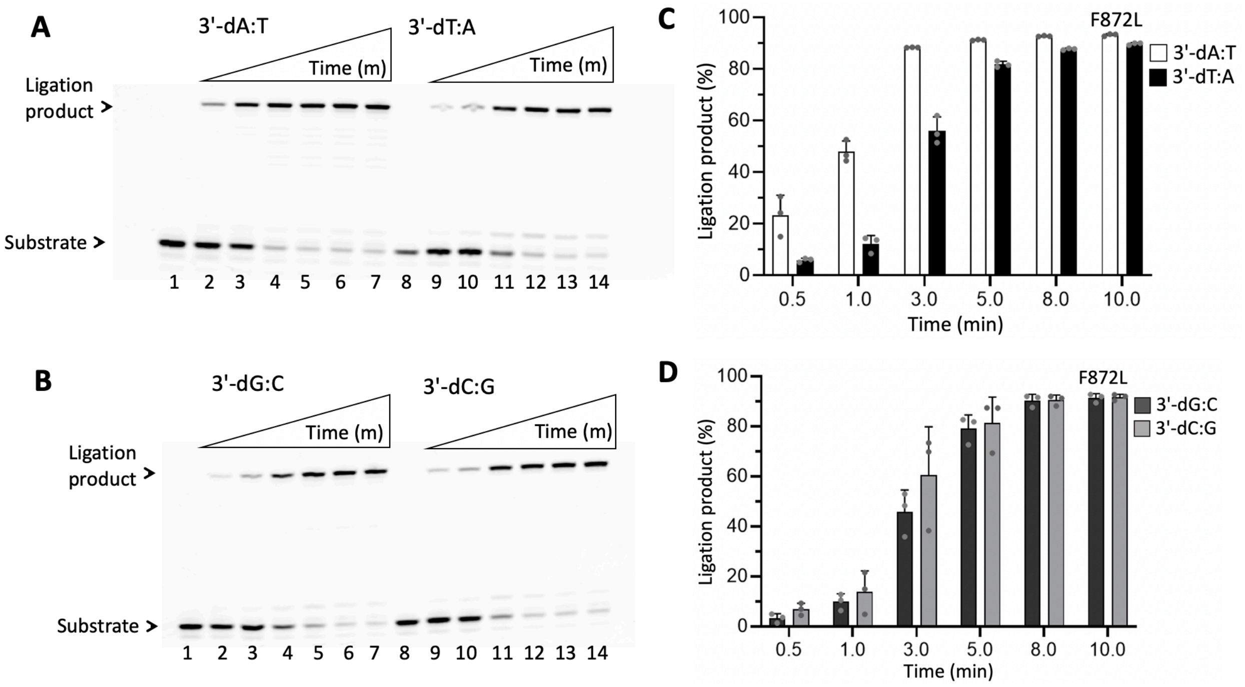


**Supplementary Figure 11. Ligation of nick DNA substrates containing canonical ends by LIG1 F872L. (A-B)** Lanes 1 and 8 are the negative enzyme controls of the nick DNA substrates containing canonical ends. Lanes 2-7 and 9-14 are the ligation products in the presence of nick DNA substrates with 3'-dA:T, 3'-dT:A, 3'-dG:C, and 3'-dC:G by LIG1 F872L, and correspond to time points of 0.5, 1, 3, 5, 8, and 10 min. **(C-D)** Graphs showing the amount of ligation products for all 4 canonical nick DNA substrates.

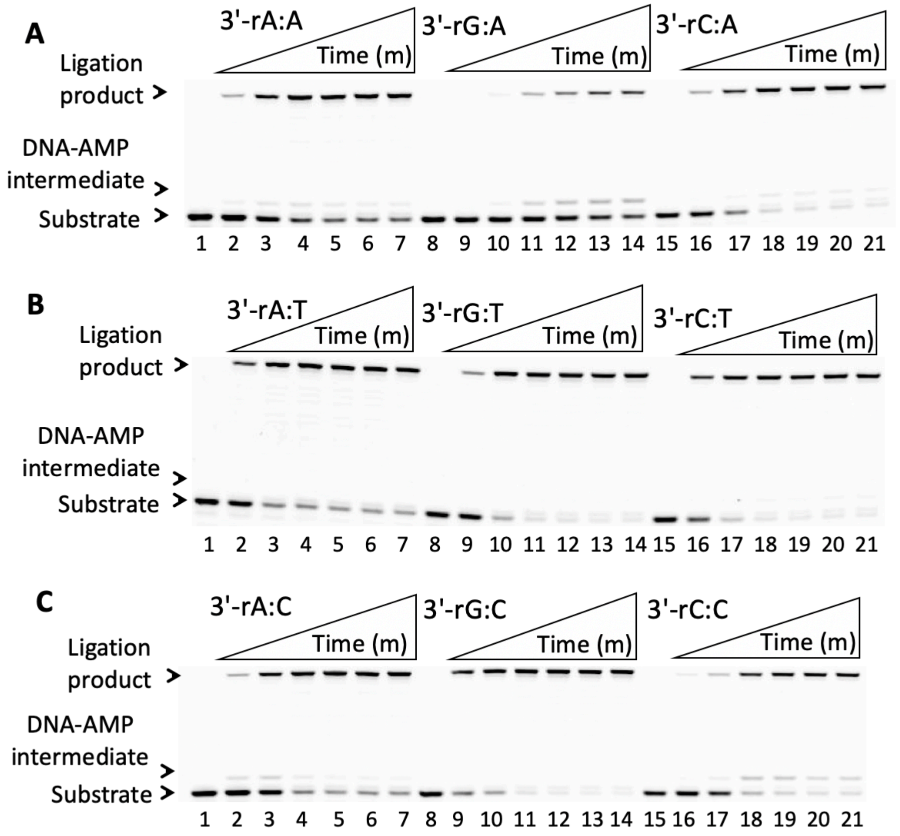


**Supplementary Figure 12. Ligation of nick DNA substrates containing 3'-ribonucleotide by LIG1 wild-type. (A-C)** Lanes 1, 8, and 15 are the negative enzyme controls of the nick DNA substrates containing template A, T, and C mismatches. Lanes 2-7, 9-14, and 16-21 are the ligation products in the presence of nick DNA substrates with a single ribonucleotide 3'-end by LIG1 wild-type, and correspond to time points of 0.5, 1, 3, 5, 8, and 10 min. Graphs showing time-dependent changes in the amounts of ligation products are presented in Figure 11.

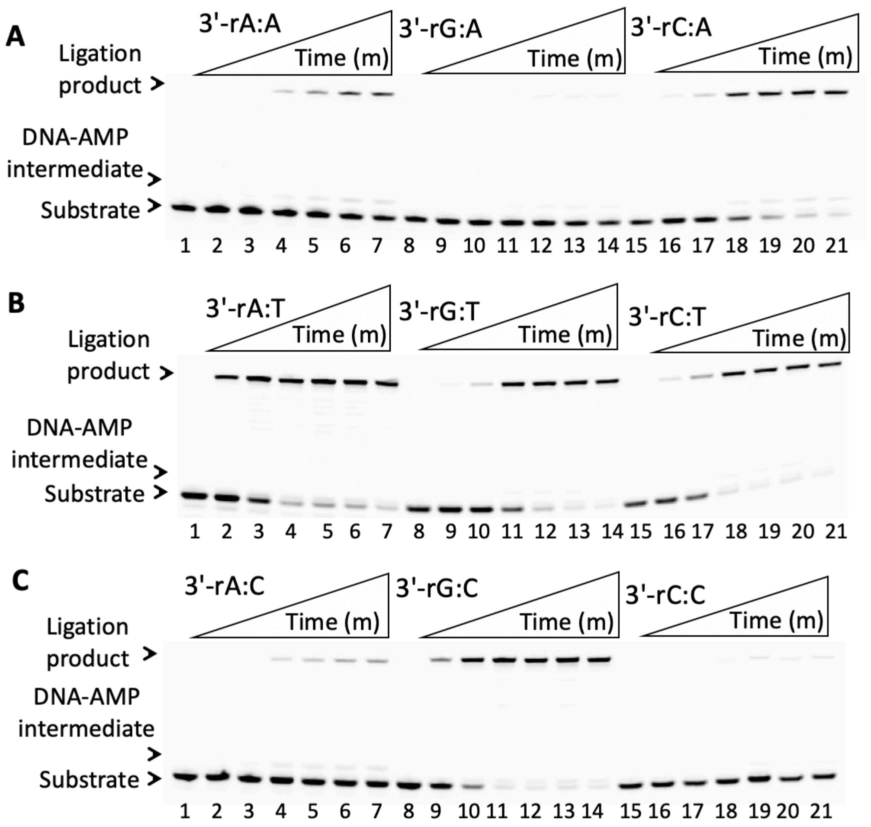


**Supplementary Figure 13. Ligation of nick DNA substrates containing 3'-ribonucleotide by LIG1 F635A. (A-C)** Lanes 1, 8, and 15 are the negative enzyme controls of the nick DNA substrates containing template A, T, and C mismatches. Lanes 2-7, 9-14, and 16-21 are the ligation products in the presence of nick DNA substrates with a single ribonucleotide 3'-end by LIG1 F635A, and correspond to time points of 0.5, 1, 3, 5, 8, and 10 min. Graphs showing show time-dependent changes in the amounts of ligation products are presented in Figure 12.

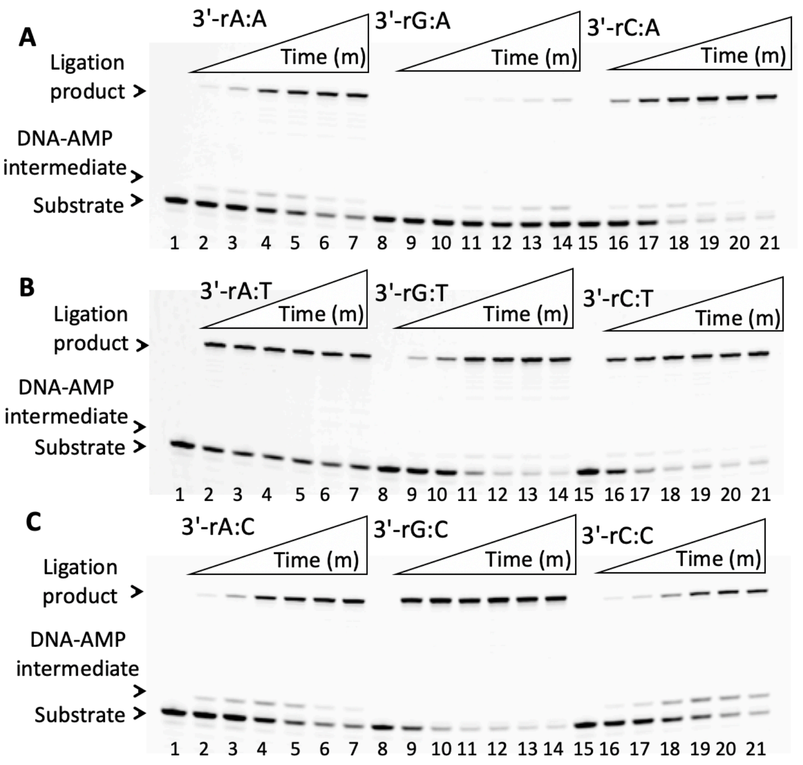


**Supplementary Figure 14. Ligation of nick DNA substrates containing 3'-ribonucleotide by LIG1 F635L. (A-C)** Lanes 1, 8, and 15 are the negative enzyme controls of the nick DNA substrates containing template A, T, and C mismatches. Lanes 2-7, 9-14, and 16-21 are the ligation products in the presence of nick DNA substrates with a single ribonucleotide 3'-end by LIG1 F635L, and correspond to time points of 0.5, 1, 3, 5, 8, and 10 min. Graphs showing show time-dependent changes in the amounts of ligation products are presented in Figure 13.

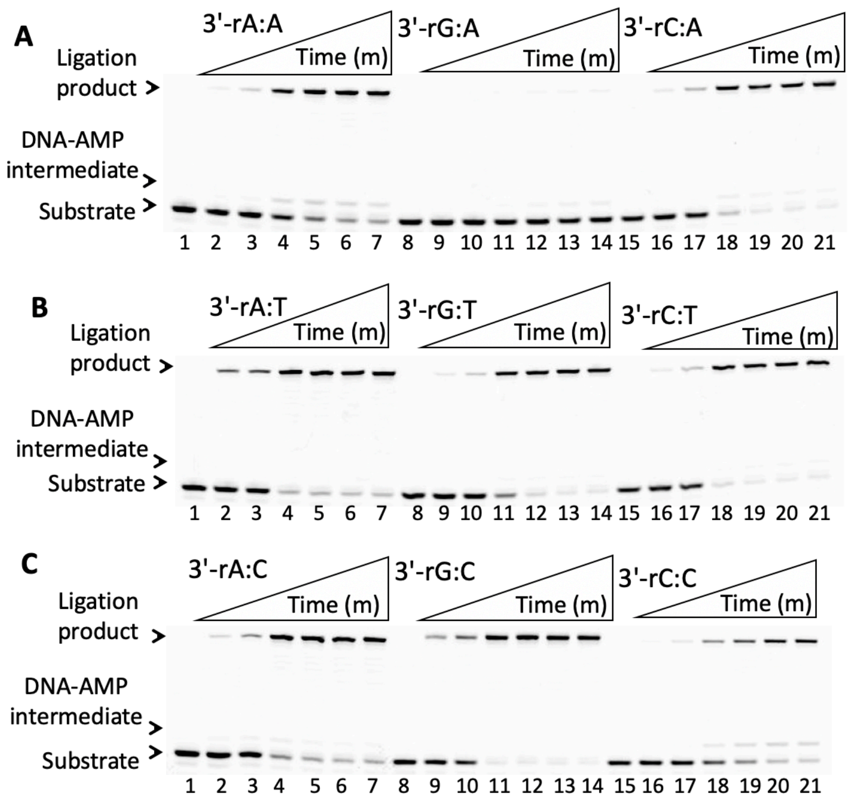


**Supplementary Figure 15. Ligation of nick DNA substrates containing 3'-ribonucleotide by LIG1 F872A. (A-C)** Lanes 1, 8, and 15 are the negative enzyme controls of the nick DNA substrates containing template A, T, and C mismatches. Lanes 2-7, 9-14, and 16-21 are the ligation products in the presence of nick DNA substrates with a single ribonucleotide 3'-end by LIG1 F872A, and correspond to time points of 0.5, 1, 3, 5, 8, and 10 min. Graphs showing show time-dependent changes in the amounts of ligation products are presented in Figure 14.

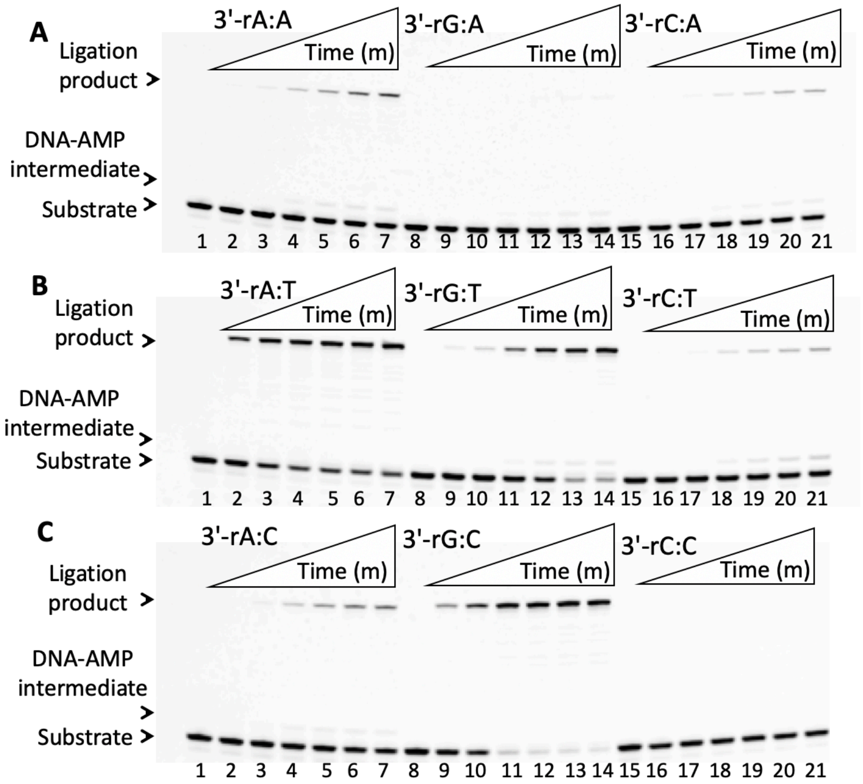


**Supplementary Figure 16. Ligation of nick DNA substrates containing 3'-ribonucleotide by LIG1 F872L. (A-C)** Lanes 1, 8, and 15 are the negative enzyme controls of the nick DNA substrates containing template A, T, and C mismatches. Lanes 2-7, 9-14, and 16-21 are the ligation products in the presence of nick DNA substrates with a single ribonucleotide 3'-end by LIG1 F872L, and correspond to time points of 0.5, 1, 3, 5, 8, and 10 min. Graphs showing show time-dependent changes in the amounts of ligation products are presented in Figure 15.

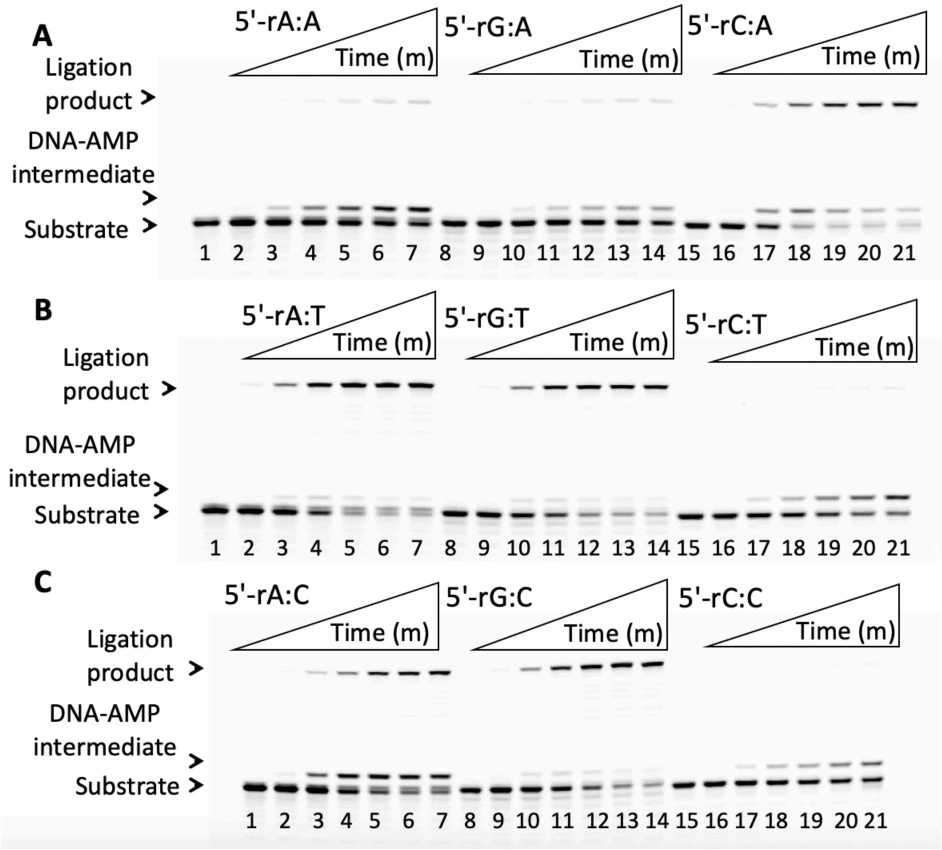


**Supplementary Figure 17. Ligation of nick DNA substrates containing 5'-ribonucleotide by LIG1 wild-type. (A-C)** Lanes 1, 8, and 15 are the negative enzyme controls of the nick DNA substrates containing template A, T, and C mismatches. Lanes 2-7, 9-14, and 16-21 are the ligation products in the presence of nick DNA substrates with a single ribonucleotide 5'-end by LIG1 wild-type, and correspond to time points of 1, 5, 15, 30, 45, and 60 min. Graphs showing time-dependent changes in the amounts of ligation products are presented in Figure 16.

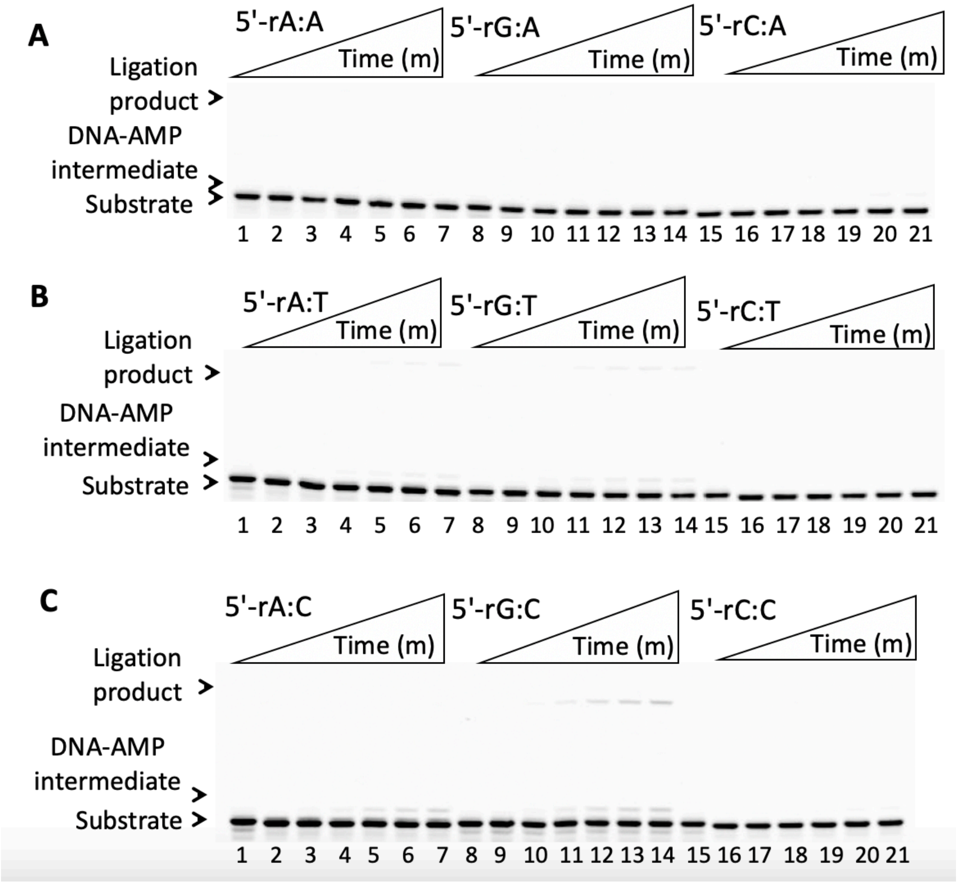


**Supplementary Figure 18. Ligation of nick DNA substrates containing 5'-ribonucleotide by LIG1 F635A. (A-C)** Lanes 1, 8, and 15 are the negative enzyme controls of the nick DNA substrates containing template A, T, and C mismatches. Lanes 2-7, 9-14, and 16-21 are the ligation products in the presence of nick DNA substrates with a single ribonucleotide 5'-end by LIG1 F635A, and correspond to time points of 1, 5, 15, 30, 45, and 60 min. Graphs showing time-dependent changes in the amounts of ligation products are presented in Figure 17.

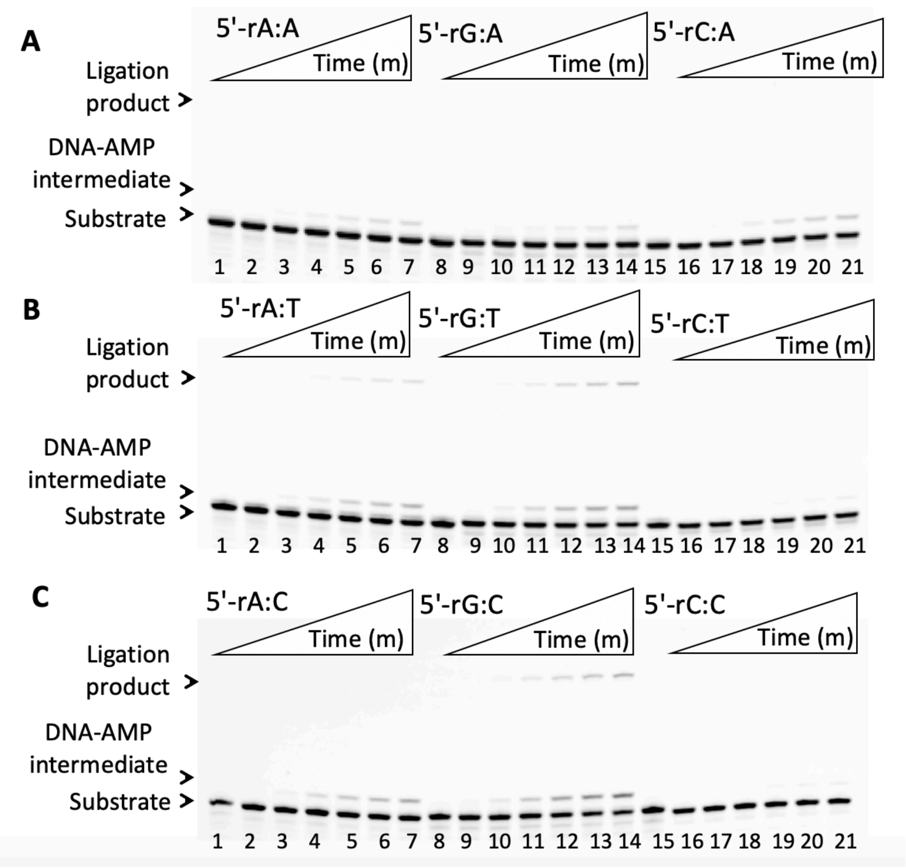


**Supplementary Figure 19. Ligation of nick DNA substrates containing 5'-ribonucleotide by LIG1 F635L. (A-C)** Lanes 1, 8, and 15 are the negative enzyme controls of the nick DNA substrates containing template A, T, and C mismatches. Lanes 2-7, 9-14, and 16-21 are the ligation products in the presence of nick DNA substrates with a single ribonucleotide 5'-end by LIG1 F635L, and correspond to time points of 1, 5, 15, 30, 45, and 60 min Graphs showing time-dependent changes in the amounts of ligation products are presented in Figure 18.

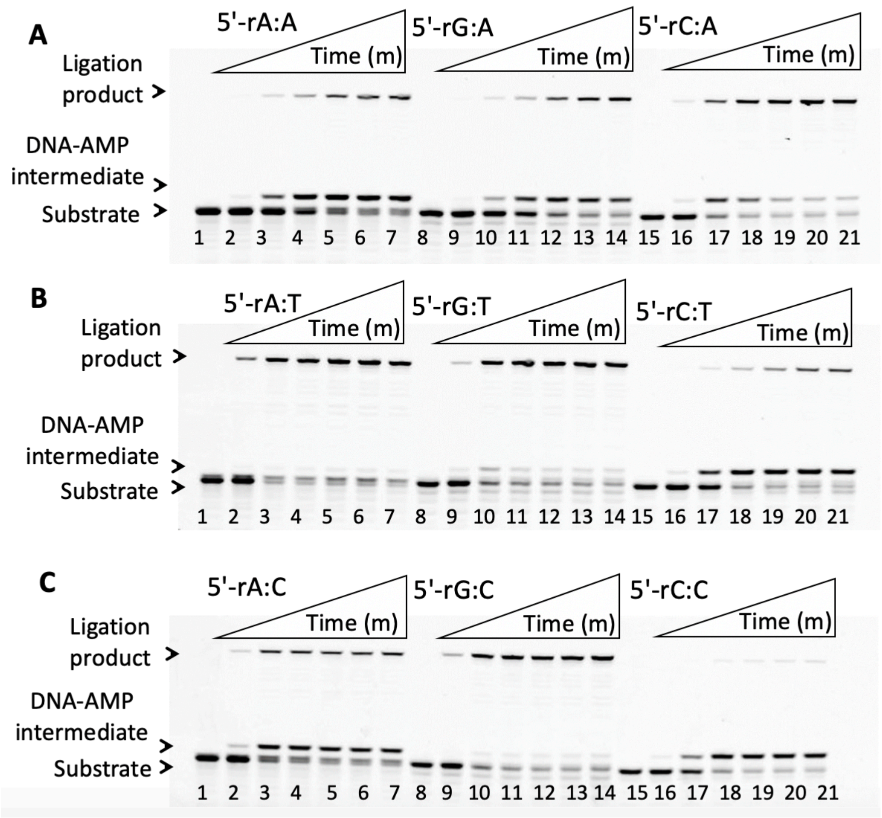


**Supplementary Figure 20. Ligation of nick DNA substrates containing 5'-ribonucleotide by LIG1 F872A. (A-C)** Lanes 1, 8, and 15 are the negative enzyme controls of the nick DNA substrates containing template A, T, and C mismatches. Lanes 2-7, 9-14, and 16-21 are the ligation products in the presence of nick DNA substrates with a single ribonucleotide 5'-end by LIG1 F872A, and correspond to time points of 1, 5, 15, 30, 45, and 60 min. Graphs showing time-dependent changes in the amounts of ligation products are presented in Figure 19.

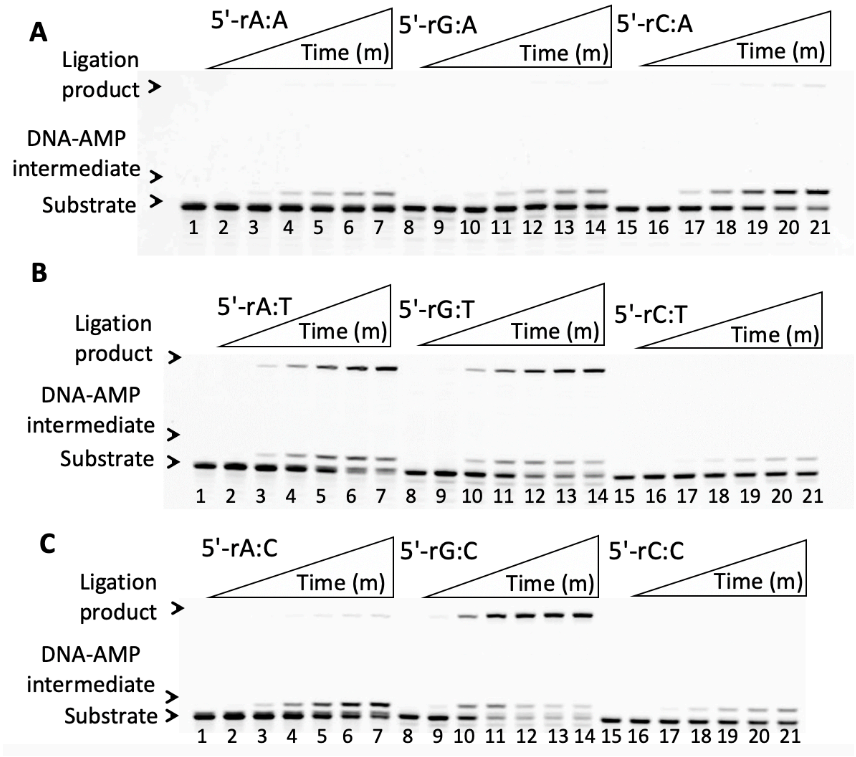


**Supplementary Figure 21. Ligation of nick DNA substrates containing 5'-ribonucleotide by LIG1 F872L. (A-C)** Lanes 1, 8, and 15 are the negative enzyme controls of the nick DNA substrates containing template A, T, and C mismatches. Lanes 2-7, 9-14, and 16-21 are the ligation products in the presence of nick DNA substrates with a single ribonucleotide 5'-end by LIG1 F872L, and correspond to time points of 1, 5, 15, 30, 45, and 60 min. Graphs showing time-dependent changes in the amounts of ligation products are presented in Figure 20.

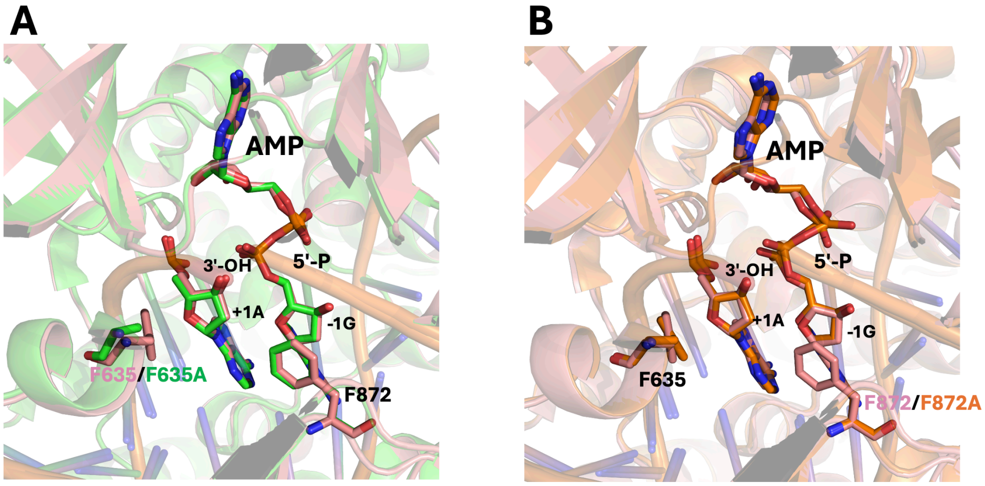


LIG1$^{EE/AA}$ /A:T (Step 2) vs
LIG1$^{EE/AA}$ F635A/A:T (Step 2)

LIG1$^{EE/AA}$ /A:T (Step 2) vs
LIG1$^{EE/AA}$ F872A/A:T (Step 2)

**Supplementary Figure 22. Structures of LIG1 active site mutants demonstrate a reliance on proper base pairing for end alignment at nick.** Superimposition of LIG1 structures demonstrate no significant difference in the position of the 3'- and 5'-ends of nick, and AMP across mutants in the presence of EE/AA mutation that refers to low-fidelity variant of LIG1.

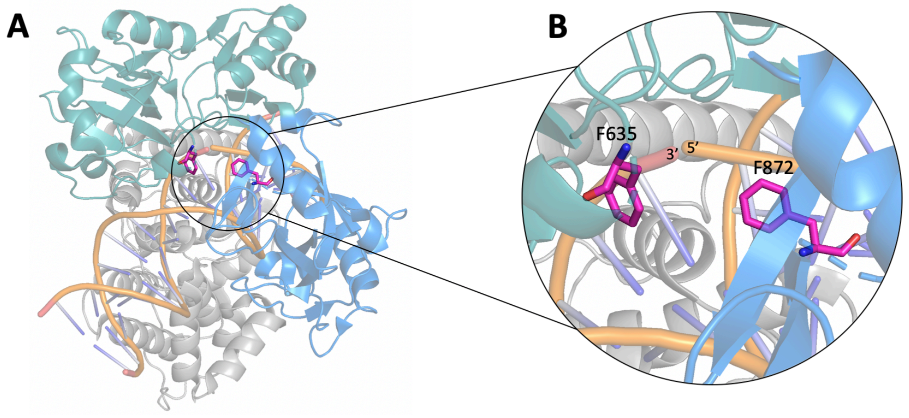


**C**

| DNA ligase | UniProt | n-5 | n-4 | n-3 | n-2 | n-1 | n | n+1 | n+2 | n+3 | n+4 | n+5 |
|---|---|---|---|---|---|---|---|---|---|---|---|---|
| DNA ligase I – *H. sapiens* | P18858 | K | Q | I | Q | P | F | Q | V | L | T | T |
| DNA ligase IIIα – *H. sapiens* | P49916 | G | K | P | L | P | F | G | T | L | G | V |
| DNA ligase I – *M. musculus* | P37913 | K | Q | I | Q | P | F | Q | V | L | T | T |
| DNA ligase I – *S. cerevisiae* | P04819 | G | K | I | L | P | F | Q | V | L | S | T |
| DNA ligase – *PBCV-1* | O41026 | F | S | Y | Y | W | F | D | Y | V | T | D |
| DNA ligase A – *E. coli* | P15042 | D | A | R | R | T | G | G | K | V | F | A |
| DNA ligase – *T4 bacteriophage* | P00970 | S | K | A | K | E | F | A | E | V | A | E |
| DNA ligase – *T7 bacteriophage* | P00969 | L | R | T | K | W | T | D | T | K | N | Q |

Green – exact alignment with LIG1 sequence
Yellow – same type of residue with LIG1 sequence (nonpolar, polar, positive charge, negative charge)
Red – no alignment with LIG1 sequence

**D**

| DNA ligase | UniProt | n-5 | n-4 | n-3 | n-2 | n-1 | n | n+1 | n+2 | n+3 | n+4 | n+5 |
|---|---|---|---|---|---|---|---|---|---|---|---|---|
| DNA ligase I – *H. sapiens* | P18858 | G | I | S | L | R | F | P | R | F | I | R |
| DNA ligase IIIα – *H. sapiens* | P49916 | G | I | S | I | R | F | P | R | C | T | R |
| DNA ligase I – *M. musculus* | P37913 | G | I | S | L | R | F | P | R | F | I | R |
| DNA ligase I – *S. cerevisiae* | P04819 | G | V | S | L | R | F | P | R | F | L | R |
| DNA ligase – *PBCV-1* | O41026 | K | D | C | P | R | F | P | V | F | I | G |
| DNA ligase A – *E. coli* | P15042 | T | T | F | A | R | F | L | Y | A | L | G |
| DNA ligase – *T4 bacteriophage* | P00970 | Y | V | K | L | F | L | P | I | A | I | R |
| DNA ligase – *T7 bacteriophage* | P00969 | D | G | S | L | R | H | P | S | F | V | M |

Green – exact alignment with LIG1 sequence
Yellow – same type of residue with LIG1 sequence (nonpolar, polar, positive charge, negative charge)
Red – no alignment with LIG1 sequence

**Supplementary Scheme 1. Positions of LIG1 F635 and F872 active site residues. (A)** Structure model of LIG1 encircling nick. F635 and F872 are colored purple. **(B)** Zoomed into the active site to highlight proximity of F635 and F872 to 3'- and 5'-end of nick, respectively. **(C-D)** The amino acid sequence alignments for the region corresponding to 629-640 and 867-877 residues of LIG1 demonstrates the conservation of the phenylalanine (F) residue at F635 and F872, respectively, where green represents an exact amino acid match, yellow represents an amino acid class match (polar, nonpolar, positively charged, or negatively charged), and red represents no match.

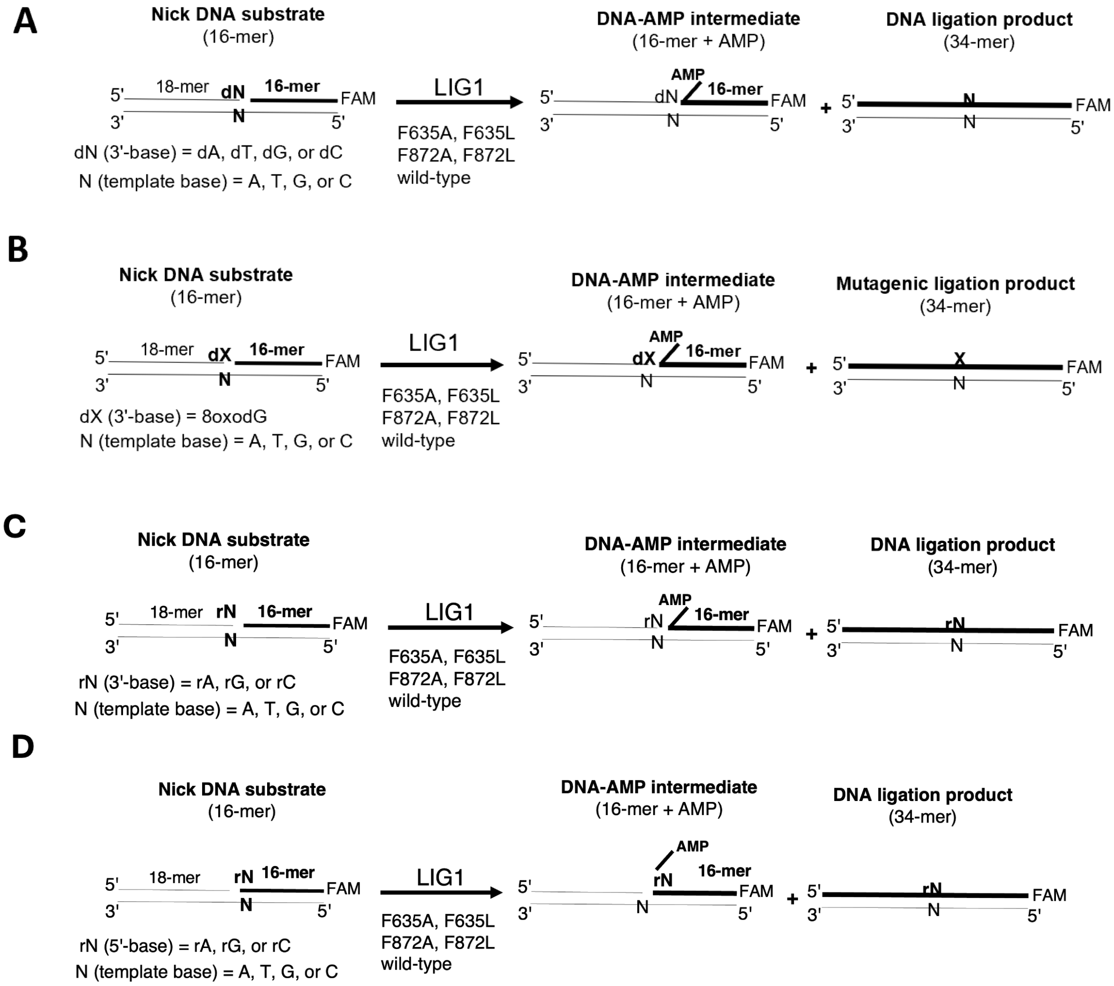


**Supplementary Scheme 2. Illustrations of DNA ligation assays used in this study. (A-D)** Ligation assays were used to evaluate the substrate specificity of LIG1 wild-type and active site mutants for the nick DNA substrates including 3'-mismatches (A), 3'-8oxodG (B), 3'-ribonucleotide (C), 5'-ribonucleotide (D). Reaction products include nick sealing and DNA-AMP intermediate with 5'-adenylate (AMP).

| Nick DNA Substrates | Sequence |
|---|---|
| 3'-dA:A | 5'-CATGGGCGGCATGAACCAGAGGCCCATCCTCACC-3'-FAM<br>3'-GTACCCGCCGTACTTGGACTCCGGGTAGGAGTGG-5' |
| 3'-dG:A | 5'-CATGGGCGGCATGAACCGGAGGCCCATCCTCACC-3'-FAM<br>3'-GTACCCGCCGTACTTGGACTCCGGGTAGGAGTGG-5' |
| 3'-dC:A | 5'-CATGGGCGGCATGAACCCGAGGCCCATCCTCACC-3'-FAM<br>3'-GTACCCGCCGTACTTGGACTCCGGGTAGGAGTGG-5' |
| 3'-dT:A | 5'-CATGGGCGGCATGAACCTGAGGCCCATCCTCACC-3'-FAM<br>3'-GTACCCGCCGTACTTGGACTCCGGGTAGGAGTGG-5' |
| 3'-dT:T | 5'-CATGGGCGGCATGAACCTGAGGCCCATCCTCACC-3'-FAM<br>3'-GTACCCGCCGTACTTGGTCTCCGGGTAGGAGTGG-5' |
| 3'-dG:T | 5'-CATGGGCGGCATGAACCGGAGGCCCATCCTCACC-3'-FAM<br>3'-GTACCCGCCGTACTTGGTCTCCGGGTAGGAGTGG-5' |
| 3'-dC:T | 5'-CATGGGCGGCATGAACCCGAGGCCCATCCTCACC-3'-FAM<br>3'-GTACCCGCCGTACTTGGTCTCCGGGTAGGAGTGG-5' |
| 3'-dA:T | 5'-CATGGGCGGCATGAACCAGAGGCCCATCCTCACC-3'-FAM<br>3'-GTACCCGCCGTACTTGGTCTCCGGGTAGGAGTGG-5' |
| 3'-dA:G | 5'-CATGGGCGGCATGAACCAGAGGCCCATCCTCACC-3'-FAM<br>3'-GTACCCGCCGTACTTGGGCTCCGGGTAGGAGTGG-5' |
| 3'-dT:G | 5'-CATGGGCGGCATGAACCTGAGGCCCATCCTCACC-3'-FAM<br>3'-GTACCCGCCGTACTTGGGCTCCGGGTAGGAGTGG-5' |
| 3'-dG:G | 5'-CATGGGCGGCATGAACCGGAGGCCCATCCTCACC-3'-FAM<br>3'-GTACCCGCCGTACTTGGGCTCCGGGTAGGAGTGG-5' |
| 3'-dC:G | 5'-CATGGGCGGCATGAACCCGAGGCCCATCCTCACC-3'-FAM<br>3'-GTACCCGCCGTACTTGGGCTCCGGGTAGGAGTGG-5' |
| 3'-dA:C | 5'-CATGGGCGGCATGAACCAGAGGCCCATCCTCACC-3'-FAM<br>3'-GTACCCGCCGTACTTGGCCTCCGGGTAGGAGTGG-5' |
| 3'-dT:C | 5'-CATGGGCGGCATGAACCTGAGGCCCATCCTCACC-3'-FAM<br>3'-GTACCCGCCGTACTTGGCCTCCGGGTAGGAGTGG-5' |
| 3'-dC:C | 5'-CATGGGCGGCATGAACCCGAGGCCCATCCTCACC-3'-FAM<br>3'-GTACCCGCCGTACTTGGCCTCCGGGTAGGAGTGG-5' |
| 3'-dG:C | 5'-CATGGGCGGCATGAACCGGAGGCCCATCCTCACC-3'-FAM<br>3'-GTACCCGCCGTACTTGGCCTCCGGGTAGGAGTGG-5' |

**Supplementary Table 1. Nick DNA substrates containing 3'-mismatches.** Nick DNA substrates with 3'-preinserted dA, dT, dG, dC opposite template base A, T, G, or C were used in the ligation assays to investigate the mismatch specificity of LIG1 wild-type and active site mutants. FAM denotes a fluorescent tag and is located at 3'-end of DNA substrates. The base at the template position is underlined.

| Nick DNA Substrates | Sequence |
|---|---|
| 3'-8oxodG:A | 5'-CATGGGCGGCATGAACC**X**GAGGCCCATCCTCACC-3'-FAM<br>3'-GTACCCGCCGTACTTGGACTCCGGGTAGGAGTGG-5' |
| 3'-8oxodG:C | 5'-CATGGGCGGCATGAACC**X**GAGGCCCATCCTCACC-3'-FAM<br>3'-GTACCCGCCGTACTTGGCCTCCGGGTAGGAGTGG-5' |
| 3'-8oxodG:G | 5'-CATGGGCGGCATGAACC**X**GAGGCCCATCCTCACC-3'-FAM<br>3'-GTACCCGCCGTACTTGGGCTCCGGGTAGGAGTGG-5' |
| 3'-8oxodG:T | 5'-CATGGGCGGCATGAACC**X**GAGGCCCATCCTCACC-3'-FAM<br>3'-GTACCCGCCGTACTTGGTCTCCGGGTAGGAGTGG-5' |

**Supplementary Table 2. Nick DNA substrates containing damaged ends.** Nick DNA substrates with 3'-preinserted 8-oxodG opposite template base A, T, G, or C were used in the ligation assays to investigate the ligation efficiency of LIG1 wild-type and active site mutants. FAM denotes a fluorescent tag and is located at 3'-end of DNA substrates. The base at the template position is underlined. The damaged base is shown in bold.

| Nick DNA Substrates | Sequence |
| --- | --- |
| 3'-rA:A | 5'-CATGGGCGGCATGAACC**A**GAGGCCCATCCTCACC-3'-FAM<br>3'-GTACCCGCCGTACTTGGACTCCGGGTAGGAGTGG-5' |
| 3'-rG:A | 5'-CATGGGCGGCATGAACC**G**GAGGCCCATCCTCACC-3'-FAM<br>3'-GTACCCGCCGTACTTGGACTCCGGGTAGGAGTGG-5' |
| 3'-rC:A | 5'-CATGGGCGGCATGAACC**C**GAGGCCCATCCTCACC-3'-FAM<br>3'-GTACCCGCCGTACTTGGACTCCGGGTAGGAGTGG-5' |
| 3'-rG:T | 5'-CATGGGCGGCATGAACC**G**GAGGCCCATCCTCACC-3'-FAM<br>3'-GTACCCGCCGTACTTGGTCTCCGGGTAGGAGTGG-5' |
| 3'-rC:T | 5'-CATGGGCGGCATGAACC**C**GAGGCCCATCCTCACC-3'-FAM<br>3'-GTACCCGCCGTACTTGGTCTCCGGGTAGGAGTGG-5' |
| 3'-rA:T | 5'-CATGGGCGGCATGAACC**A**GAGGCCCATCCTCACC-3'-FAM<br>3'-GTACCCGCCGTACTTGGTCTCCGGGTAGGAGTGG-5' |
| 3'-rA:G | 5'-CATGGGCGGCATGAACC**A**GAGGCCCATCCTCACC-3'-FAM<br>3'-GTACCCGCCGTACTTGGGCTCCGGGTAGGAGTGG-5' |
| 3'-rG:G | 5'-CATGGGCGGCATGAACC**G**GAGGCCCATCCTCACC-3'-FAM<br>3'-GTACCCGCCGTACTTGGGCTCCGGGTAGGAGTGG-5' |
| 3'-rC:G | 5'-CATGGGCGGCATGAACC**C**GAGGCCCATCCTCACC-3'-FAM<br>3'-GTACCCGCCGTACTTGGGCTCCGGGTAGGAGTGG-5' |
| 3'-rA:C | 5'-CATGGGCGGCATGAACC**A**GAGGCCCATCCTCACC-3'-FAM<br>3'-GTACCCGCCGTACTTGGCCTCCGGGTAGGAGTGG-5' |
| 3'-rC:C | 5'-CATGGGCGGCATGAACC**C**GAGGCCCATCCTCACC-3'-FAM<br>3'-GTACCCGCCGTACTTGGCCTCCGGGTAGGAGTGG-5' |
| 3'-rG:C | 5'-CATGGGCGGCATGAACC**G**GAGGCCCATCCTCACC-3'-FAM<br>3'-GTACCCGCCGTACTTGGCCTCCGGGTAGGAGTGG-5' |

**Supplementary Table 3. Nick DNA substrates containing 3'-ribonucleotides.** Nick DNA substrates with 3'-preinserted rA, rG, rC opposite template base A, T, G, or C were used in the ligation assays to investigate the sugar discrimination of LIG1 wild-type and active site mutants against nick DNA substrates congaining 3'-ribonucleotide. FAM denotes a fluorescent tag and is located at 3'-end of DNA substrates. The base at the template position is underlined. The ribonucleotide at 3'-end of nick is shown in bold.

| Nick DNA Substrates | Sequence |
|---|---|
| 5'-rA:A | 5'-CATGGGCGGCATGAACC**A**GAGGCCCATCCTCACC-3'-FAM<br>3'-GTACCCGCCGTACTTGGACTCCGGGTAGGAGTGG-5' |
| 5'-rG:A | 5'-CATGGGCGGCATGAACC**G**GAGGCCCATCCTCACC-3'-FAM<br>3'-GTACCCGCCGTACTTGGACTCCGGGTAGGAGTGG-5' |
| 5'-rC:A | 5'-CATGGGCGGCATGAACC**C**GAGGCCCATCCTCACC-3'-FAM<br>3'-GTACCCGCCGTACTTGGACTCCGGGTAGGAGTGG-5' |
| 5'-rG:T | 5'-CATGGGCGGCATGAACC**G**GAGGCCCATCCTCACC-3'-FAM<br>3'-GTACCCGCCGTACTTGGTCTCCGGGTAGGAGTGG-5' |
| 5'-rC:T | 5'-CATGGGCGGCATGAACC**C**GAGGCCCATCCTCACC-3'-FAM<br>3'-GTACCCGCCGTACTTGGTCTCCGGGTAGGAGTGG-5' |
| 5'-rA:T | 5'-CATGGGCGGCATGAACC**A**GAGGCCCATCCTCACC-3'-FAM<br>3'-GTACCCGCCGTACTTGGTCTCCGGGTAGGAGTGG-5' |
| 5'-rA:G | 5'-CATGGGCGGCATGAACC**A**GAGGCCCATCCTCACC-3'-FAM<br>3'-GTACCCGCCGTACTTGGGCTCCGGGTAGGAGTGG-5' |
| 5'-rG:G | 5'-CATGGGCGGCATGAACC**G**GAGGCCCATCCTCACC-3'-FAM<br>3'-GTACCCGCCGTACTTGGGCTCCGGGTAGGAGTGG-5' |
| 5'-rC:G | 5'-CATGGGCGGCATGAACC**C**GAGGCCCATCCTCACC-3'-FAM<br>3'-GTACCCGCCGTACTTGGGCTCCGGGTAGGAGTGG-5' |
| 5'-rA:C | 5'-CATGGGCGGCATGAACC**A**GAGGCCCATCCTCACC-3'-FAM<br>3'-GTACCCGCCGTACTTGGCCTCCGGGTAGGAGTGG-5' |
| 5'-rC:C | 5'-CATGGGCGGCATGAACC**C**GAGGCCCATCCTCACC-3'-FAM<br>3'-GTACCCGCCGTACTTGGCCTCCGGGTAGGAGTGG-5' |
| 5'-rG:C | 5'-CATGGGCGGCATGAACC**G**GAGGCCCATCCTCACC-3'-FAM<br>3'-GTACCCGCCGTACTTGGCCTCCGGGTAGGAGTGG-5' |

**Supplementary Table 4. Nick DNA substrates containing 5'-ribonucleotides.** Nick DNA substrates with 5'-preinserted rA, rG, rC opposite template base A, T, G, or C were used in the ligation assays to investigate the sugar discrimination of LIG1 wild-type and active site mutants against nick DNA substrates containing 5'-ribonucleotide. FAM denotes a fluorescent tag and is located at 3'-end of DNA substrates. The base at the template position is underlined. The ribonucleotide at 5'-end of nick is shown in bold.

| Oligonucleotide | Sequence (5'-3') |
|---|---|
| Template T | GTCCGAC<u>T</u>ACGCATCAGC |
| Upstream A | GCTGATGCGT**A** |
| Downstream (5'-P) | P-GTCGGAC |

**Supplementary Table 5. Oligonucleotides used in LIG1 crystallization.** Upstream oligonucleotide (3'-A), downstream oligonucleotide with phosphate (P) at the 5'-end, and template oligonucleotide containing T on a template position were used to prepare the nick DNA substrate with 3'-A:T for LIG1 crystallizations. The base at template base position is underlined and the base position at the 3'-end of nick is shown in bold.

| Oligonucleotide | Sequence |
|---|---|
| Up-OH | 5'-Bio-CATGGGCGGCATGAACC**A**-3' |
| Template T | 5'-GGTGAGGATGGGCCTC<u>T</u>GGTTCATGCCGCCCATG-3' |
| Down | 5'-(P)GAGGCCCATCCTCACC-AF488-3' |

**Supplementary Table 6. Nick DNA substrate used in the TIRF.** Up-OH, Template T, and down oligonucleotides were used to prepare the DNA substrate with a single nick site for single-molecule characterization of LIG1 nick DNA binding. Bio denotes a Biotin label located at 5'-end, AF488 is a green-fluorescent dye located at 3'-end, and P stands for a phosphate at 5'-end. The base at 3'-end is shown as bold and the template base is underlined.
